# Maternal experience alters brain-wide representation of infant cries

**DOI:** 10.64898/2026.06.26.734581

**Authors:** Briana R. McRae, Dianne-Lee K.D. Ferguson, Kalei Swanier, Genevera Allen, Bianca J. Marlin

## Abstract

**One Sentence:** Maternal experience fundamentally reorganizes infant cue processing across the entire brain into a more efficient and specialized caregiving state.

Pup calls signal the presence of a mouse infant in need of care (Bernd Haack et al., 1983; Ehret, 2005). Previous work demonstrated that experience-dependent plasticity within the left primary auditory cortex is oxytocin-facilitated and enhances neural and behavioral responses to these distress vocalizations (Marlin et al., 2015). However, whether maternal experience reshapes pup call processing beyond auditory regions remains unknown. Here, we show that both innate and learned maternal experience shape pup call-evoked behavior and brain-wide neural responses. Combining behavioral assays, whole-brain activity mapping, and oxytocinergic projection analyses, we find that pup calls recruit distinct large-scale neural networks in naïve virgins, experienced virgins, and mothers. With more maternal experience we observe a distributed pup call response network that is sparser yet more strongly coactivated, consistent with a more efficient and specialized neural representation of infant cues. We identified an oxytocin-dense circuit that shows an experience-dependent pup call response pattern. Therefore, we propose that experience-dependent refinement of the brain-wide response to pup calls is facilitated by oxytocin. Taken together, our findings reveal that maternal experience refines brain-wide sensory processing to support adaptive caregiving behavior.

## Introduction

Motherhood requires rapid adaptation to infant-derived sensory cues that guide caregiving behavior. Across mammals, infants communicate their needs through vocalizations, including crying in humans and ultrasonic vocalizations in house mice (*Mus musculus*). These vocal signals are among the most salient sensory cues encountered by caregivers and can powerfully shape maternal behavior.

Mouse pups emit ultrasonic isolation calls when separated from the nest (Ehret, 2005; Liu et al., 2003; Noirot, 1972). These vocalizations drive maternal behavior, guiding caregivers toward displaced offspring and eliciting pup retrieval (Ehret, 2005; Hernandez-Miranda et al., 2017a; Schiavo et al., 2020). Because auditory cues can signal the location of a pup from a distance, pup calls represent highly salient sensory signals for caregivers, even in the absence of pup cues of other sensory modalities (Bernd Haack et al., 1983; McRae et al., 2023). Consistent with this, playback of pup calls alone is sufficient to attract mothers toward a sound source, eliciting the phonotaxis response that marks the initiation of pup retrieval (Ehret, 1987; Schiavo et al., 2020; Sewell, 1970; Tasaka et al., 2020).

Mothers are experts at pup retrieval, even generalizing to retrieve pups that are not their own (Bendesky et al., 2017; Chantrey & Jenkins, 1982; Khadraoui et al., 2022). On the contrary, naïve virgin females typically do not retrieve pups reliably (Carcea et al., 2021; Ehret et al., 1987; Ehret & Koch, 1989; Koch & Ehret, 1989; Marlin et al., 2015; Schiavo et al., 2020).

However, following cohousing with a mother and litter for multiple days, virgin females can learn to perform pup retrieval expertly, reclassifying them as experienced virgins (Carcea et al., 2021; Dvorkin & Shea, 2022; Ehret et al., 1987; Koch & Ehret, 1989; Marlin et al., 2015; Nowlan et al., 2024; Rosenblatt, 1967; Schiavo et al., 2020). With this shift in pup-directed behavior following maternal experience comes a shift in the neural representation of pup calls, particularly in the left primary auditory cortex (A1). Both mothers and experienced virgins, but not naïve virgins, demonstrate pup call-evoked, time-locked neural activity in left A1 (Liu et al., 2006; Liu & Schreiner, 2007; Marlin et al., 2015; Rothschild et al., 2013). This experience-dependent neural plasticity in left A1 is often referred to as maternal plasticity, and it has been shown to be facilitated by the neuropeptide oxytocin (OXT). OXT acts on left A1 by rebalancing excitatory and inhibitory responses to pup calls, thus enhancing pup call-evoked spiking and enabling adaptive maternal care (Marlin et al., 2015; Schiavo et al., 2020).

It remains unknown whether, alongside this auditory maternal plasticity, other brain regions exhibit similar changes in their pup call responses, leaving our understanding of the maternal brain incomplete. While some studies have reported experience-dependent responses to pup calls in select higher-order auditory areas (Fichtel & Ehret, 1999; Tasaka et al., 2020), no study to date has surveyed whether such responses coincide with alternations throughout the entire mouse brain. Given the complexity of pup call-evoked behaviors that maternally experienced mice exhibit, we hypothesized that pup calls engage neural circuity beyond auditory areas, in a manner that reflects differences in maternal experience among female mice.

In this study, we utilized refined behavioral assays and whole-brain activity mapping via tissue clearing and c-Fos staining to evaluate how pup call behavioral and neural responses change with maternal experience. By examining naïve virgins, experienced virgins, and mothers, we sought to disentangle innate and learned aspects of maternal plasticity. Our findings revealed unexpected behavioral differences between experienced virgins and mothers, as well as differences in the coordinated activity of brain regions upon pup call presentation across all three groups that may serve to allow for more efficient maternal responses to infant distress cues in experienced animals.

### Pup retrieval and pup call preference change with maternal experience

To characterize how maternal experience arising from either innate or learned experience affects behavioral and neural responses to pup calls, we studied three groups: naïve virgin females (no maternal behavior), experienced virgin females (learned maternal behavior), and mothers (innate maternal behavior). Naïve virgins were female mice who have had no experience caring for pups. Experienced virgins were female mice who initially had no experience caring for pups, then were cohoused with a mother and her newborn litter for three to six days – a technique shown to initiate maternal behaviors in females via social learning (Carcea et al., 2021; Dvorkin & Shea, 2022; Ehret et al., 1987; Koch & Ehret, 1989; Marlin et al., 2015; Nowlan et al., 2024; Rosenblatt, 1967; Schiavo et al., 2020). Mothers were non-pregnant female mice who had successfully weaned two litters, with the second litter weaned one to four weeks prior to experiments.

To establish these experimental groups, each mouse underwent a pup retrieval assay (**Figure 1A**, **Supplemental Video 1**). To be considered naïve, virgins had to retrieve fewer than 10% of pups (thus screening out spontaneous retrievers); to be considered experienced, post-cohousing virgins and mothers had to retrieve at least 70% of pups (to only include experienced mice that retrieve pups reliably). In addition to screening mice prior to experiments, the pup retrieval assay also served to eliminate novelty as a potential confound in experimental results, as each mouse was exposed to pups and pup calls through this procedure. Our results showed that experienced virgins and mothers exhibited similar pup retrieval success (**Figure 1B**), and experienced virgins and mothers also exhibited indistinguishable pup retrieval latency (**Extended Figure 1**), highlighting the successful nature of pup-retrieval in the maternally experienced groups.

**Figure 1:**
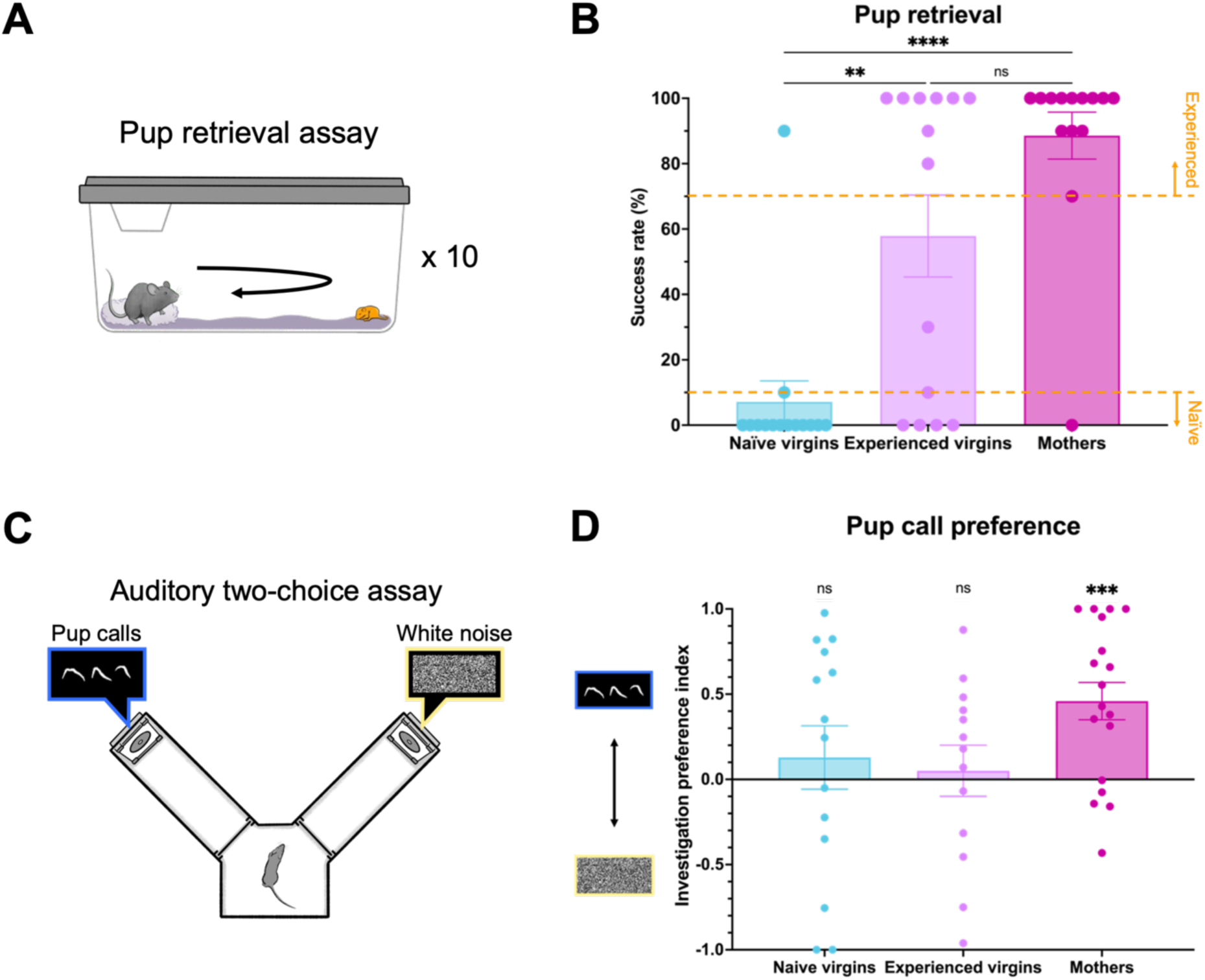
Mothers and experienced virgins retrieve pups successfully, but only mothers prefer pup calls. (A) Pup retrieval assay protocol, see Methods for details. (B) Pup retrieval success per mouse. Kruskal-Wallis test revealed significant group effect (p < 0.0001). Comparisons on graph represent Tukey’s post-hoc tests (naïve virgins vs. experienced virgins: p = 0.0097; naïve virgins vs. mothers: p < 0.0001; experienced virgins vs. mothers: p = 0.4431; n = 14 per group). Error bars denote mean ± SEM. **** p < 0.0001, ** p < 0.01. (C) Auditory 2-choice assay protocol (pup calls vs. white noise), see Methods for details. (D) Pup call investigation preference index per mouse during first 2 minutes of trial, see Methods for details. 1-sample t-tests (versus 0) revealed only mothers prefer pup calls (naïve virgins: p = 0.5028; experienced virgins: p = 0.7424; mothers: p = 0.0006; n = 14 naïve virgins, 13 experienced virgins, 18 mothers). 1-way ANOVA revealed no significant group effect (p = 0.2674; n = 14 naïve virgins, 13 experienced virgins, 18 mothers). Error bars denote mean ± SEM. Comparisons on graph represent 1-sample t-tests. *** p < 0.001.

Pup calls are crucial for eliciting the act of pup retrieval (Hernandez-Miranda et al., 2017b; Schiavo et al., 2020). Prior work has shown that the presentation of pup calls is sufficient to attract mothers, but not naïve virgins, to a speaker (Ehret, 1987; Ehret et al., 1987; Schiavo et al., 2020; Sewell, 1970; Tasaka et al., 2020). This maternal attraction towards pup calls has been demonstrated in various studies through two-choice assays, in which competing speakers in an arena play different auditory stimuli at the same time, and the behavior of a mouse in that arena is quantified. One limitation of prior studies that used two-choice assays to assess pup call preference, however, is that pup odors were introduced into the arena prior to testing via pup retrieval or the presence of pups in the nest. This presents a confound, as prior research has shown that the presentation of pup odors amplifies A1 neural responses to pup calls in experienced virgins and mothers (Cohen et al., 2011; Cohen & Mizrahi, 2015).

To investigate preference towards pup calls in the absence of other multisensory pup cues, we recorded pup calls from female and male six-day-old pups. We then confirmed that the temporal and spectral features of these pup calls aligned with those reported in prior literature (Fonseca et al., 2021; Schiavo et al., 2020), and created auditory stimulus files (**Extended Figure 2**). We then designed an auditory two-choice assay that explicitly did not involve the placement of pups nor bedding in the arena at any point, eliminating pup olfactory cues (**Figure 1C**, **Supplemental Videos 2-4**). Given two minutes to freely explore a custom-built two-choice arena while competing auditory stimuli played, mothers had a significant preference towards interacting with a speaker playing pup calls, compared to a speaker playing white noise, while naïve virgins did not. Surprisingly, however, experienced virgins also showed no significant preference to pup calls, suggesting that while both mothers and experienced virgins retrieve pups, the neural circuitry supporting this behavior may differ (**Figure 1D**). These results held true whether preference was calculated based on time spent investigating the speakers (**Figure 1D**) or on the number of times the speaker investigation zones were entered (**Extended Figure 3A**), as prior studies have analyzed (Ehret, 1987; Ehret et al., 1987; Lin et al., 2013a; Schiavo et al., 2020; Sewell, 1970; Tasaka et al., 2020). If the assay was extended to 10 minutes, no group had a sustained preference (**Extended Figure 3B**). Preference during the first two minutes could not be explained by group differences in distance traveled (**Extended Figure 3C**), though it should be noted that naïve virgins froze significantly more than mothers, perhaps indicating that mothers were more active in their investigation behavior (**Extended Figure 3D**).

### Maternal experience alters the brain-wide representation of pup calls

Based on our findings from the pup retrieval and auditory two-choice assays, as well as prior work, we hypothesized that pup calls engage different neural circuitry in females with different maternal experience. We suspected that by investigating pup call responses beyond auditory areas, we could begin to better explain the unexpected difference in pup call preference we observed between experienced virgins and mothers. To do so, we measured pup call-evoked activity across the brain by using intact tissue clearing and activity mapping via c-Fos, a protein marker for neuronal activity.

To characterize brain-wide c-Fos expression following pup call presentation, we designed a sound exposure assay (**Figure 2A**). As described earlier, mice belonging to either the naïve virgin, experienced virgin, or mother group were established via pup retrieval and cohousing (**Extended Figure 4A**). For the purposes of this experiment, pup retrieval of cohoused virgins was tested each day to monitor any major differences in learning over time, of which we found none (**Extended Figure 4B**). Then, for the sound exposure assay, mice were exposed for 30 min to either the presentation of pup calls or a baseline condition (no stimulus presented, with the same ambient noise as the pup calls condition). 15–30 min later, their brains were perfused, cleared and stained for c-Fos via the iDISCO+ protocol (Renier et al., 2014), imaged with a light-sheet microscope, and reconstructed in 3D (**Figure 2B**, **Supplemental Videos 5-6**). We then used the ClearMap pipeline to detect c-Fos+ cells, identify their spatial coordinates within the sample, and align the sample to the Allen Brain Atlas to assign each c-Fos+ cell to a brain region (**Figure 2B**, **Supplemental Video 7**) (Renier et al., 2016). We simplified the Allen Brain Atlas into 215 regions that still tiled the whole brain but limited the granularity of parcellations (**Extended Figure 5**, **Table 1**). We also summed c-Fos+ cell counts per brain region bilaterally. Before doing so, however, we first examined whether there was any lateralization in c-Fos expression. As expected based on prior literature (Ehret, 1987; Levy et al., 2019; Marlin et al., 2015), we observed higher c-Fos expression in the left A1 of pup call-exposed samples; surprisingly, however, c-Fos expression in baseline samples was also left lateralized in A1, across all groups (**Extended Figure 6A**). It should be noted that hemisphere imaging order affected c-Fos+ cell counts due to bleaching near the midline after the first imaging session, yet still left-lateralization in A1 prevailed. However, aside from the effect of hemisphere imaging order, we did not see the same lateralization across any conditions when comparing across the whole brain, justifying the bilateral summation of c-Fos+ cell counts within each brain region (**Extended Figure 6B**).

**Figure 2:**
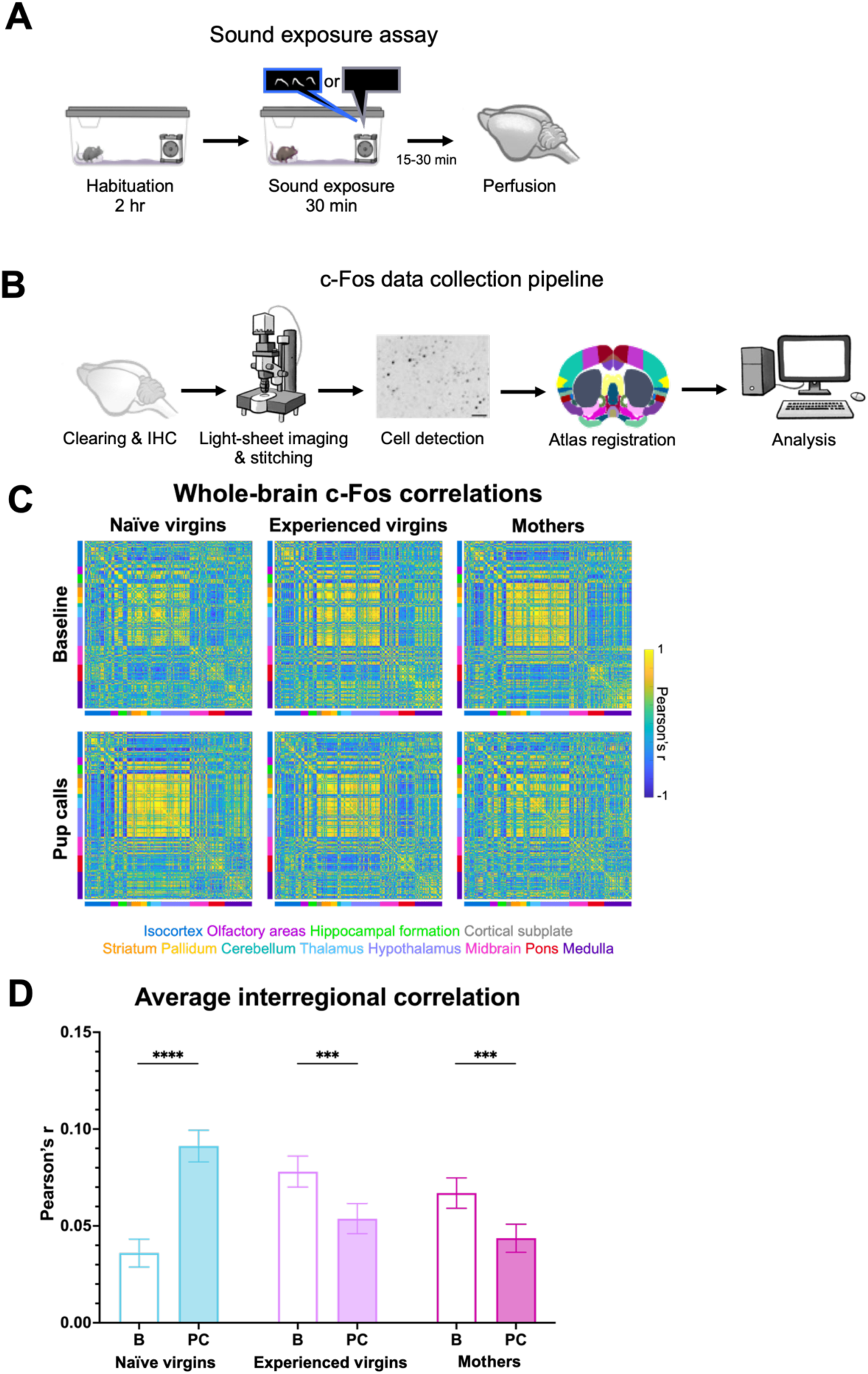
Pup calls exert an experience-dependent effect on brain-wide synchrony. (A) Sound exposure assay protocol, in which mice are presented with either pup calls or baseline (no stimulus), see Methods for details. (B) c-Fos data collection pipeline, which included iDISCO+ clearing, anti-c-Fos immunohistochemistry (IHC), light-sheet imaging and tile stitching, c-Fos+ cell detection via ClearMap, Allen Brain Atlas registration, and analysis (see Methods for details). Scale bar in representative image represents 25µm. (C) Pearson correlation matrices indicating interregional correlations, based on normalized c-Fos expression in baseline samples (top) and pup call-exposed samples (bottom) across groups (n = 8 x 215 regions). Color-blocked bars on axes correspond to broader brain areas according to the Allen Brain Atlas, see text below matrices for color coding. Matrix pixel colors represent Pearson correlation coefficients, see scale to the right. (D) Mean interregional correlation per condition (B= baseline, PC = pup calls). 2-way ANOVA following Fisher r-to-z transformation revealed significant effects of stimulus (p = 0.02) and of group and stimulus interaction (p < 0.0001). Comparisons on graph represent Tukey’s post-hoc tests of within-group baseline vs. pup calls (naïve virgins: p < 0.0001; experienced virgins: p = 0.0001796; mothers: p = 0.0004070; n = 8 x 215 regions). See Table 3 for other relevant pairwise comparisons. Error bars denote mean ± 95% CI. *** p < 0.001. **** p < 0.0001.

**Table 1:**
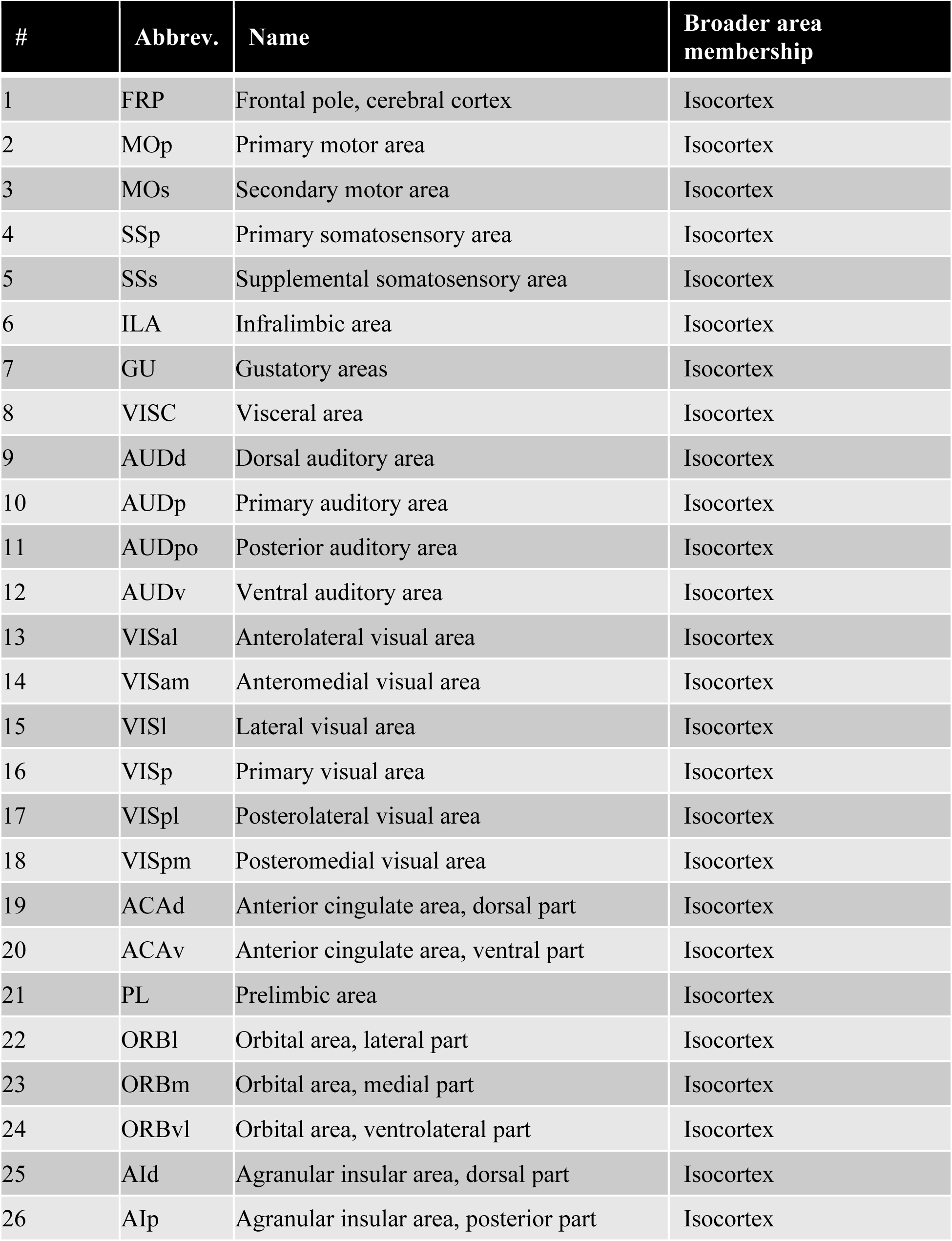
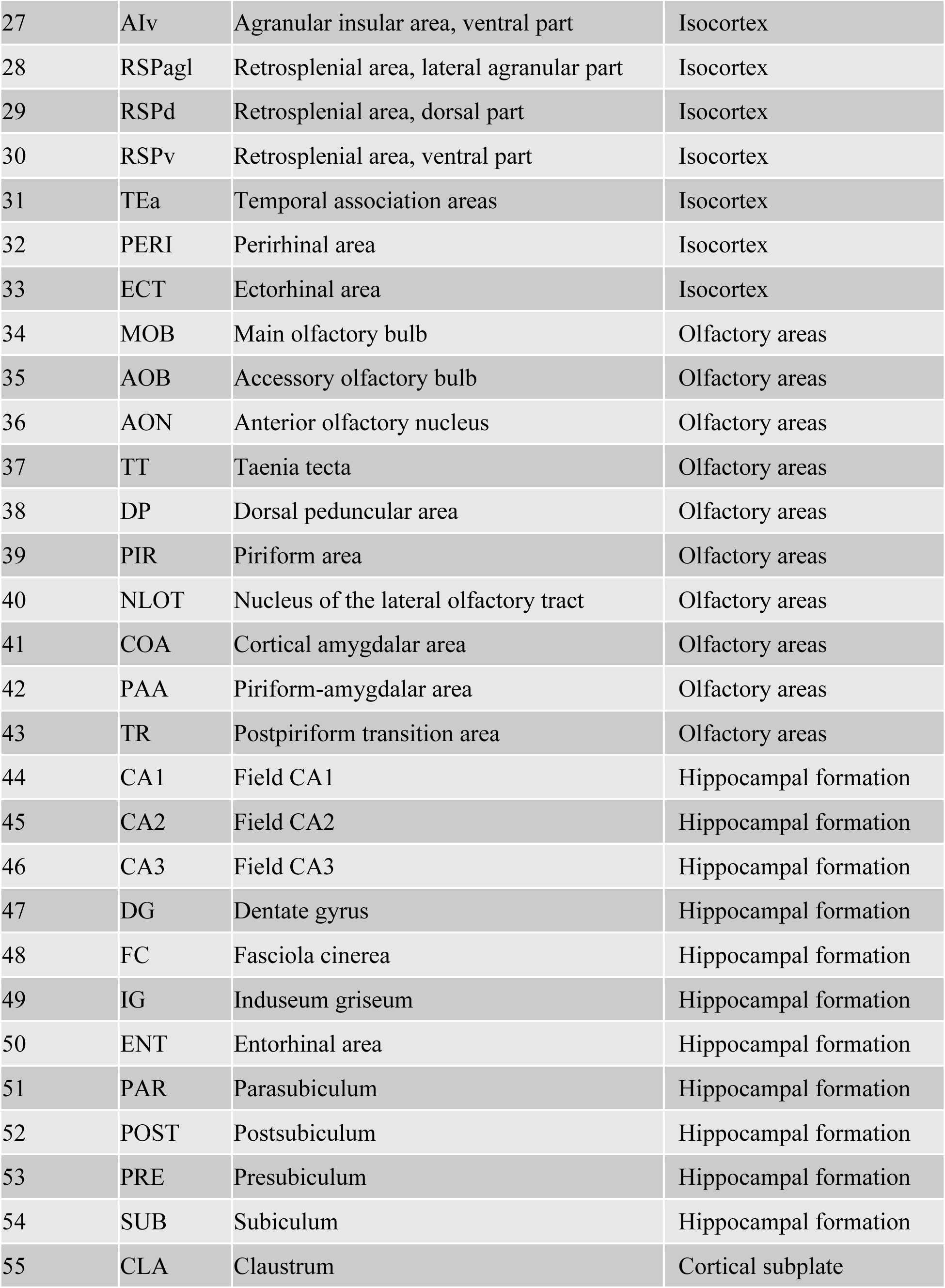
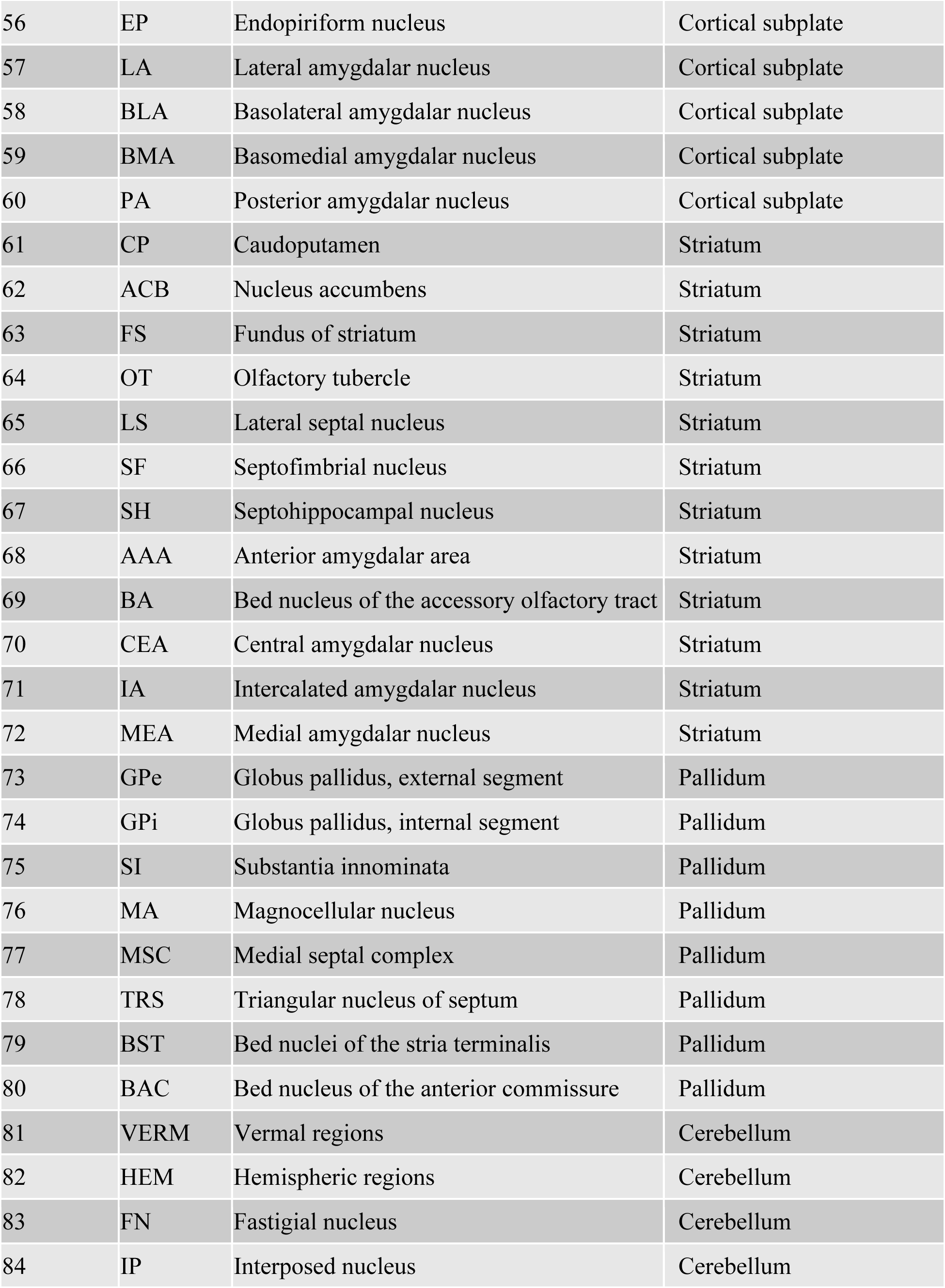
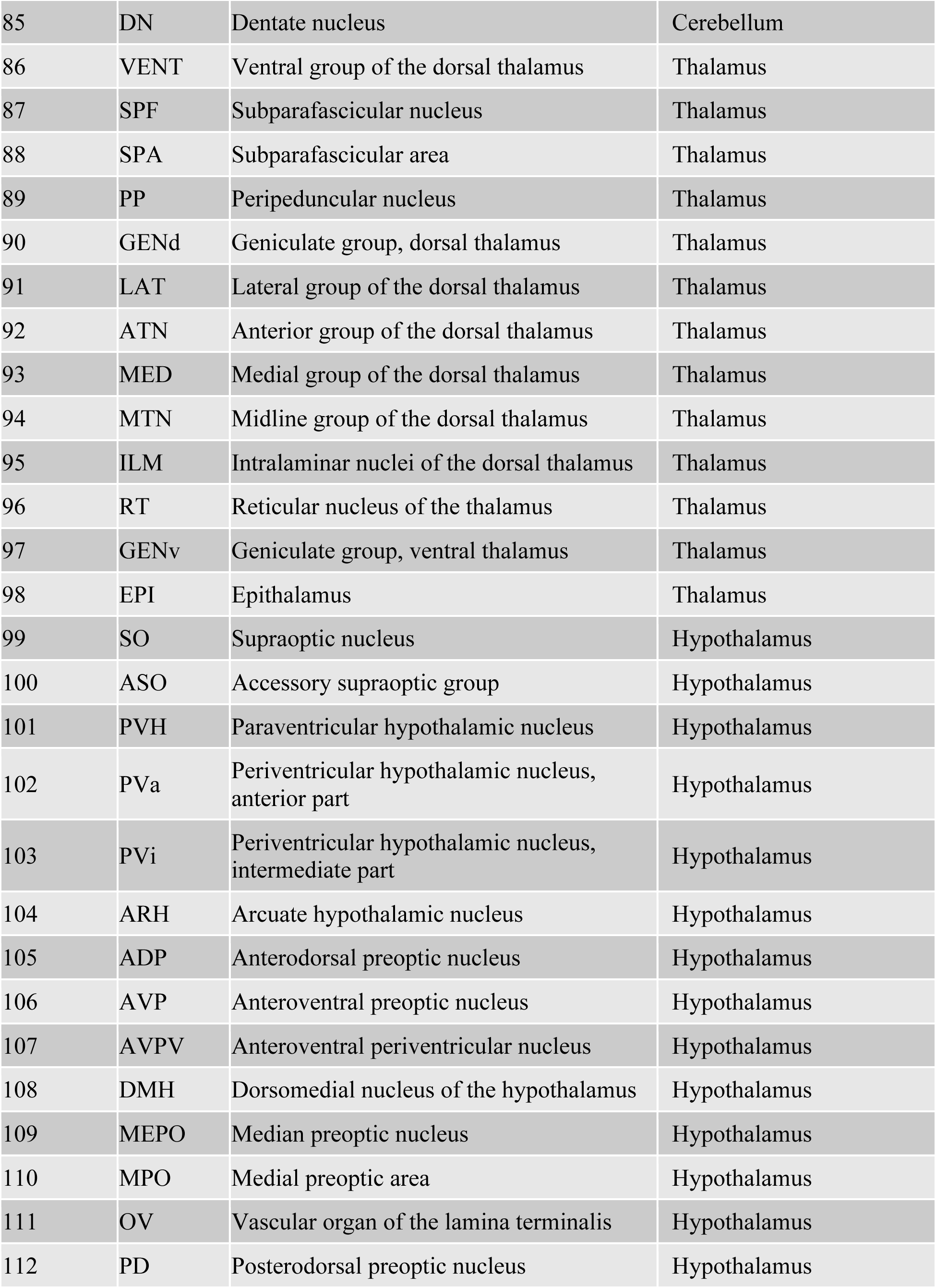
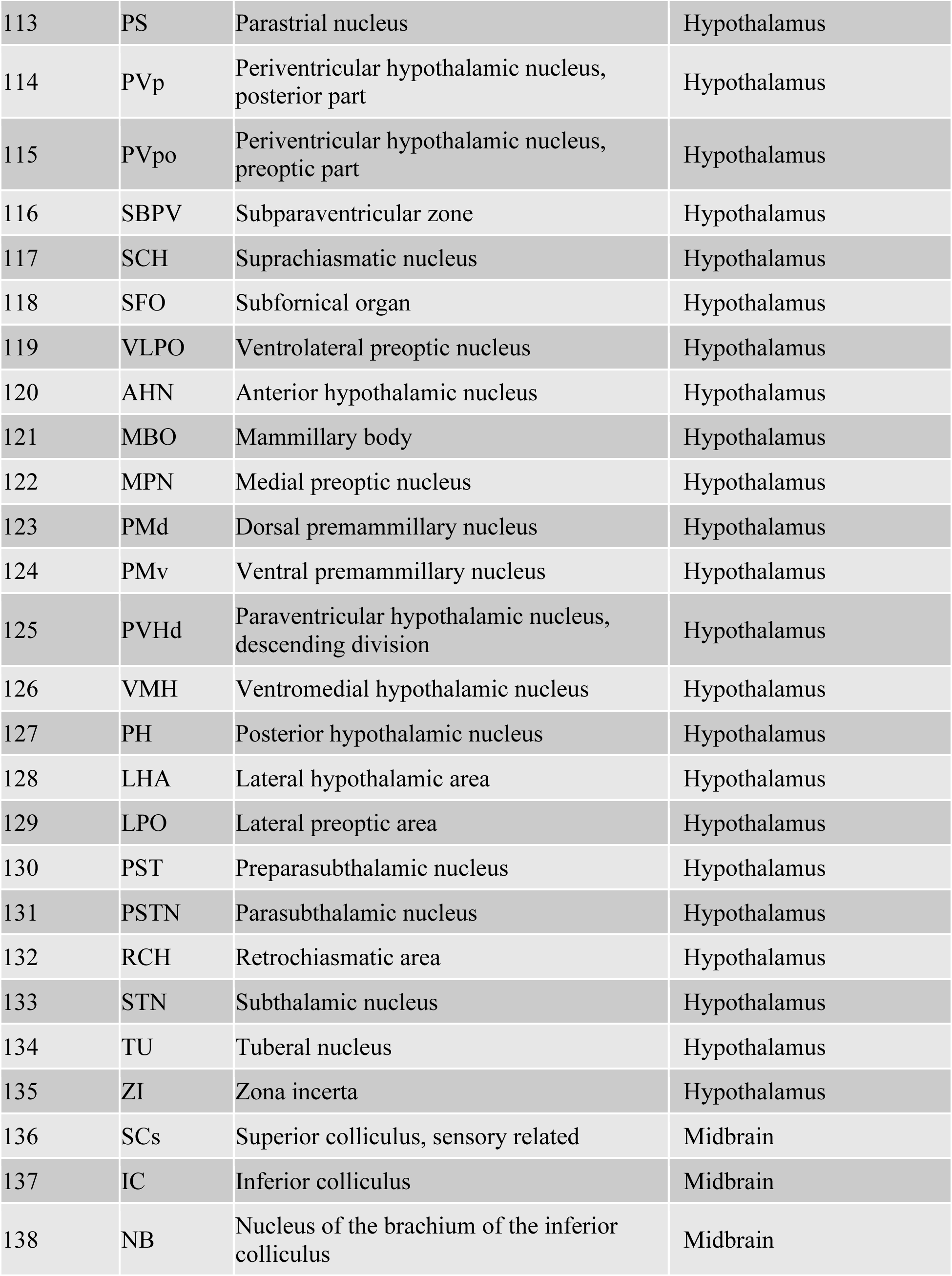
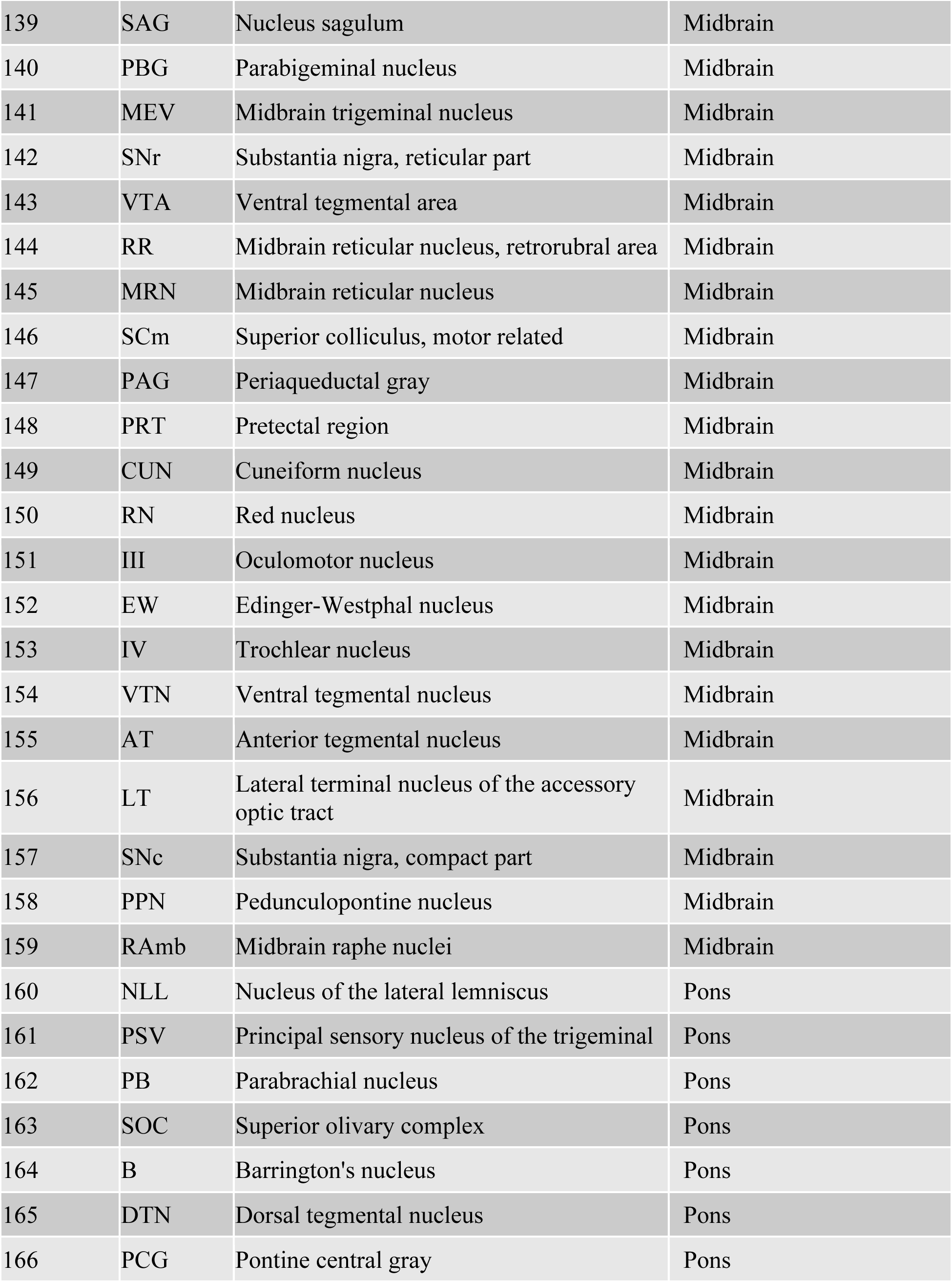
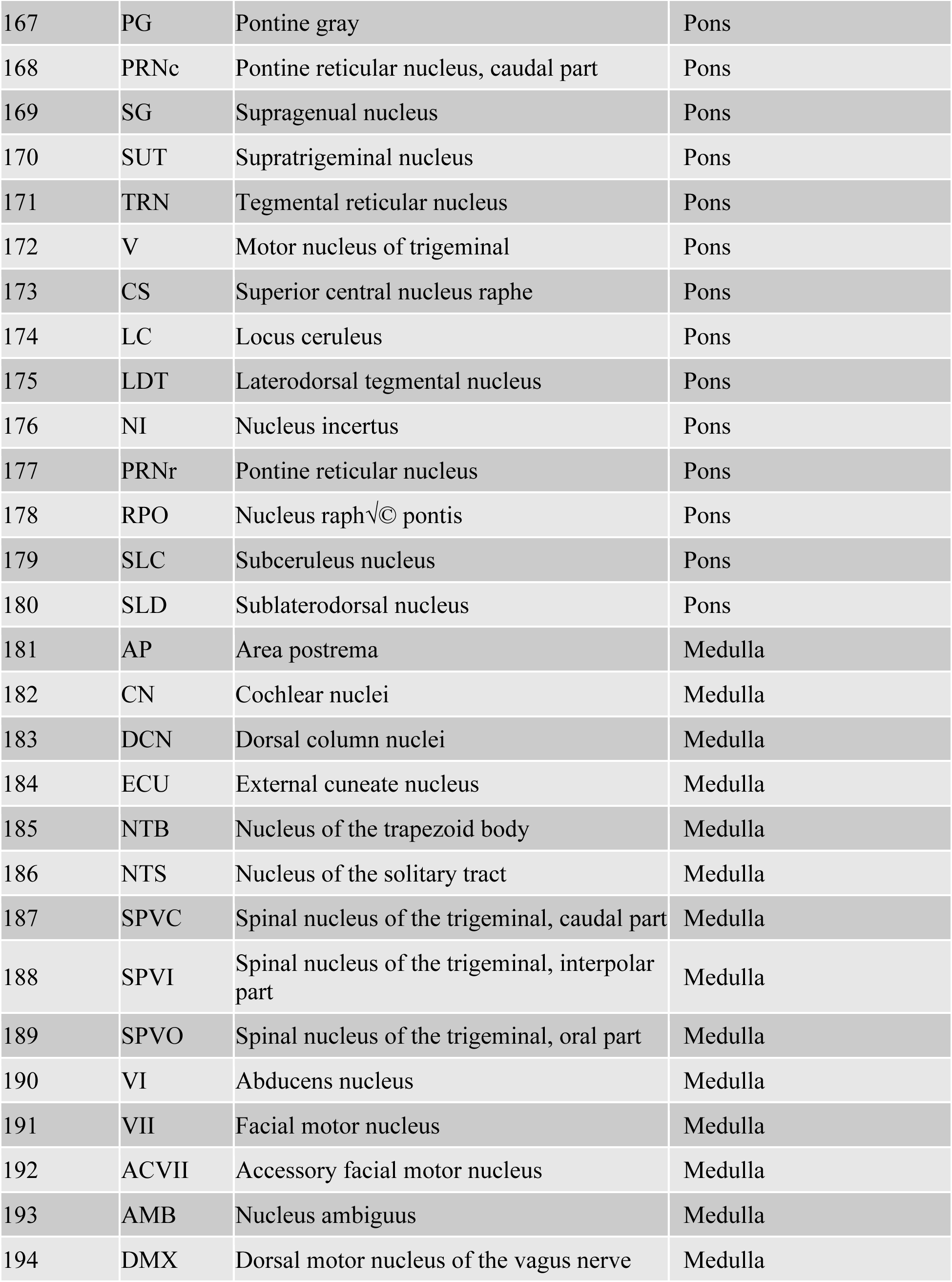
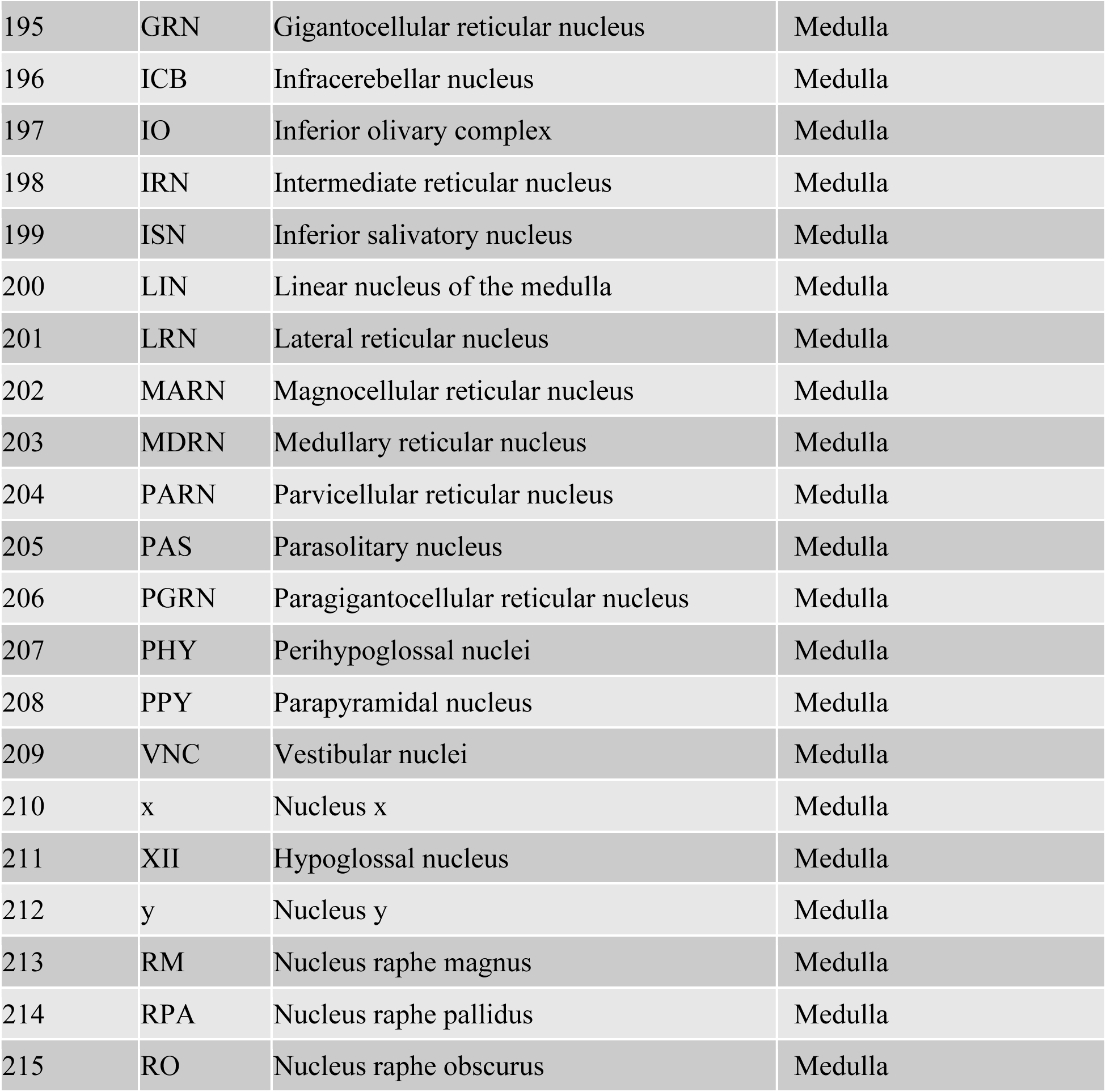
Allen Brain Atlas regions and abbreviations. For each of the 215 brain regions conserved from Allen Brain Atlas, this table lists their abbreviation, full name, and broader brain area membership.

To assess any potential sources of movement confounds in our c-Fos data, we analyzed the behavior of mice during the sound exposure assay. We tracked the mice with DeepLabCut, a deep learning-based pose estimation pipeline (Mathis et al., 2018). We used tracking of the nose to assess distance traveled and time spent in the defined speaker investigation zone during sound exposure (**Extended Figure 4C**). Our behavioral analysis results revealed no significant group effects on distance traveled nor speaker investigation time during the 30-minute trial, indicating that no correction for movement was required for group-level c-Fos expression analyses (**Extended Figure 4D-E**). However, there was a significant effect of stimulus (pup calls versus baseline) on distance traveled during the 30-minute trial (**Extended Figure 4D**), as well as on speaker investigation time when analysis was restricted to the first two minutes of the trial (**Extended Figure 4F**), both indicating successful stimulus delivery and perception.

Inspired by previous studies that analyzed similar datasets, we modeled c-Fos+ cell counts per region using a generalized linear mixed model (GLMM; see **Methods**) (Zimmerman et al., 2024). This analysis yielded 10 regions for which the inclusion of group and stimulus information provided better explanatory power, but only two regions passed correction for multiple comparisons (**Table 2**). It should be noted that compared to prior work using GLMMs to model brain-wide c-Fos, we were poorly powered, perhaps limiting the conclusions that can be drawn from this analysis regarding differences in regional c-Fos expression amplitude. We also observed a strong batch effect, which is a technical challenge for whole-brain immunohistochemistry. To address batch effects, we normalized c-Fos+ cell counts per region by total counts per sample, and we ensured that all conditions (all six different combinations of group and stimulus type) were equally represented per batch. Additionally, close examination of the spatial distribution of brain regions with significant batch effects after normalization revealed no apparent spatial pattern across the brain (**Extended Figure 7**). Given that our batches were balanced and batch effects were spread seemingly randomly throughout the brain, we moved forward with an analysis method that is robust to batch effect: correlation of normalized regional c-Fos expression across samples within each condition (Jin et al., 2025; Rogers et al., 2026; Wheeler et al., 2013).

**Table 2:** GLMM results. Results from fitting a full and reduced GLMM (see Methods) per brain region, and comparing the fit of the 2 models. Significant p-value would indicate that full GLMM explains the c-Fos+ counts in that brain region better than the reduced GLMM. Middle column indicates p-value of model comparison (green shading highlights regions with p < 0.05), rightmost column indicates adjusted p-value of model comparison (red shading highlights regions with FDR < 0.10).

| Region | p-value from model comparison | Adjusted p-value (FDR < 0.10) |
| --- | --- | --- |
| FRP | 0.86287857 | 0.92568707 |
| MOp | 0.07578609 | 0.68099235 |
| MOs | 0.77321023 | 0.8883698 |
| SSp | 0.37586341 | 0.79443153 |
| SSs | 0.70608113 | 0.8883698 |
| ILA | 0.47035576 | 0.88106721 |
| GU | 0.02492231 | 0.68099235 |
| VISC | 0.54794771 | 0.8883698 |
| AUDd | 0.28344322 | 0.73477471 |
| AUDp | 0.18566147 | 0.68099235 |
| AUDpo | 0.01198419 | 0.68099235 |
| AUDv | 0.8677685 | 0.92568707 |
| VISal | 0.13984382 | 0.68099235 |
| VISam | 0.62675302 | 0.8883698 |
| VISl | 8.81549290463625e-06 | 0.00189533 |
| VISp | 0.57569667 | 0.8883698 |
| VISpl | 0.00012337 | 0.01326226 |
| VISpm | 0.77733329 | 0.8883698 |
| ACAd | 0.32909511 | 0.76754788 |
| ACAv | 0.51974454 | 0.88686569 |
| PL | 0.65680196 | 0.8883698 |
| ORBl | 0.64396994 | 0.8883698 |
| ORBm | 0.19784725 | 0.68099235 |
| ORBvl | 0.24070838 | 0.72049635 |
| AId | 0.5657859 | 0.8883698 |
| AIp | 0.09555557 | 0.68099235 |
| AIv | 0.87402082 | 0.92568707 |
| RSPagl | 0.74845858 | 0.8883698 |
| RSPd | 0.5869099 | 0.8883698 |
| RSPv | 0.12006543 | 0.68099235 |
| TEa | 0.05687867 | 0.68099235 |
| PERI | 0.23879685 | 0.72049635 |
| ECT | 0.35773909 | 0.79292686 |
| MOB | 0.8373986 | 0.92425134 |
| AOB | 0.54683733 | 0.8883698 |
| AON | 0.74570139 | 0.8883698 |
| TT | 0.72137967 | 0.8883698 |
| DP | 0.57927024 | 0.8883698 |
| PIR | 0.2786311 | 0.73477471 |
| NLOT | 0.75866684 | 0.8883698 |
| COA | 0.1459944 | 0.68099235 |
| PAA | 0.89317323 | 0.927692 |
| TR | 0.66819247 | 0.8883698 |
| CA1 | 0.7509442 | 0.8883698 |
| CA2 | 0.92710861 | 0.94918263 |
| CA3 | 0.54114115 | 0.8883698 |
| DG | 0.75333551 | 0.8883698 |
| FC | 0.03812106 | 0.68099235 |
| IG | 0.31225872 | 0.75031315 |
| ENT | 0.18954748 | 0.68099235 |
| PAR | 0.78093903 | 0.8883698 |
| POST | 0.18383133 | 0.68099235 |
| PRE | 0.36906772 | 0.7934956 |
| SUB | 0.19659414 | 0.68099235 |
| CLA | 0.48196758 | 0.88106721 |
| EP | 0.87945236 | 0.92569342 |
| LA | 0.44771134 | 0.88106721 |
| BLA | 0.1899192 | 0.68099235 |
| BMA | 0.24965274 | 0.72049635 |
| PA | 0.09762295 | 0.68099235 |
| CP | 0.57714496 | 0.8883698 |
| ACB | 0.94685232 | 0.95574295 |
| FS | 0.61834115 | 0.8883698 |
| OT | 0.59648883 | 0.8883698 |
| LS | 0.70839075 | 0.8883698 |
| SF | 0.944892 | 0.95574295 |
| SH | 0.71849869 | 0.8883698 |
| AAA | 0.67215393 | 0.8883698 |
| BA | 0.99317296 | 0.99317296 |
| CEA | 0.6491498 | 0.8883698 |
| IA | 0.72629283 | 0.8883698 |
| MEA | 0.69154274 | 0.8883698 |
| GPe | 0.15105994 | 0.68099235 |
| GPi | 0.48636662 | 0.88106721 |
| SI | 0.3768931 | 0.79443153 |
| MA | 0.49201751 | 0.88106721 |
| MSC | 0.15335546 | 0.68099235 |
| TRS | 0.61040589 | 0.8883698 |
| BST | 0.77869919 | 0.8883698 |
| BAC | 0.07513395 | 0.68099235 |
| VERM | 0.66988722 | 0.8883698 |
| HEM | 0.38479574 | 0.79925121 |
| FN | 0.8707758 | 0.92568707 |
| IP | 0.34144798 | 0.77275069 |
| DN | 0.07214301 | 0.68099235 |
| VENT | 0.0577405 | 0.68099235 |
| SPF | 0.25803823 | 0.72049635 |
| SPA | 0.56247209 | 0.8883698 |
| PP | 0.24727162 | 0.72049635 |
| GENd | 0.61882552 | 0.8883698 |
| LAT | 0.91693354 | 0.94779188 |
| ATN | 0.22862593 | 0.72049635 |
| MED | 0.18615219 | 0.68099235 |
| MTN | 0.49773969 | 0.88106721 |
| ILM | 0.13715774 | 0.68099235 |
| RT | 0.11118662 | 0.68099235 |
| GENv | 0.73897951 | 0.8883698 |
| EPI | 0.57211001 | 0.8883698 |
| SO | 0.69577456 | 0.8883698 |
| ASO | 0.5958894 | 0.8883698 |
| PVH | 0.34695293 | 0.77703001 |
| PVa | 0.75175345 | 0.8883698 |
| PVi | 0.80608841 | 0.8982942 |
| ARH | 0.51554683 | 0.88674055 |
| ADP | 0.83956841 | 0.92425134 |
| AVP | 0.29739471 | 0.73944458 |
| AVPV | 0.17861813 | 0.68099235 |
| DMH | 0.2577635 | 0.72049635 |
| MEPO | 0.08228319 | 0.68099235 |
| MPO | 0.58432967 | 0.8883698 |
| OV | 0.13424841 | 0.68099235 |
| PD | 0.72868802 | 0.8883698 |
| PS | 0.33200908 | 0.76754788 |
| PVp | 0.72895533 | 0.8883698 |
| PVpo | 0.25648885 | 0.72049635 |
| SBPV | 0.09922299 | 0.68099235 |
| SCH | 0.01831757 | 0.68099235 |
| SFO | 0.4179075 | 0.84764257 |
| VLPO | 0.38661454 | 0.79925121 |
| AHN | 0.15129479 | 0.68099235 |
| MBO | 0.17404181 | 0.68099235 |
| MPN | 0.21502807 | 0.72049635 |
| PMd | 0.29921711 | 0.73944458 |
| PMv | 0.49183504 | 0.88106721 |
| PVHd | 0.4850519 | 0.88106721 |
| VMH | 0.13670046 | 0.68099235 |
| PH | 0.54422189 | 0.8883698 |
| LHA | 0.50258828 | 0.88106721 |
| LPO | 0.36192686 | 0.7934956 |
| PST | 0.28875645 | 0.73907902 |
| PSTN | 0.44614136 | 0.88106721 |
| RCH | 0.19954659 | 0.68099235 |
| STN | 0.77705393 | 0.8883698 |
| TU | 0.31215862 | 0.75031315 |
| ZI | 0.18127427 | 0.68099235 |
| SCs | 0.22471443 | 0.72049635 |
| IC | 0.62416213 | 0.8883698 |
| NB | 0.76703956 | 0.8883698 |
| SAG | 0.47284921 | 0.88106721 |
| PBG | 0.32191668 | 0.76057238 |
| MEV | 0.80637573 | 0.8982942 |
| SNr | 0.55306018 | 0.8883698 |
| VTA | 0.04827669 | 0.68099235 |
| RR | 0.46660767 | 0.88106721 |
| MRN | 0.16134574 | 0.68099235 |
| SCm | 0.29301735 | 0.73944458 |
| PAG | 0.36685448 | 0.7934956 |
| PRT | 0.78587442 | 0.88927895 |
| CUN | 0.720967 | 0.8883698 |
| RN | 0.01900347 | 0.68099235 |
| III | 0.14009508 | 0.68099235 |
| EW | 0.31408457 | 0.75031315 |
| IV | 0.12280988 | 0.68099235 |
| VTN | 0.64096965 | 0.8883698 |
| AT | 0.56438108 | 0.8883698 |
| LT | 0.49201572 | 0.88106721 |
| SNC | 0.16412472 | 0.68099235 |
| PPN | 0.74040045 | 0.8883698 |
| RAmb | 0.10759762 | 0.68099235 |
| NLL | 0.16371381 | 0.68099235 |
| PSV | 0.13534368 | 0.68099235 |
| PB | 0.92443401 | 0.94918263 |
| SOC | 0.53599409 | 0.8883698 |
| B | 0.50405241 | 0.88106721 |
| DTN | 0.51039665 | 0.88496194 |
| PCG | 0.6068473 | 0.8883698 |
| PG | 0.26784257 | 0.73477471 |
| PRNc | 0.22967185 | 0.72049635 |
| SG | 0.34093633 | 0.77275069 |
| SUT | 0.28365721 | 0.73477471 |
| TRN | 0.05716134 | 0.68099235 |
| V | 0.72787302 | 0.8883698 |
| CS | 0.88694346 | 0.92569342 |
| LC | 0.75745057 | 0.8883698 |
| LDT | 0.72218454 | 0.8883698 |
| NI | 0.88392194 | 0.92569342 |
| PRNr | 0.16267473 | 0.68099235 |
| RPO | 0.9618055 | 0.96629992 |
| SLC | 0.05480754 | 0.68099235 |
| SLD | 0.07313899 | 0.68099235 |
| AP | 0.93567527 | 0.95341319 |
| CN | 0.02749249 | 0.68099235 |
| DCN | 0.71573777 | 0.8883698 |
| ECU | 0.67451466 | 0.8883698 |
| NTB | 0.49992408 | 0.88106721 |
| NTS | 0.1491332 | 0.68099235 |
| SPVC | 0.14583462 | 0.68099235 |
| SPVI | 0.18030031 | 0.68099235 |
| SPVO | 0.08324099 | 0.68099235 |
| VI | 0.60722706 | 0.8883698 |
| VII | 0.84257332 | 0.92425134 |
| ACVII | 0.0843698 | 0.68099235 |
| AMB | 0.1217536 | 0.68099235 |
| DMX | 0.86944024 | 0.92568707 |
| GRN | 0.19865812 | 0.68099235 |
| ICB | 0.09591202 | 0.68099235 |
| IO | 0.74087088 | 0.8883698 |
| IRN | 0.24369931 | 0.72049635 |
| ISN | 0.28309632 | 0.73477471 |
| LIN | 0.03861695 | 0.68099235 |
| LRN | 0.80472006 | 0.8982942 |
| MARN | 0.28047629 | 0.73477471 |
| MDRN | 0.85927474 | 0.92568707 |
| PARN | 0.43728046 | 0.87864765 |
| PAS | 0.13937487 | 0.68099235 |
| PGRN | 0.40355214 | 0.82632104 |
| PHY | 0.1619317 | 0.68099235 |
| PPY | 0.49863843 | 0.88106721 |
| VNC | 0.22522705 | 0.72049635 |
| x | 0.59156201 | 0.8883698 |
| XII | 0.85745385 | 0.92568707 |
| y | 0.06244422 | 0.68099235 |
| RM | 0.25358023 | 0.72049635 |
| RPA | 0.12041529 | 0.68099235 |
| RO | 0.72485467 | 0.8883698 |

To test whether pup calls engage different neural circuitry in females of different maternal experience, we correlated regional, normalized c-Fos expression across samples within our six conditions, producing six correlation matrices that represented interregional c-Fos correlations across the whole brain (**Figure 2C**). To quantify differences in brain-wide synchrony across conditions, we computed the average interregional correlation per matrix (**Figure 2D**). We found a significant effect of stimulus, as well as a significant interaction effect of group and stimulus, on average interregional correlation. This result demonstrated that pup calls (compared to baseline) had a significant effect on average interregional correlation across the whole brain, but that this effect differed per group. Indeed, compared to baseline, pup calls led to an increase in average brain-wide correlation in naïve virgins, while in the two maternally experienced groups, pup calls led to a decrease (**Figure 2D**, **Table 3**). Overall, when naïve animals perceive pup calls, we observe an increase in global synchrony. However, when experienced animals perceive pup calls, we observe a decrease in global synchrony, suggesting more localized responses in the maternally experienced groups.

**Table 3:** 2-way ANOVA results on brain-wide average interregional correlation. Tukey’s post-hoc tests on main comparisons of interest from the bar graph in Figure 2.

| Group 1 | Group 2 | p-value |
| --- | --- | --- |
| Naïve virgins + baseline | Naïve virgins + pup calls | < 0.0001 (****) |
| Experienced virgins + baseline | Experienced virgins + pup calls | 0.0001796<br>(***) |
| Mothers + baseline | Mothers + pup calls | 0.0004070<br>(***) |
| Naïve virgins + baseline | Experienced virgins + baseline | < 0.0001 (****) |
| Naïve virgins + baseline | Mothers + baseline | < 0.0001 (****) |
| Experienced virgins + baseline | Mothers + baseline | 0.347 (ns) |
| Naïve virgins + pup calls | Experienced virgins + pup calls | < 0.0001 (****) |
| Naïve virgins + pup calls | Mothers + pup calls | < 0.0001 (****) |
| Experienced virgins + pup calls | Mothers + pup calls | 0.4611 (ns) |

In addition to identifying an experience-dependent effect of pup calls on brain-wide coordination, our analysis also revealed that our three groups had significant differences in their correlated c-Fos expression at baseline (**Table 3**). To account for this fact and simplify further analyses, similar to how prior work has done (Jin et al., 2024; Rogers et al., 2026), we standardized each brain region’s pup call-exposed c-Fos expression to baseline expression within each group, calculating a measure we defined as *cFos*^*PC*^ per region (see **Methods**). As a result, *cFos*^*PC*^ represents each region’s c-Fos expression that we could attribute to pup call perception, accounting for baseline differences between naïve virgins, experienced virgins, and mothers (**Extended Figure 8**, **Supplemental Videos 8-10**). We then correlated regional *cFos*^*PC*^ per group to measure the coordinated recruitment of brain regions during pup call perception, relative to baseline (**Extended Figure 9**). It should be noted that since the definition of *cFos*^*PC*^is mathematically equivalent to Z-scoring the pup call data based on within-group baseline data, thus altering the mean but not the relationship between pup call-exposed c-Fos data points, the *cFos*^*PC*^ correlation matrices (**Extended Figure 9**) were identical to the pup call-exposed correlation matrices before baseline standardization (**Figure 2C**). Therefore, all further group comparisons were focused on *cFos*^*PC*^ correlation analyses.

### Recruitment of Pup Avoidance and Parental Care Circuits upon pup call exposure reveal group differences

Foundational work in the field of maternal behavior across various species has led to the definition of two functionally opposing circuits that control (1) pup-directed avoidance or aggression (going forward, referred to as “pup avoidance”) and (2) parental behavior (Dulac et al., 2014) (**Figure 3A**). While these circuits were defined by drawing upon studies that largely examined multisensory pup interactions, we decided to use these circuits as a priori regions of interest. We correlated *cFos*^*PC*^ across each subset of regions, allowing us to assess coordinated recruitment of these circuits upon pup call exposure in naïve virgins, experienced virgins, and mothers (**Figure 3B-C**). It is worth noting that we had to separate or combine some regions from the originally proposed circuits to map onto our level of Allen Brain Atlas parcellation specificity – for example, Dulac et al. refer to the prefrontal cortex (PFC), which we break down into the frontal pole (FRP), infralimbic area (ILA), dorsal and ventral anterior cingulate areas (ACAd, ACAv), prelimbic area (PL), and the lateral, medial, and ventrolateral orbital areas (ORBl, ORBm, ORBvl). To assess how these two circuits are similarly or differently recruited across groups, we first calculated the average interregional correlation to compare overall circuit synchrony, finding no significant group differences in either circuit (**Extended Figure 10**).

**Figure 3:**
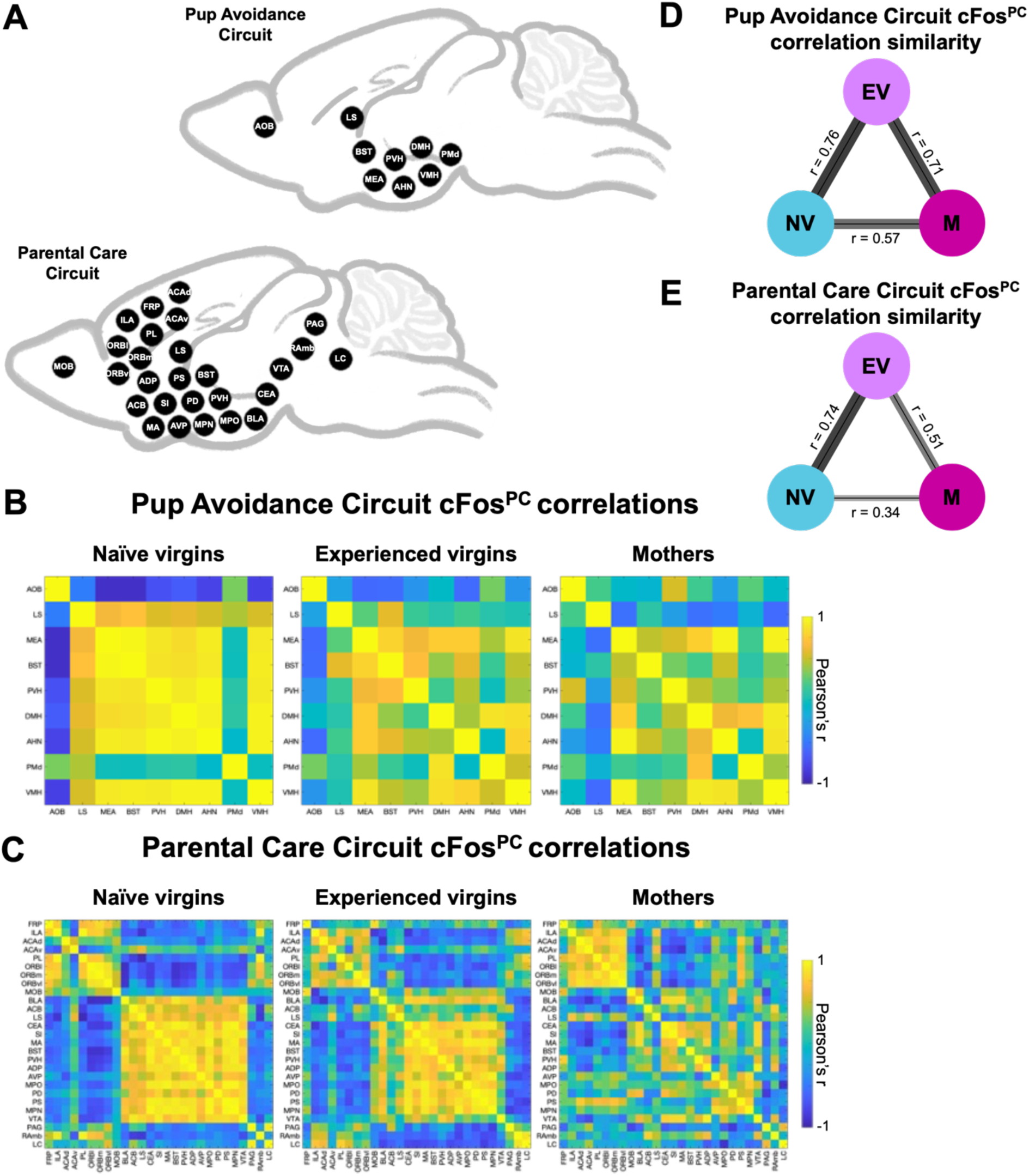
Pup Avoidance and Parental Care Circuits reveal different coordinated activation patterns upon pup call exposure between groups. (A) Regions within the Pup Avoidance and Parental Care Circuits, adapted from Dulac et al., 2014 (see Table 1 for abbreviations). (B) Pearson correlation matrices indicating interregional correlations in Pup Avoidance Circuit, based on ***cFos***^***PC***^ in pup call-exposed samples across groups (n = 8 x 9 regions). Matrix pixel colors represent Pearson correlation coefficients, see scale to the right. (C) Pearson correlation matrices indicating interregional correlations in Parental Care Circuit, based on ***cFos***^***PC***^in pup call-exposed samples across groups (n = 8 x 27 regions). Matrix pixel colors represent Pearson correlation coefficients, see scale to the right. (D) Graphical representation of Pup Avoidance Circuit matrix similarity between groups (NV = naïve virgins, EV = experienced virgins, M = mothers), determined via Mantel test (naïve virgins vs. experienced virgins: p < 0.0001; naïve virgins vs. mothers: p = 0.000412; experienced virgins vs. mothers: p < 0.0001; n = 8 x 9 regions). Edge thickness and transparency are proportional to r value. (E) Graphical representation of Parental Care Circuit matrix similarity between groups, determined via Mantel test (naïve virgins vs. experienced virgins: p < 0.0001; naïve virgins vs. mothers: p < 0.0001; experienced virgins vs. mothers: p < 0.0001; n = 8 x 27 regions). Edge thickness and transparency are proportional to r value.

However, the circuits appeared to have group-level differences in their spatial correlation patterns. To quantify and test this, we computed the similarity of these two circuits’ correlation matrices across groups (**Figure 3D-E**). In the Pup Avoidance Circuit, our results indicated that our three groups were relatively similar to each other, albeit with naïve virgins and mothers being the least similar pair (**Figure 3D**). In the Parental Care Circuit, however, there was more variation in the matrix similarities, with naïve virgins and experienced virgins being the most similar, experienced virgins and mothers being moderately similar, and naïve virgins and mothers being most dissimilar from each other (**Figure 3E**). Intriguingly, this gradient in similarity follows the gradient of maternal experience across groups. Furthermore, the fact that naïve virgins and experienced virgins showed the highest similarity in the Parental Care Circuit upon pup call exposure coincides with the surprising result of our auditory two-choice assay, which showed that, like naïve virgins, experienced virgins do not prefer pup calls, even though they do retrieve pups.

### Maternal experience leads to a more fine-tuned pup call response network

To identify how maternal experience alters the brain-wide network of regions recruited during pup call perception, we applied stringent significance thresholding to our three groups’ brain-wide *cFos*^*PC*^ correlation matrices (**Extended Figure 9**) and corrected for multiple comparisons. The surviving pairwise correlations were used to define pup call response networks, in which each node is one of our 215 brain regions, and each edge is a surviving *cFos*^*PC*^ correlation between two regions (**Figure 4A**). Then, the network density, average node degree, and average correlation magnitude were calculated for the pup call response network of naïve virgins, experienced virgins, and mothers, and group-level differences were assessed via permutation testing (**Figure 4B-D**).

**Figure 4:**
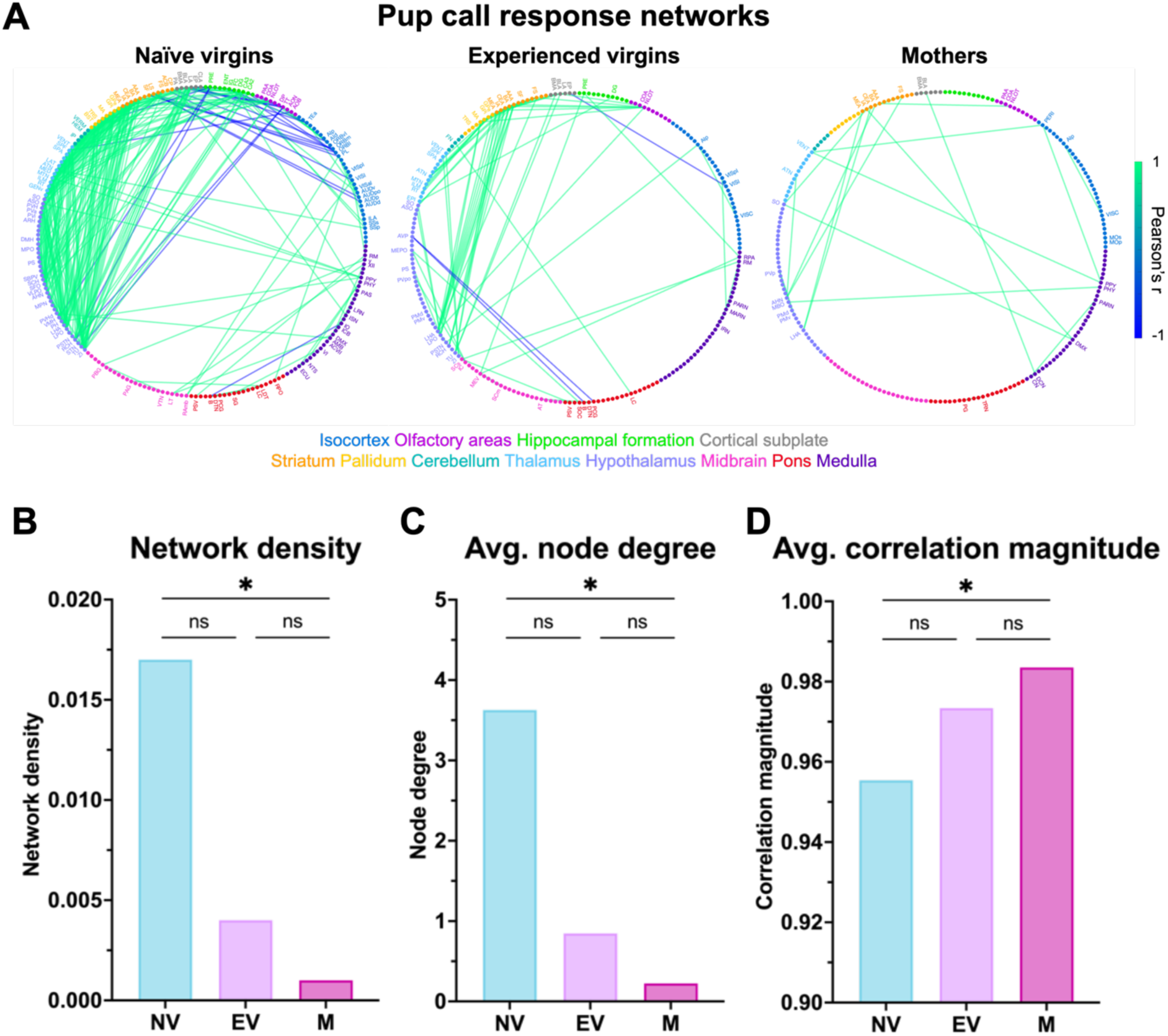
Maternal experience leads to sparser, but stronger, pup call response networks. (A) Pup call response networks, represented as undirected graphs, derived from ***cFos***^***PC***^correlation matrices, thresholded at FDR < 0.05 and corrected for multiple comparisons via the Benjamini, Krieger and Yekutieli 2-stage step-up method. Nodes represent 215 brain regions, node marker colors correspond to broader brain areas according to the Allen Brain Atlas, see text below graphs for color coding. Only nodes that participate in edges are labeled (see Table 1 for abbreviations). Edge color represents Pearson correlation coefficients, see scale to the right. (B) Network density per group’s pup call response network (NV = naïve virgins, EV = experienced virgins, M = mothers). Comparisons on graph derived from permutation tests (naïve virgins vs. experienced virgins: p = 0.094; naïve virgins vs. mothers: p = 0.038; experienced virgins vs. mothers: p = 0.309; n = 8 x 215 regions; n = 1,000 permutations). Bars denote observed values. * p < 0.05. (C) Average unweighted node degree per group’s pup call response network. Comparisons on graph derived from permutation tests (naïve virgins vs. experienced virgins: p = 0.094; naïve virgins vs. mothers: p = 0.038; experienced virgins vs. mothers: p = 0.308; n = 8 x 215 regions; n = 1,000 permutations). Bars denote observed values. * p < 0.05. (D) Average correlation magnitude per each group’s pup call response network. Comparisons on graph derived from permutation tests (naïve virgins vs. experienced virgins: p = 0.098492; naïve virgins vs. mothers: p = 0.020325; experienced virgins vs. mothers: p = 0.2438; n = 8 x 215 regions; n= 1,000 permutations). Bars denote observed values. * p < 0.05.

We found that naïve virgins had the highest pup call response network density, followed by experienced virgins, and then mothers (**Figure 4B**). The same pattern was revealed for average node degree (**Figure 4C**, **Table 4**), and the reverse pattern was found for average correlation magnitude (**Figure 4D**, **Table 5**). These three network measures once again identified gradients in the brain-wide representation of pup calls that parallel the gradient of maternal experience across groups. Permutation tests of these network measures confirmed that mothers had a significantly lower network density, lower average node degree, and higher average correlation magnitude, compared to naïve virgins, setting these two groups apart as the most distinct. Comparisons between naïve virgins versus experienced virgins, as well as between experienced virgins versus mothers, revealed trending though non-significant results, suggesting that experienced virgins represent an intermediate phenotype between naïve virgins and mothers that is more similar to mothers. Given that network density is particularly affected by the thresholding used to turn *cFos*^*PC*^ correlation matrices into pup call response networks, we conducted a sensitivity analysis to confirm that our results held regardless of threshold choice, which we found to be true (**Table 6**). Taken together, these network-level analyses revealed that maternally experienced animals exhibit a sparser, yet stronger, pup call response network, compared to naïve animals. This finding reveals a more specialized brain-wide pup call response that is reflected in pup retrieval success in experienced animals.

**Table 4:**
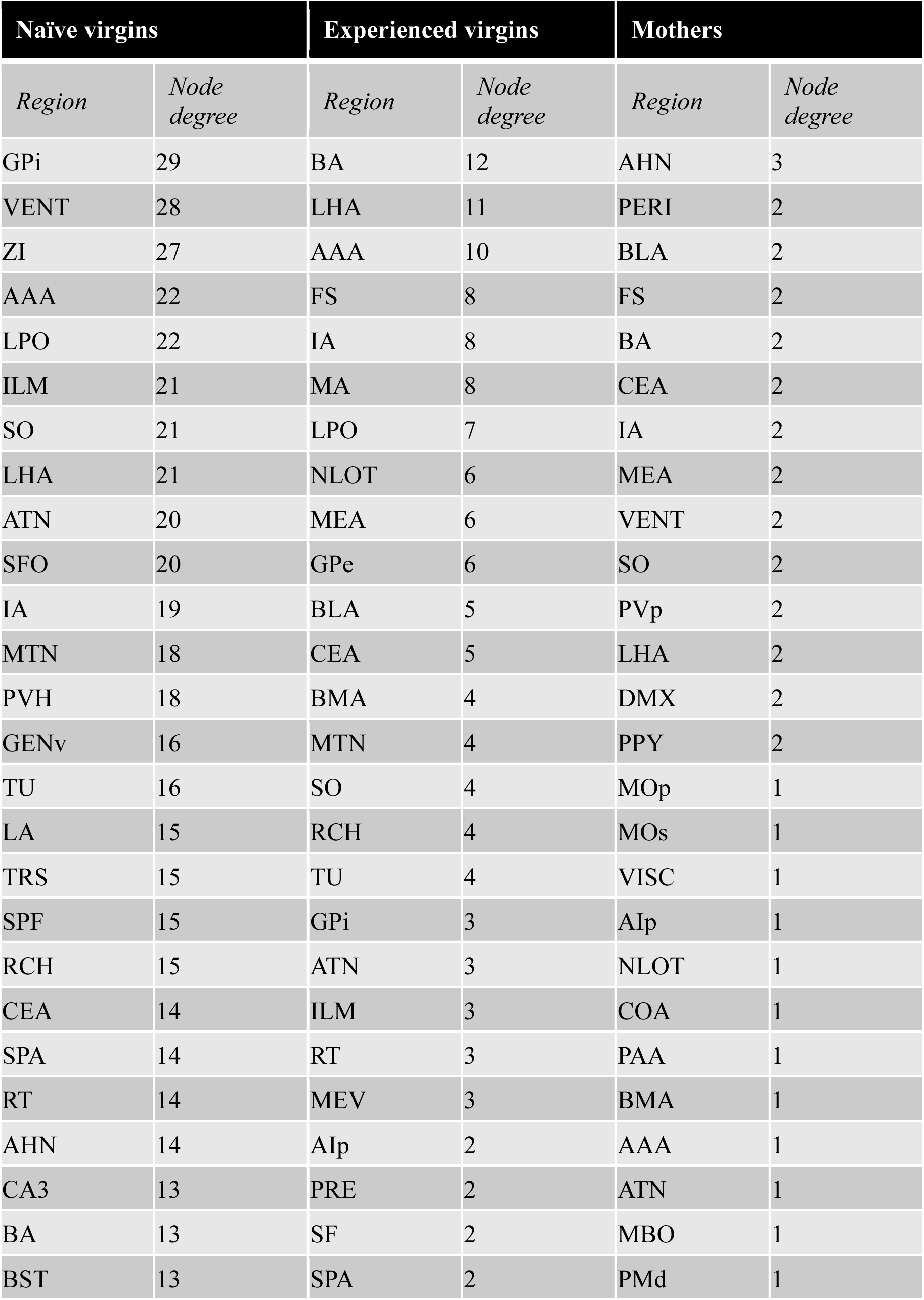
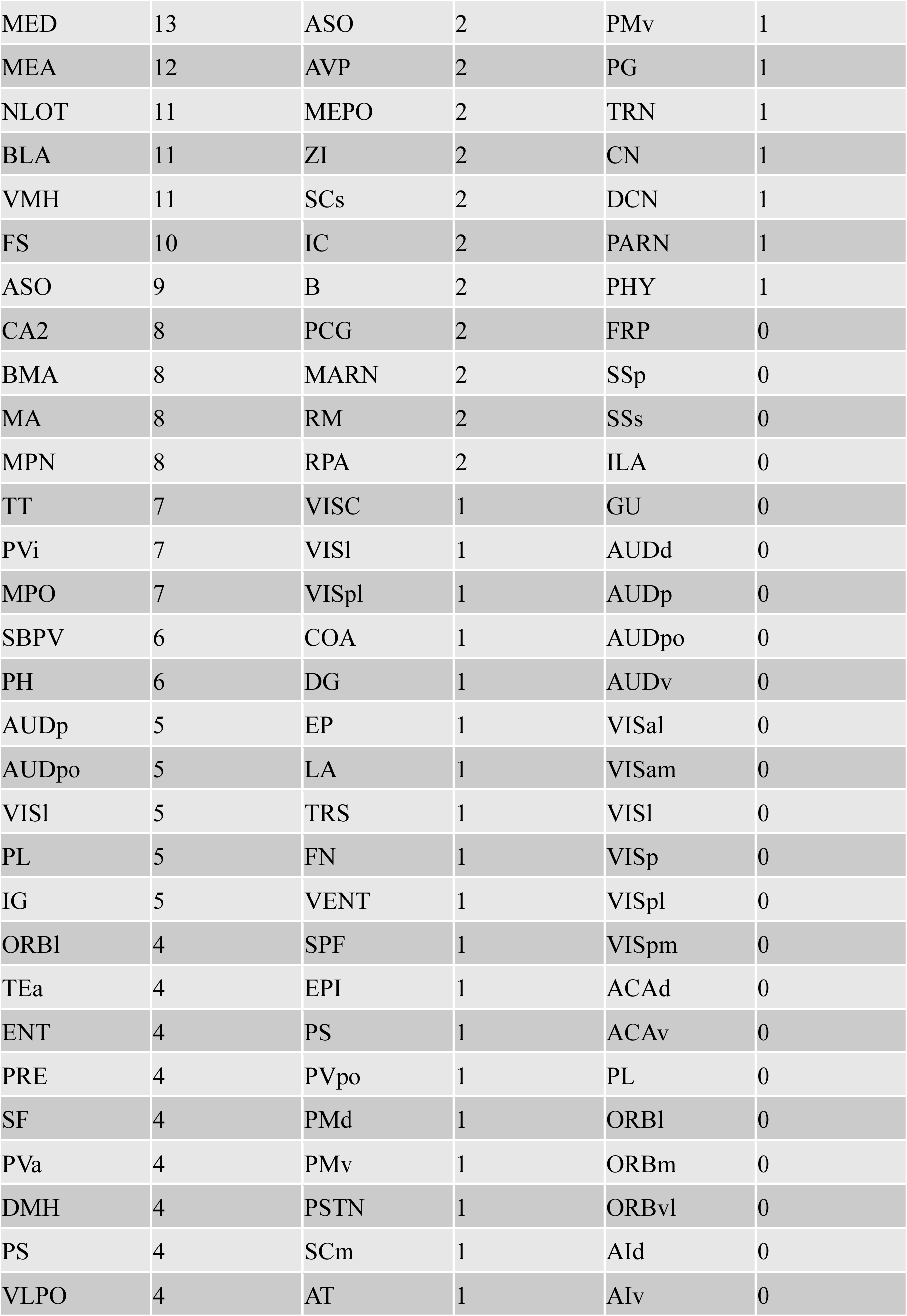
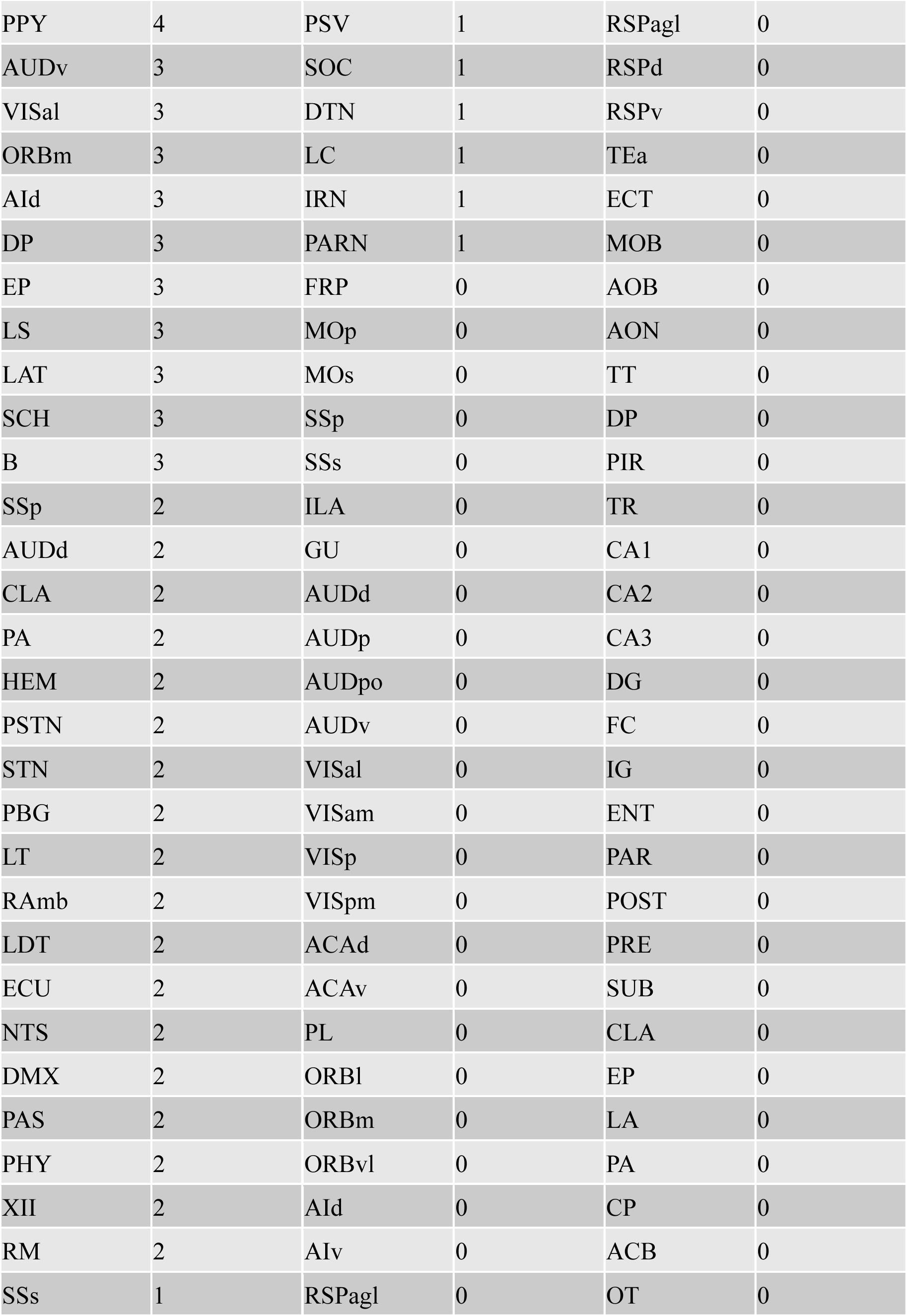
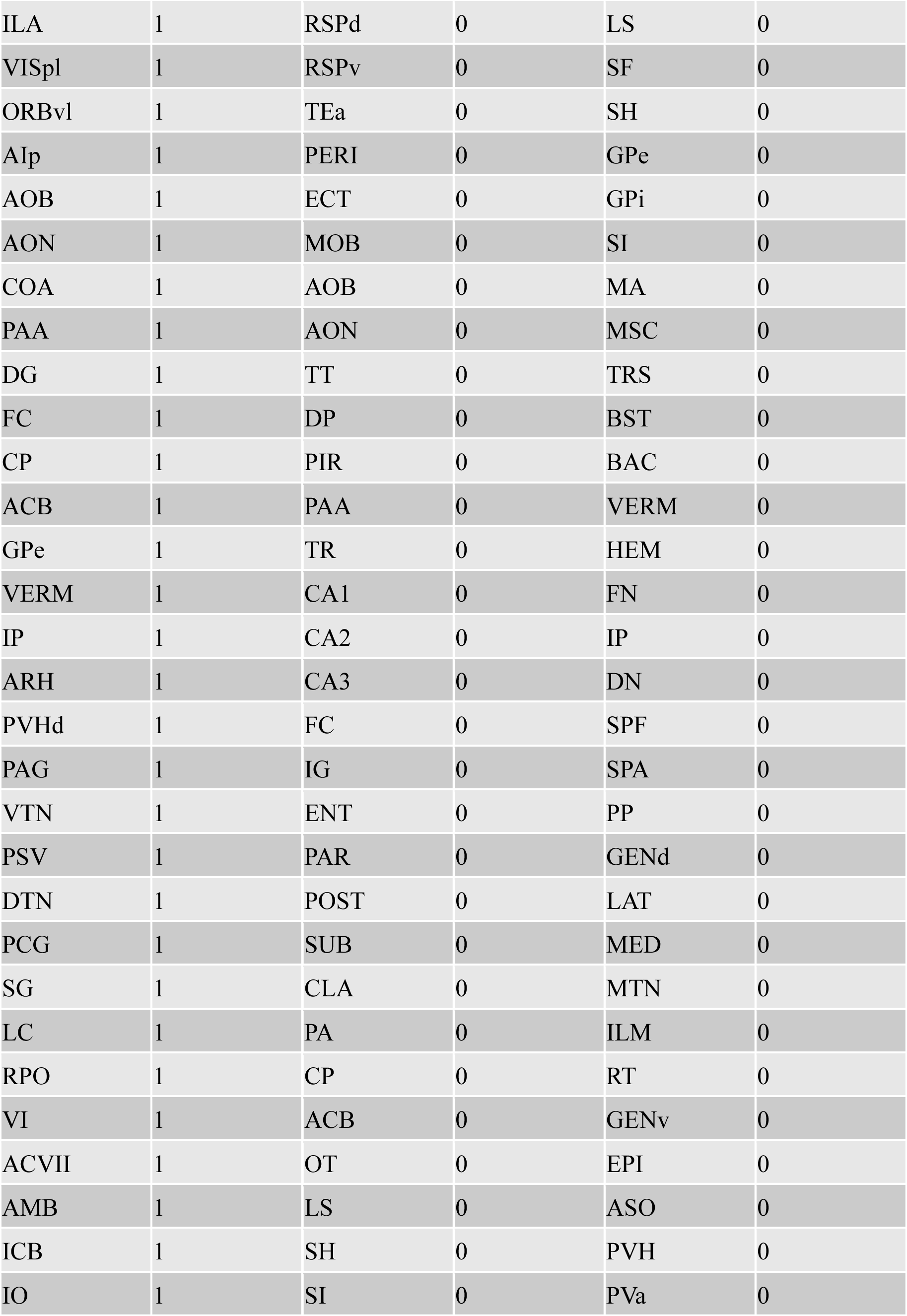
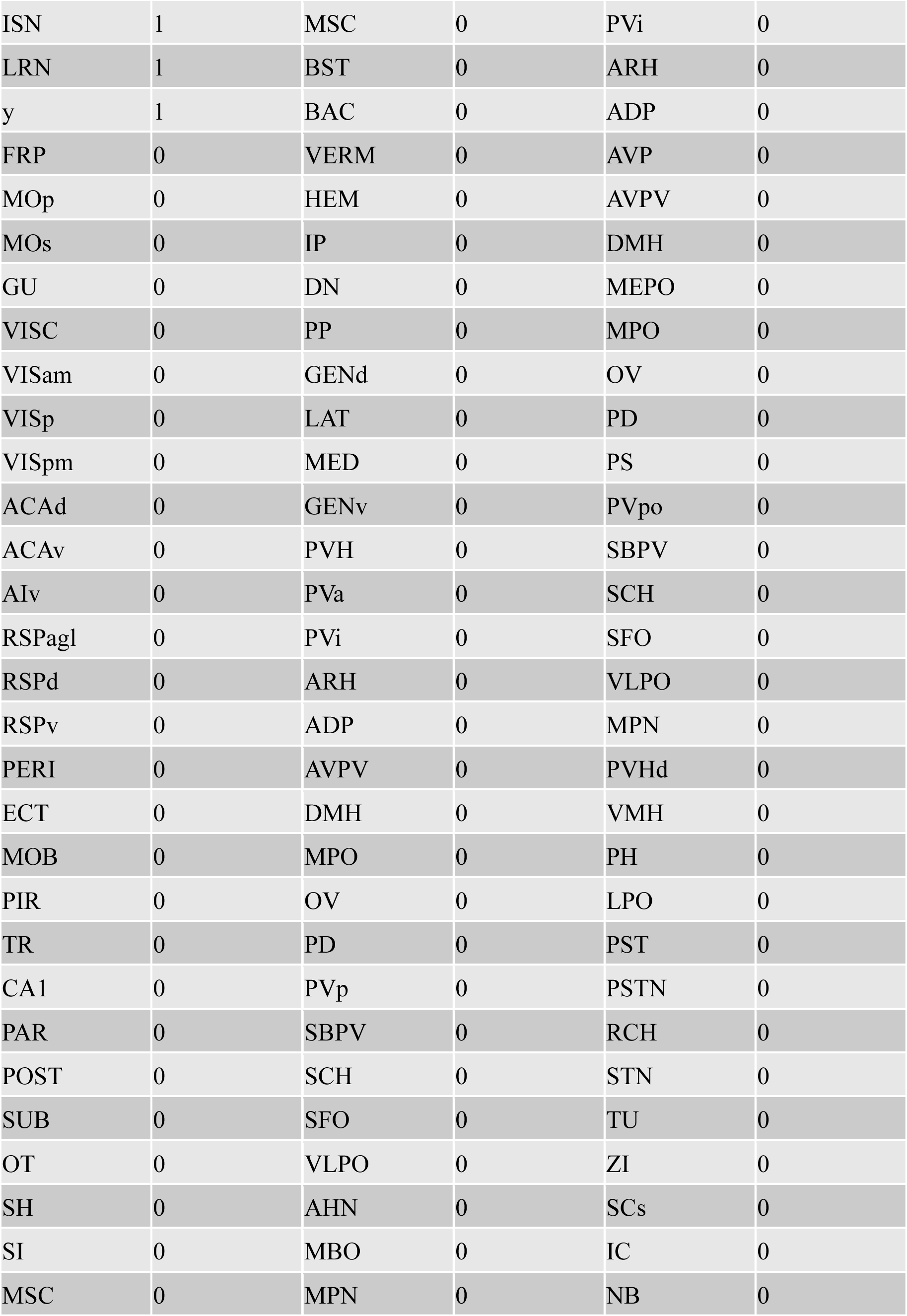
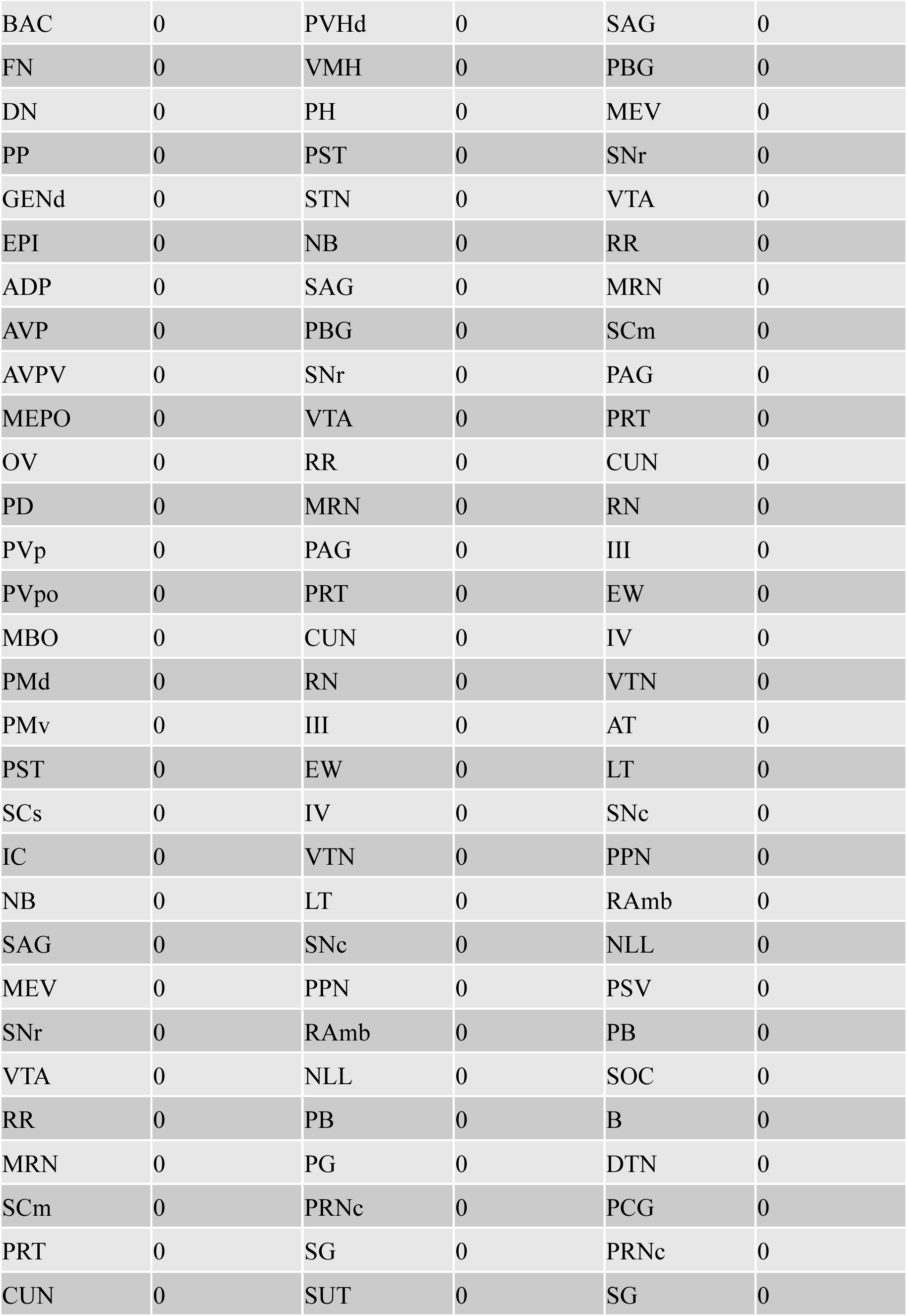
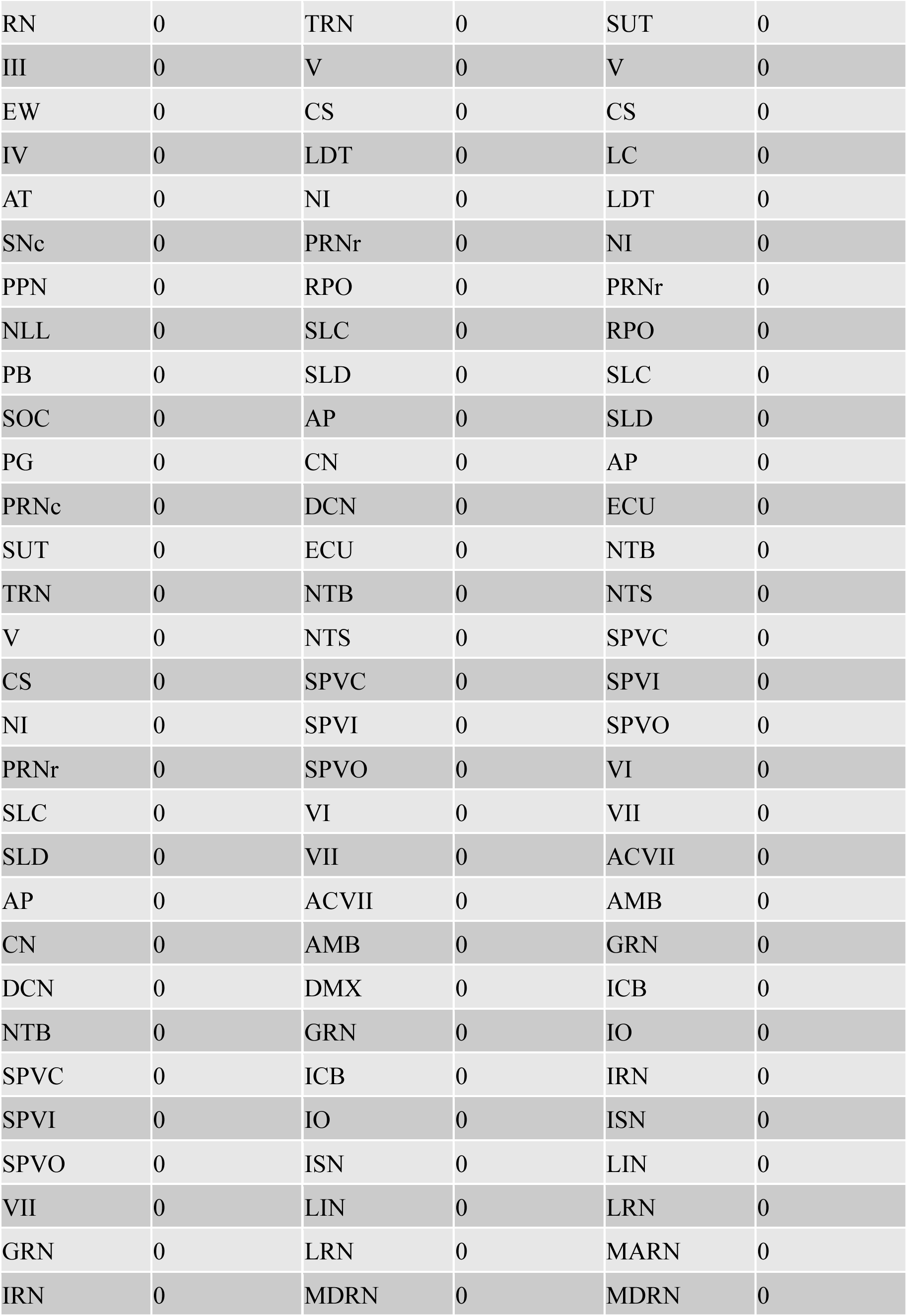
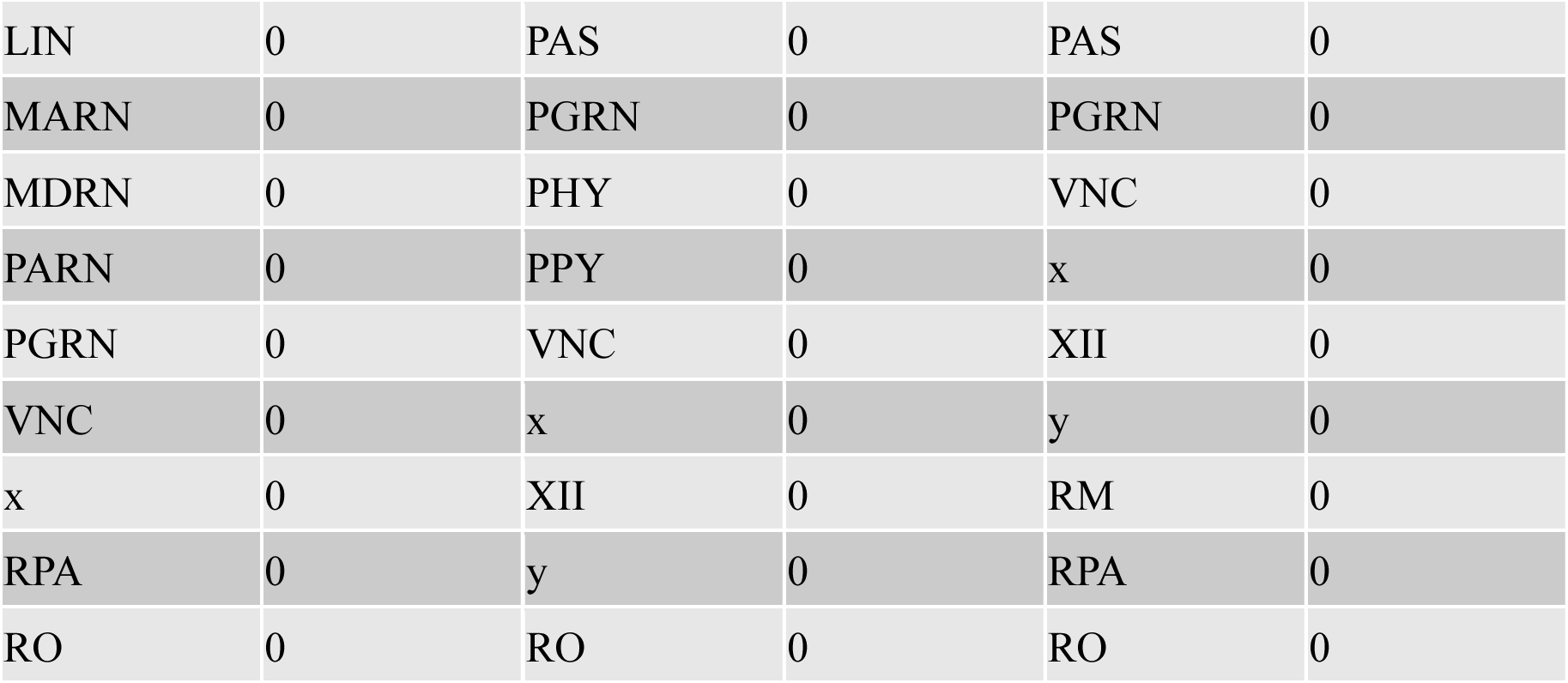
Node degree per each group’s pup call response network. For each of the 215 brain regions conserved from Allen Brain Atlas, this table lists their abbreviation (see Table 1 for abbreviations) and node degree in our pup call response networks per group, in descending degree order.

**Table 5:** Correlation magnitude per each group’s pup call response network. For each edge, this table lists the 2 regions’ abbreviation (see Table 1 for abbreviations) and their correlation magnitude in descending magnitude order.

| Naïve virgins |  |  | Experienced virgins |  |  | Mothers |  |  |
| --- | --- | --- | --- | --- | --- | --- | --- | --- |
| *Region 1* | *Region 2* | $\vert r\vert $ | *Region 1* | *Region 2* | $\vert r\vert $ | *Region 1* | *Region 2* | $\vert r\vert $ |
| FS | SPA | 0.9952591 | GPI | RT | 0.99472199 | MEA | LHA | 0.99654035 |
| GPI | ZI | 0.99491456 | AAA | BA | 0.99432302 | VENT | PPY | 0.99612666 |
| SPA | LPO | 0.99460961 | BMA | MEA | 0.99238565 | CEA | MEA | 0.9949246 |
| ATN | MTN | 0.99439707 | FS | IA | 0.99061804 | CN | PARN | 0.99357551 |
| GPI | VENT | 0.99434908 | IA | LHA | 0.98997118 | CEA | LHA | 0.99318389 |
| ECU | NTS | 0.99387602 | VENT | ZI | 0.98990426 | PERI | VENT | 0.98699671 |
| GEN<sub>v</sub> | SFO | 0.99276357 | FS | LHA | 0.9898262 | FS | BA | 0.98632598 |
| ASO | PH | 0.99209735 | SCs | PSV | 0.98949698 | MO<sub>p</sub> | MO<sub>s</sub> | 0.9852229 |
| ECU | PHY | 0.99202644 | IA | MEA | 0.98926609 | PERI | PPY | 0.98488693 |
| VENT | ZI | 0.99040328 | PRE | RPA | 0.98917445 | NLOT | AAA | 0.9847623 |
| RCH | TU | 0.99028434 | AAA | LHA | 0.98866741 | IA | SO | 0.98469854 |
| LA | VMH | 0.9894186 | PV<sub>po</sub> | IC | 0.98781784 | FS | AHN | 0.98391291 |
| CEA | LHA | 0.98894142 | PM<sub>d</sub> | PM<sub>v</sub> | 0.98764963 | COA | PAA | 0.9829132 |
| MED | RT | 0.98748092 | BA | CEA | 0.98763123 | IA | DMX | 0.98174173 |
| AAA | ILM | 0.98715511 | MEA | TU | 0.98702168 | PV<sub>p</sub> | PM<sub>d</sub> | 0.98082613 |
| BMA | TU | 0.98669667 | BA | LHA | 0.98682915 | SO | DMX | 0.98051936 |
| ILM | SFO | 0.98612156 | SO | ASO | 0.98628052 | VISC | AI<sub>p</sub> | 0.97856717 |
| VENT | LHA | 0.98601948 | CEA | RCH | 0.98509411 | BLA | BMA | 0.97816897 |
| MTN | ILM | 0.98580413 | MEV | SC<sub>m</sub> | 0.98413823 | PV<sub>p</sub> | PM<sub>v</sub> | 0.97692784 |
| TRS | ZI | 0.98541654 | ILM | RT | 0.98340026 | ATN | AHN | 0.97688804 |
| ATN | ILM | 0.98523099 | GPI | ILM | 0.98188737 | BLA | DCN | 0.97642629 |
| MED | ILM | 0.98438092 | CEA | ATN | 0.98169254 | MBO | PHY | 0.9741497 |
| NTS | PHY | 0.98368391 | PRE | MEV | 0.98152868 | PG | TRN | 0.97364022 |
| ILM | RT | 0.98355037 | DG | EPI | 0.97916482 | BA | AHN | 0.97284977 |
| AAA | LPO | 0.98347653 | SO | LPO | 0.97905518 |  |  |  |
| LT | DMX | 0.98346252 | NLOT | LHA | 0.97819672 |  |  |  |
| PVi | SBPV | 0.98318204 | BA | LPO | 0.97802404 |  |  |  |
| IA | VENT | 0.9830418 | BMA | IA | 0.97798901 |  |  |  |
| IA | ZI | 0.98272906 | MEV | RPA | 0.97758972 |  |  |  |
| SPA | SO | 0.98237542 | FS | AAA | 0.97671324 |  |  |  |
| MA | VLPO | 0.98196213 | CEA | LHA | 0.97618839 |  |  |  |
| GPi | LHA | 0.98192152 | AAA | CEA | 0.97555113 |  |  |  |
| MTN | RT | 0.98183476 | SPA | MTN | 0.97545325 |  |  |  |
| PVi | VMH | 0.98181379 | NLOT | FS | 0.97534644 |  |  |  |
| FS | LPO | 0.98175226 | AT | DTN | 0.97519767 |  |  |  |
| SFO | ZI | 0.98144224 | BA | MA | 0.97518303 |  |  |  |
| FS | SO | 0.98109781 | PS | LPO | 0.97457085 |  |  |  |
| VISI | RM | 0.98087608 | COA | BLA | 0.97426579 |  |  |  |
| GPi | SFO | 0.98068408 | BA | SO | 0.97369113 |  |  |  |
| TRS | MED | 0.98059598 | NLOT | IA | 0.97342729 |  |  |  |
| MEA | TU | 0.97993239 | GPi | MTN | 0.97272578 |  |  |  |
| PVi | DMH | 0.97975789 | MEPO | SOC | 0.97259141 |  |  |  |
| NLOT | IA | 0.97962748 | AAA | SO | 0.9724462 |  |  |  |
| IA | MEA | 0.97937728 | B | PCG | 0.97223253 |  |  |  |
| PA | ISN | 0.9790929 | FN | LC | 0.97214086 |  |  |  |
| NLOT | MEA | 0.97900433 | MTN | RT | 0.97183629 |  |  |  |
| BST | RCH | 0.97872866 | LA | BLA | 0.97159017 |  |  |  |
| AAA | SO | 0.97819026 | BLA | BMA | 0.97158871 |  |  |  |
| AAA | SPA | 0.97807482 | BA | ATN | 0.97126504 |  |  |  |
| TRS | ILM | 0.97791398 | AAA | IA | 0.97110547 |  |  |  |
| NLOT | SPF | 0.97777278 | BA | RCH | 0.97105915 |  |  |  |
| VENT | ILM | 0.97757837 | AAA | RCH | 0.97089587 |  |  |  |
| AHN | LHA | 0.97749651 | FS | BA | 0.97077276 |  |  |  |
| ILM | ZI | 0.97724338 | AAA | LPO | 0.97038993 |  |  |  |
| AAA | SFO | 0.97630995 | NLOT | AAA | 0.97031217 |  |  |  |
| MEA | LHA | 0.97630575 | MEPO | IC | 0.97002632 |  |  |  |
| TRS | RT | 0.976241 | MEA | LHA | 0.96999558 |  |  |  |
| TRS | SFO | 0.97556335 | GPe | MA | 0.96963311 |  |  |  |
| VENT | SFO | 0.97553939 | SPF | ZI | 0.9696289 |  |  |  |
| PH | PPY | 0.97553101 | MA | LHA | 0.96887988 |  |  |  |
| ATN | RT | 0.9749498 | LHA | RCH | 0.96855914 |  |  |  |
| DMH | SBPV | 0.97485407 | SCs | RM | 0.96829362 |  |  |  |
| IA | GPI | 0.97434058 | GPe | LHA | 0.96808389 |  |  |  |
| PVH | AHN | 0.97389966 | BMA | TU | 0.96742229 |  |  |  |
| LHA | ZI | 0.97383621 | BA | GPe | 0.96707846 |  |  |  |
| MED | MTN | 0.97374021 | MARN | RM | 0.96658407 |  |  |  |
| MEA | BST | 0.97354425 | NLOT | MA | 0.96612243 |  |  |  |
| RT | ZI | 0.97330109 | VISC | AIp | 0.96563265 |  |  |  |
| BMA | RCH | 0.97327508 | AAA | MA | 0.96531968 |  |  |  |
| AAA | IA | 0.97322876 | FS | MA | 0.96457409 |  |  |  |
| MA | SPA | 0.9731978 | MA | LPO | 0.96420684 |  |  |  |
| HEM | PSTN | 0.97273064 | BLA | MEA | 0.96337629 |  |  |  |
| SO | LPO | 0.97260045 | GPe | SPA | 0.96273893 |  |  |  |
| CEA | AHN | 0.97225469 | GPe | ATN | 0.9626597 |  |  |  |
| MA | LPO | 0.97225467 | SF | TRS | 0.96263163 |  |  |  |
| AUDp | AUDpo | 0.97217598 | IA | TU | 0.96257061 |  |  |  |
| BA | SO | 0.97204873 | FS | GPe | 0.96243871 |  |  |  |
| MEA | VENT | 0.97204273 | BLA | TU | 0.96142944 |  |  |  |
| VENT | RT | 0.9719636 | AIp | PSTN | 0.96115449 |  |  |  |
| CEA | MEA | 0.97191415 | FS | MEA | 0.96106396 |  |  |  |
| IA | ILM | 0.97186286 | BA | IA | 0.96085709 |  |  |  |
| ILM | SO | 0.97157189 | IRN | PARN | 0.95915673 |  |  |  |
| MEA | RCH | 0.97148221 | MTN | ILM | 0.95896409 |  |  |  |
| ATN | SFO | 0.97147958 | LHA | LPO | 0.95851458 |  |  |  |
| CA3 | GPI | 0.97088741 | NLOT | BA | 0.95846941 |  |  |  |
| ATN | MED | 0.97077736 | ASO | LPO | 0.95807675 |  |  |  |
| DMX | XII | 0.97055248 | VISpl | MARN | 0.95804401 |  |  |  |
| RT | SFO | 0.97048505 | SF | MA | 0.95788101 |  |  |  |
| MTN | SFO | 0.97043519 | VISl | EP | -0.9577848 |  |  |  |
| GPI | ILM | 0.97027041 | AVP | B | -0.9595474 |  |  |  |
| CA2 | RCH | 0.97016889 | AVP | PCG | -0.9636026 |  |  |  |
| AUDpo | VISal | 0.96957952 |  |  |  |  |  |  |
| TEa | ENT | 0.96949133 |  |  |  |  |  |  |
| ILA | PL | 0.96941203 |  |  |  |  |  |  |
| MEA | ZI | 0.96906687 |  |  |  |  |  |  |
| ILM | GENv | 0.96906262 |  |  |  |  |  |  |
| GPI | TRS | 0.96903616 |  |  |  |  |  |  |
| BST | TU | 0.96875415 |  |  |  |  |  |  |
| MTN | GENv | 0.96827329 |  |  |  |  |  |  |
| AAA | VENT | 0.96823413 |  |  |  |  |  |  |
| MEA | GPI | 0.96815153 |  |  |  |  |  |  |
| CEA | MPN | 0.96804623 |  |  |  |  |  |  |
| GPI | RT | 0.96782524 |  |  |  |  |  |  |
| ATN | GENv | 0.96750114 |  |  |  |  |  |  |
| MPO | MPN | 0.96743018 |  |  |  |  |  |  |
| IA | SPF | 0.96719969 |  |  |  |  |  |  |
| LHA | RCH | 0.96692519 |  |  |  |  |  |  |
| RT | GENv | 0.96690115 |  |  |  |  |  |  |
| VERM | IP | 0.96668883 |  |  |  |  |  |  |
| LT | XII | 0.96662342 |  |  |  |  |  |  |
| CA3 | STN | 0.96660385 |  |  |  |  |  |  |
| SBPV | VMH | 0.96607217 |  |  |  |  |  |  |
| TT | DP | 0.96561784 |  |  |  |  |  |  |
| MED | ZI | 0.96560196 |  |  |  |  |  |  |
| TU | ZI | 0.96518505 |  |  |  |  |  |  |
| MPO | PS | 0.96516151 |  |  |  |  |  |  |
| PS | MPN | 0.96508606 |  |  |  |  |  |  |
| SO | ASO | 0.96462351 |  |  |  |  |  |  |
| GPI | GEN<sub>v</sub> | 0.96459797 |  |  |  |  |  |  |
| MTN | SO | 0.96456062 |  |  |  |  |  |  |
| AAA | MTN | 0.96451871 |  |  |  |  |  |  |
| ORB<sub>vl</sub> | AON | 0.96442254 |  |  |  |  |  |  |
| TRS | GEN<sub>v</sub> | 0.96439375 |  |  |  |  |  |  |
| FS | AAA | 0.96430388 |  |  |  |  |  |  |
| ATN | SO | 0.96401986 |  |  |  |  |  |  |
| SFO | LPO | 0.96395729 |  |  |  |  |  |  |
| GEN<sub>v</sub> | ZI | 0.96351926 |  |  |  |  |  |  |
| AAA | ZI | 0.96351911 |  |  |  |  |  |  |
| PVH | SFO | 0.96348979 |  |  |  |  |  |  |
| PVH<sub>d</sub> | AMB | 0.96253001 |  |  |  |  |  |  |
| MPO | LPO | 0.96226878 |  |  |  |  |  |  |
| CA2 | BLA | 0.96224772 |  |  |  |  |  |  |
| BA | IA | 0.96221187 |  |  |  |  |  |  |
| AHN | MPN | 0.96207821 |  |  |  |  |  |  |
| IA | MED | 0.96198174 |  |  |  |  |  |  |
| BLA | TU | 0.96185937 |  |  |  |  |  |  |
| LA | SCH | 0.96166177 |  |  |  |  |  |  |
| VMH | RCH | 0.96161336 |  |  |  |  |  |  |
| TRS | VENT | 0.96151825 |  |  |  |  |  |  |
| AAA | MED | 0.96140966 |  |  |  |  |  |  |
| ASO | PPY | 0.96136166 |  |  |  |  |  |  |
| SO | SFO | 0.96117071 |  |  |  |  |  |  |
| ORB<sub>l</sub> | ORB<sub>m</sub> | 0.96093567 |  |  |  |  |  |  |
| MEA | SPF | 0.96082241 |  |  |  |  |  |  |
| ATN | PH | 0.96052096 |  |  |  |  |  |  |
| B | LC | 0.96038324 |  |  |  |  |  |  |
| PV<sub>a</sub> | PV<sub>i</sub> | 0.96014675 |  |  |  |  |  |  |
| CEA | VENT | 0.96003441 |  |  |  |  |  |  |
| MED | SFO | 0.95995925 |  |  |  |  |  |  |
| LA | RCH | 0.95979027 |  |  |  |  |  |  |
| CA<sub>3</sub> | VENT | 0.95952875 |  |  |  |  |  |  |
| ILM | LPO | 0.95927037 |  |  |  |  |  |  |
| AAA | ATN | 0.95898671 |  |  |  |  |  |  |
| A<sub>ld</sub> | DP | 0.95876644 |  |  |  |  |  |  |
| IA | TRS | 0.95855717 |  |  |  |  |  |  |
| SO | PVH | 0.95845407 |  |  |  |  |  |  |
| CA<sub>2</sub> | TU | 0.95844856 |  |  |  |  |  |  |
| BST | LHA | 0.95840109 |  |  |  |  |  |  |
| BA | VENT | 0.95828109 |  |  |  |  |  |  |
| VENT | SO | 0.95820454 |  |  |  |  |  |  |
| BMA | MEA | 0.95818082 |  |  |  |  |  |  |
| GP<sub>i</sub> | TU | 0.95808833 |  |  |  |  |  |  |
| IA | LHA | 0.95797625 |  |  |  |  |  |  |
| CEA | BST | 0.95774687 |  |  |  |  |  |  |
| BMA | BST | 0.9571536 |  |  |  |  |  |  |
| AUD<sub>v</sub> | TE<sub>a</sub> | 0.95700799 |  |  |  |  |  |  |
| SPF | ZI | 0.95691215 |  |  |  |  |  |  |
| PVH | LHA | 0.95668296 |  |  |  |  |  |  |
| PVH | LPO | 0.95649433 |  |  |  |  |  |  |
| ICB | PAS | 0.95648235 |  |  |  |  |  |  |
| GP<sub>i</sub> | RCH | 0.95621026 |  |  |  |  |  |  |
| CA3 | BA | 0.95617482 |  |  |  |  |  |  |
| FS | MA | 0.95613224 |  |  |  |  |  |  |
| AUDd | VISal | 0.95587856 |  |  |  |  |  |  |
| SPA | MPO | 0.95585075 |  |  |  |  |  |  |
| GPI | SPF | 0.95577861 |  |  |  |  |  |  |
| VMH | LHA | 0.95553031 |  |  |  |  |  |  |
| LA | GPI | 0.95530294 |  |  |  |  |  |  |
| MA | MPO | 0.95504784 |  |  |  |  |  |  |
| MTN | PH | 0.95501881 |  |  |  |  |  |  |
| AAA | GPI | 0.95471232 |  |  |  |  |  |  |
| RCH | ZI | 0.95463573 |  |  |  |  |  |  |
| SFO | LHA | 0.95445523 |  |  |  |  |  |  |
| IA | RT | 0.95414325 |  |  |  |  |  |  |
| NLOT | TU | 0.95405248 |  |  |  |  |  |  |
| AAA | BA | 0.9539877 |  |  |  |  |  |  |
| IA | SFO | 0.95393571 |  |  |  |  |  |  |
| SSs | DG | 0.95388305 |  |  |  |  |  |  |
| SSp | CA3 | 0.95387948 |  |  |  |  |  |  |
| VENT | GENv | 0.95377795 |  |  |  |  |  |  |
| PAG | LDT | 0.95375554 |  |  |  |  |  |  |
| VENT | PVH | 0.9535483 |  |  |  |  |  |  |
| LAT | ATN | 0.95350035 |  |  |  |  |  |  |
| LHA | TU | 0.95342703 |  |  |  |  |  |  |
| NLOT | ZI | 0.9529778 |  |  |  |  |  |  |
| VENT | MED | 0.95288927 |  |  |  |  |  |  |
| AAA | RT | 0.95268315 |  |  |  |  |  |  |
| LA | PVi | 0.95255334 |  |  |  |  |  |  |
| AUDpo | VISl | 0.95216683 |  |  |  |  |  |  |
| ATN | PPY | 0.95213251 |  |  |  |  |  |  |
| CA2 | BST | 0.95210774 |  |  |  |  |  |  |
| DMH | VMH | 0.95205786 |  |  |  |  |  |  |
| PL | RAmb | 0.95201317 |  |  |  |  |  |  |
| AAA | TRS | 0.95181913 |  |  |  |  |  |  |
| TRS | ATN | 0.95177257 |  |  |  |  |  |  |
| MTN | PPY | 0.95174492 |  |  |  |  |  |  |
| IA | TU | 0.95172339 |  |  |  |  |  |  |
| BA | ILM | 0.95160261 |  |  |  |  |  |  |
| AAA | GEN<sub>v</sub> | 0.95126826 |  |  |  |  |  |  |
| VENT | AHN | 0.95125246 |  |  |  |  |  |  |
| TRS | MTN | 0.95104375 |  |  |  |  |  |  |
| GEN<sub>v</sub> | PVH | 0.95103017 |  |  |  |  |  |  |
| BLA | BMA | 0.95055784 |  |  |  |  |  |  |
| SBPV | AHN | 0.95029013 |  |  |  |  |  |  |
| AHN | VMH | 0.95010062 |  |  |  |  |  |  |
| CLA | GPe | 0.94999635 |  |  |  |  |  |  |
| ILM | PVH | 0.94999084 |  |  |  |  |  |  |
| CA2 | BMA | 0.94980978 |  |  |  |  |  |  |
| CEA | GPe | 0.94978474 |  |  |  |  |  |  |
| FS | ASO | 0.94977563 |  |  |  |  |  |  |
| CA3 | ZI | 0.94974937 |  |  |  |  |  |  |
| SG | VI | 0.94942935 |  |  |  |  |  |  |
| SPA | PVH | 0.94927656 |  |  |  |  |  |  |
| VENT | TU | 0.94899082 |  |  |  |  |  |  |
| AAA | MA | 0.94897738 |  |  |  |  |  |  |
| NLOT | BST | 0.94895451 |  |  |  |  |  |  |
| MED | GEN<sub>v</sub> | 0.94877833 |  |  |  |  |  |  |
| ATN | ASO | 0.94875195 |  |  |  |  |  |  |
| LA | LHA | 0.94828904 |  |  |  |  |  |  |
| SPA | ILM | 0.94811604 |  |  |  |  |  |  |
| CEA | RCH | 0.94795125 |  |  |  |  |  |  |
| SPA | SFO | 0.94772691 |  |  |  |  |  |  |
| GPI | AHN | 0.94769869 |  |  |  |  |  |  |
| VENT | MTN | 0.94769417 |  |  |  |  |  |  |
| GPI | MED | 0.94768776 |  |  |  |  |  |  |
| VENT | SPF | 0.94679307 |  |  |  |  |  |  |
| SPA | ASO | 0.94665624 |  |  |  |  |  |  |
| AAA | SPF | 0.94653553 |  |  |  |  |  |  |
| MTN | ASO | 0.94625006 |  |  |  |  |  |  |
| AId | TT | 0.94615173 |  |  |  |  |  |  |
| SO | PH | 0.945965 |  |  |  |  |  |  |
| AUDp | VISal | 0.94594586 |  |  |  |  |  |  |
| AUDd | AUDp | 0.94558539 |  |  |  |  |  |  |
| MPN | LHA | 0.94551754 |  |  |  |  |  |  |
| CA3 | SFO | 0.94551285 |  |  |  |  |  |  |
| BLA | RCH | 0.94548918 |  |  |  |  |  |  |
| GPI | PVH | 0.94541751 |  |  |  |  |  |  |
| PVa | PSTN | 0.94533537 |  |  |  |  |  |  |
| VENT | RCH | 0.94491126 |  |  |  |  |  |  |
| FS | MPO | 0.94457886 |  |  |  |  |  |  |
| VISl | RAmb | 0.94456715 |  |  |  |  |  |  |
| CLA | PS | 0.9443564 |  |  |  |  |  |  |
| LA | ZI | 0.94422114 |  |  |  |  |  |  |
| BA | GPI | 0.94383565 |  |  |  |  |  |  |
| BA | SPF | 0.94379759 |  |  |  |  |  |  |
| DMH | AHN | 0.94359867 |  |  |  |  |  |  |
| VENT | ATN | 0.9431805 |  |  |  |  |  |  |
| MPN | LPO | 0.94311174 |  |  |  |  |  |  |
| IG | EP | 0.94298431 |  |  |  |  |  |  |
| VENT | LPO | 0.9427567 |  |  |  |  |  |  |
| PVa | SBPV | 0.9424615 |  |  |  |  |  |  |
| AUD<sub>po</sub> | PL | 0.9422474 |  |  |  |  |  |  |
| VISl | PL | 0.94210523 |  |  |  |  |  |  |
| BLA | ZI | 0.94190109 |  |  |  |  |  |  |
| MTN | PVH | 0.94189211 |  |  |  |  |  |  |
| AAA | PVH | 0.94180615 |  |  |  |  |  |  |
| SPF | SFO | 0.94180593 |  |  |  |  |  |  |
| BST | ZI | 0.94164204 |  |  |  |  |  |  |
| AAA | LHA | 0.94161134 |  |  |  |  |  |  |
| SCH | VMH | 0.941403 |  |  |  |  |  |  |
| DTN | LDT | 0.9413247 |  |  |  |  |  |  |
| PVi | AHN | 0.94123473 |  |  |  |  |  |  |
| B | PCG | 0.94083108 |  |  |  |  |  |  |
| MTN | ZI | 0.9407045 |  |  |  |  |  |  |
| PBG | y | 0.94057951 |  |  |  |  |  |  |
| GPi | SO | 0.94046233 |  |  |  |  |  |  |
| VISpl | RPO | 0.94032944 |  |  |  |  |  |  |
| NLOT | VENT | 0.94024287 |  |  |  |  |  |  |
| GPi | VMH | 0.94015375 |  |  |  |  |  |  |
| ILM | LHA | 0.94014749 |  |  |  |  |  |  |
| IA | SO | 0.94009384 |  |  |  |  |  |  |
| ATN | ZI | 0.94005974 |  |  |  |  |  |  |
| BLA | TRS | 0.93967564 |  |  |  |  |  |  |
| VLPO | LPO | 0.93953464 |  |  |  |  |  |  |
| GPi | BST | 0.93943156 |  |  |  |  |  |  |
| PBG | VTN | 0.939317 |  |  |  |  |  |  |
| CEA | ZI | 0.93914887 |  |  |  |  |  |  |
| PSV | RM | 0.93908893 |  |  |  |  |  |  |
| GPI | ATN | 0.93896587 |  |  |  |  |  |  |
| FS | PVH | 0.93864564 |  |  |  |  |  |  |
| CEA | IA | 0.93859169 |  |  |  |  |  |  |
| GEN<sub>v</sub> | SO | 0.93854332 |  |  |  |  |  |  |
| FS | BA | 0.93843587 |  |  |  |  |  |  |
| NLOT | GPI | 0.93837297 |  |  |  |  |  |  |
| BA | SPA | 0.93834738 |  |  |  |  |  |  |
| CA3 | RT | 0.93785128 |  |  |  |  |  |  |
| BA | STN | 0.93765381 |  |  |  |  |  |  |
| IA | LPO | 0.93741348 |  |  |  |  |  |  |
| CA3 | ILM | 0.93735763 |  |  |  |  |  |  |
| LA | TU | 0.93725814 |  |  |  |  |  |  |
| NLOT | BMA | 0.93709655 |  |  |  |  |  |  |
| GPI | MTN | 0.9369888 |  |  |  |  |  |  |
| LAT | MTN | 0.93690998 |  |  |  |  |  |  |
| SFO | AHN | 0.93685218 |  |  |  |  |  |  |
| SPA | MPN | 0.93640551 |  |  |  |  |  |  |
| SSp | ATN | 0.93628147 |  |  |  |  |  |  |
| LHA | LPO | 0.93619244 |  |  |  |  |  |  |
| AUD<sub>p</sub> | AUD<sub>v</sub> | 0.9361108 |  |  |  |  |  |  |
| SPF | LPO | 0.93598856 |  |  |  |  |  |  |
| NLOT | AAA | 0.93593061 |  |  |  |  |  |  |
| LA | GEN<sub>v</sub> | 0.93568661 |  |  |  |  |  |  |
| IG | SF | 0.93561799 |  |  |  |  |  |  |
| FC | PAS | 0.93541275 |  |  |  |  |  |  |
| RT | SO | 0.93540938 |  |  |  |  |  |  |
| BA | ATN | 0.93520006 |  |  |  |  |  |  |
| ASO | PVH | 0.93518927 |  |  |  |  |  |  |
| LPO | ZI | 0.93501205 |  |  |  |  |  |  |
| PVH | ZI | 0.93486845 |  |  |  |  |  |  |
| GEN<sub>v</sub> | LPO | 0.93485454 |  |  |  |  |  |  |
| CA3 | ATN | 0.93483265 |  |  |  |  |  |  |
| SPA | VLPO | 0.93477788 |  |  |  |  |  |  |
| CA2 | LA | 0.93438468 |  |  |  |  |  |  |
| AUD<sub>v</sub> | ENT | 0.93417613 |  |  |  |  |  |  |
| AIp | NLOT | 0.93402644 |  |  |  |  |  |  |
| LA | VENT | 0.93369715 |  |  |  |  |  |  |
| CA3 | GEN<sub>v</sub> | 0.93358751 |  |  |  |  |  |  |
| SPF | LHA | 0.93354676 |  |  |  |  |  |  |
| CEA | PVH | 0.93347423 |  |  |  |  |  |  |
| ASO | LPO | 0.93328741 |  |  |  |  |  |  |
| ATN | PVH | 0.93306051 |  |  |  |  |  |  |
| VENT | SPA | 0.93272476 |  |  |  |  |  |  |
| ARH | SCH | 0.93271344 |  |  |  |  |  |  |
| ACVII | LRN | 0.93231827 |  |  |  |  |  |  |
| CA3 | IA | 0.93230107 |  |  |  |  |  |  |
| LA | BLA | 0.9321628 |  |  |  |  |  |  |
| CEA | LPO | 0.93214947 |  |  |  |  |  |  |
| LA | SBPV | 0.93213559 |  |  |  |  |  |  |
| GPI | LPO | 0.93189661 |  |  |  |  |  |  |
| BA | ZI | 0.93183444 |  |  |  |  |  |  |
| BLA | GPI | 0.93174762 |  |  |  |  |  |  |
| SO | AHN | 0.93168775 |  |  |  |  |  |  |
| CEA | PS | 0.93145403 |  |  |  |  |  |  |
| FS | ILM | 0.93140978 |  |  |  |  |  |  |
| PV<sub>a</sub> | VMH | 0.93140772 |  |  |  |  |  |  |
| MPO | VLPO | 0.93136452 |  |  |  |  |  |  |
| COA | PAA | 0.93131347 |  |  |  |  |  |  |
| LA | AHN | 0.93126611 |  |  |  |  |  |  |
| AHN | LPO | 0.9310919 |  |  |  |  |  |  |
| LAT | MED | 0.9310646 |  |  |  |  |  |  |
| TRS | LHA | 0.9310111 |  |  |  |  |  |  |
| TRS | TU | 0.93099284 |  |  |  |  |  |  |
| SO | ZI | 0.93098661 |  |  |  |  |  |  |
| HEM | PVi | 0.93087907 |  |  |  |  |  |  |
| SO | LHA | 0.93051935 |  |  |  |  |  |  |
| BLA | SF | 0.93041925 |  |  |  |  |  |  |
| MA | MPN | 0.93027973 |  |  |  |  |  |  |
| SPF | ILM | 0.93024165 |  |  |  |  |  |  |
| BST | VENT | 0.93017928 |  |  |  |  |  |  |
| BST | SPF | 0.93014724 |  |  |  |  |  |  |
| MA | SO | 0.93013266 |  |  |  |  |  |  |
| CA3 | SPF | 0.9300586 |  |  |  |  |  |  |
| PVH | PH | 0.92999176 |  |  |  |  |  |  |
| MTN | LPO | 0.92993105 |  |  |  |  |  |  |
| CEA | TU | 0.92979948 |  |  |  |  |  |  |
| ORBl | IG | -0.9309675 |  |  |  |  |  |  |
| IG | ENT | -0.9313086 |  |  |  |  |  |  |
| TEa | IG | -0.932185 |  |  |  |  |  |  |
| TT | TU | -0.9326137 |  |  |  |  |  |  |
| B | IO | -0.9330159 |  |  |  |  |  |  |
| AId | LA | -0.9338227 |  |  |  |  |  |  |
| VISl | ACB | -0.933958 |  |  |  |  |  |  |
| TT | RCH | -0.9356735 |  |  |  |  |  |  |
| PRE | PA | -0.9372874 |  |  |  |  |  |  |
| PRE | VMH | -0.9379279 |  |  |  |  |  |  |
| ORBl | SF | -0.9411831 |  |  |  |  |  |  |
| TT | BMA | -0.9411929 |  |  |  |  |  |  |
| AUDpo | LS | -0.9433956 |  |  |  |  |  |  |
| PRE | RCH | -0.9444777 |  |  |  |  |  |  |
| ORBm | SF | -0.9455869 |  |  |  |  |  |  |
| AOB | SPF | -0.9474364 |  |  |  |  |  |  |
| PRE | BST | -0.949783 |  |  |  |  |  |  |
| TEa | EP | -0.9497843 |  |  |  |  |  |  |
| ORBm | CP | -0.9506136 |  |  |  |  |  |  |
| PL | LS | -0.9512804 |  |  |  |  |  |  |
| ENT | EP | -0.9522718 |  |  |  |  |  |  |
| DP | BLA | -0.9606578 |  |  |  |  |  |  |
| AUDp | LS | -0.9614194 |  |  |  |  |  |  |
| ORBl | CA2 | -0.9640302 |  |  |  |  |  |  |
| TT | CA2 | -0.9653638 |  |  |  |  |  |  |
| TT | BLA | -0.9756476 |  |  |  |  |  |  |

**Table 6:** Sensitivity analysis. Results from permutation testing (n = 1,000 permutations) pairwise group-level differences in network density, using different thresholding methods to derive pup call response networks from *cFos*^*PC*^ correlation matrices.

| Naïve virgins vs. Experienced virgins |  |  | Naïve virgins vs. Mothers |  |  | Experienced virgins vs. Mothers |  |  |
| --- | --- | --- | --- | --- | --- | --- | --- | --- |
| *Edge thresh.* | *Obs. $\Delta$ network density* | *p-value* | *Edge thresh.* | *Obs. $\Delta$ network density* | *p-value* | *Edge thresh.* | *Obs. $\Delta$ network density* | *p-value* |
| p < 0.05 | 0.028429 | 0.106 (ns) | p < 0.05 | 0.048685 | 0.019 (*) | p < 0.05 | 0.020256 | 0.128 (ns) |
| p < 0.01 | 0.018735 | 0.069 (ns) | p < 0.01 | 0.028689 | 0.021 (*) | p < 0.01 | 0.0099544 | 0.17 (ns) |
| p < 0.001 | 0.0069115 | 0.136 (ns) | p < 0.001 | 0.010433 | 0.059 (ns) | p < 0.001 | 0.003521 | 0.226 (ns) |
| FDR-BH ($\alpha = 0.05$) | 0.013128 | 0.095 (ns) | FDR-BH ($\alpha = 0.05$) | 0.016257 | 0.037 (*) | FDR-BH ($\alpha = 0.05$) | 0.0031298 | 0.307 (ns) |
| FDR-BKY ($\alpha = 0.05$) | 0.012997 | 0.094 (ns) | FDR-BKY ($\alpha = 0.05$) | 0.01591 | 0.038 (*) | FDR-BKY ($\alpha = 0.05$) | 0.0029124 | 0.309 (ns) |

To understand which brain regions may play a key role in coordinating each group’s brain-wide pup call response network, we extracted the top 10% of connected nodes, representing the hub regions in each group’s pup call response network (**Figure 5A, Extended Figure 11A-B, Table 4**). We then compared the sets of hub regions between groups (**Figure 5A, Extended Figure 11B**). This exploratory analysis revealed that there is minimal overlap in hub regions between groups, aside from four fully overlapping regions: the central amygdala (CEA), intercalated amygdalar nucleus (IA), lateral hypothalamic area (LHA), and supraoptic nucleus (SO) (**Figure 5A-B**). Other hub regions stand out as being shared between virgin groups (naïve and experienced virgins), or between maternally experienced groups (experienced virgins and mothers) (**Figure 5B**).

**Figure 5:**
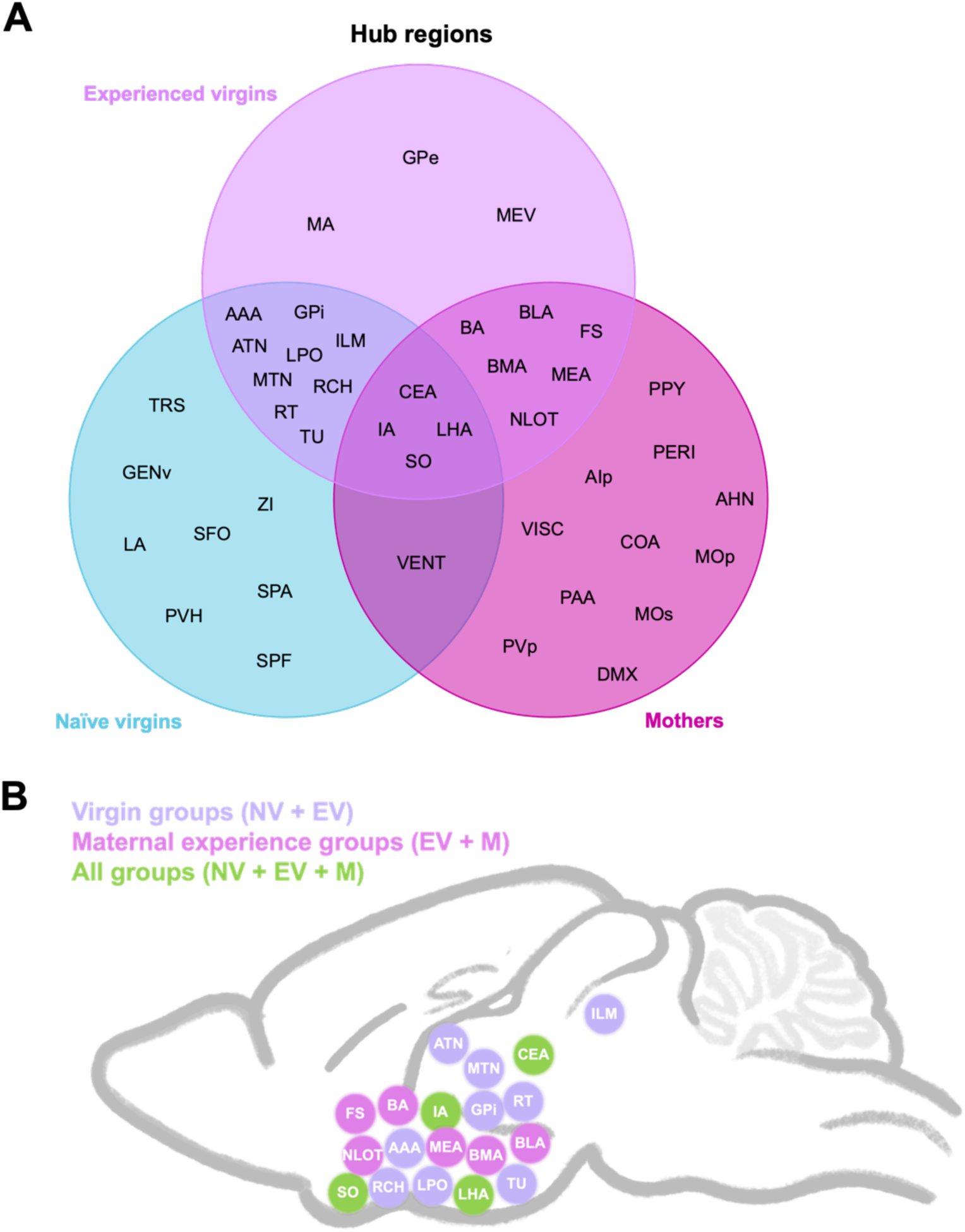
Pup call response hub regions across groups. (A) Venn diagram depicting abbreviations of unique and overlapping hub regions (hubs defined as top 10% most connected nodes per pup call response network) between groups (see Table 1 for region abbreviations). (B) Schematic of pup call response hub regions (purple = hub shared across virgin groups, naïve and experienced virgins; pink = hub shared across maternally experienced groups, experienced virgins and mothers; green = hub shared across all 3 groups).

Overall, our pup call response network analyses revealed that pup calls recruit different networks in naïve virgins, experienced virgins, and mothers. Furthermore, mothers recruit a pup call response network that is sparser, yet stronger, than that of naïve virgins, with experienced virgins appearing to show an intermediate effect of maternal experience.

### Pup call responses in OXT projection-dense regions

Prior work revealed that maternal plasticity in left A1 is facilitated by OXT, which acts by rebalancing excitatory and inhibitory responses to pup calls to sensitize A1 to this behaviorally-relevant stimulus (Carcea et al., 2021; Marlin et al., 2015). OXT+ cells in the paraventricular nucleus of the hypothalamus (PVH) have been shown to respond to pup calls, receiving inputs from a non-canonical auditory pathway, allowing for pup call-evoked OXT release throughout the brain (Valtcheva et al., 2023). Furthermore, previous studies have shown that expression of the OXT receptor is particularly prevalent in inhibitory interneurons (K. Li et al., 2016; Marlin et al., 2015; Mitre et al., 2016; Nakajima et al., 2014; Schimmer et al., 2026), perhaps explaining why exogenous OXT injection was shown to have an overall inhibitory effect on brain activity (Choe et al., 2022). With this in mind, we hypothesized that pup calls recruit a sparser network in mothers due to brain-wide inhibition via OXT signaling.

To probe whether OXT may be playing the same sensitization role in the whole brain as it does in A1, we mapped OXT projections across the entire brain in naïve virgins, experienced virgins, and mothers. Using an anti-OXT antibody, we stained for OXT+ cells and their projections in the same samples that underwent iDISCO+ clearing, c-Fos staining, and light-sheet imaging (**Figure 6A**, **Supplemental Video 11**) (Renier et al., 2014). We then built upon published methods (Friedmann et al., 2020; Gongwer et al., 2023) to develop a deep learning-based pipeline to automatically segment OXT projections, identify their coordinates within the sample, and align the sample to the Allen Brain Atlas – applying the same Elastix transformations generated from the ClearMap pipeline – to assign each OXT+ voxel to one of our 215 brain regions (**Figure 6A**). We focused our analyses on the left hemisphere, as prior studies have tested and found no lateralization of OXT projections (Marlin et al., 2015; Mitre et al., 2016). We modeled OXT+ voxel counts per region using a GLMM (see **Methods**) (Zimmerman et al., 2024). This analysis yielded 16 regions for which the inclusion of group information provided better explanatory power, but no regions passed correction for multiple comparisons (**Table 7**). Again, compared to prior work using GLMMs, we were poorly powered, perhaps limiting the conclusions that can be drawn from this analysis regarding differences in regional OXT expression. Furthermore, our light-sheet imaging parameters were primarily optimized for c-Fos expression, thus we cannot rule out the possibility that increased resolution may reveal more nuanced insights regarding OXT projections. Nevertheless, through this analysis we found no significant regional differences in OXT projection density between naïve virgins, experienced virgins, and mothers. Given that we found no group differences, we averaged relative OXT projection density (see **Methods)** per region across all animals (**Table 8**, **Supplemental Video 12**) and extracted the top 10% of OXT projection-dense regions. This allowed us to establish an OXT projection-dense circuit that is common across all groups studied (**Figure 6B**).

**Figure 6:**
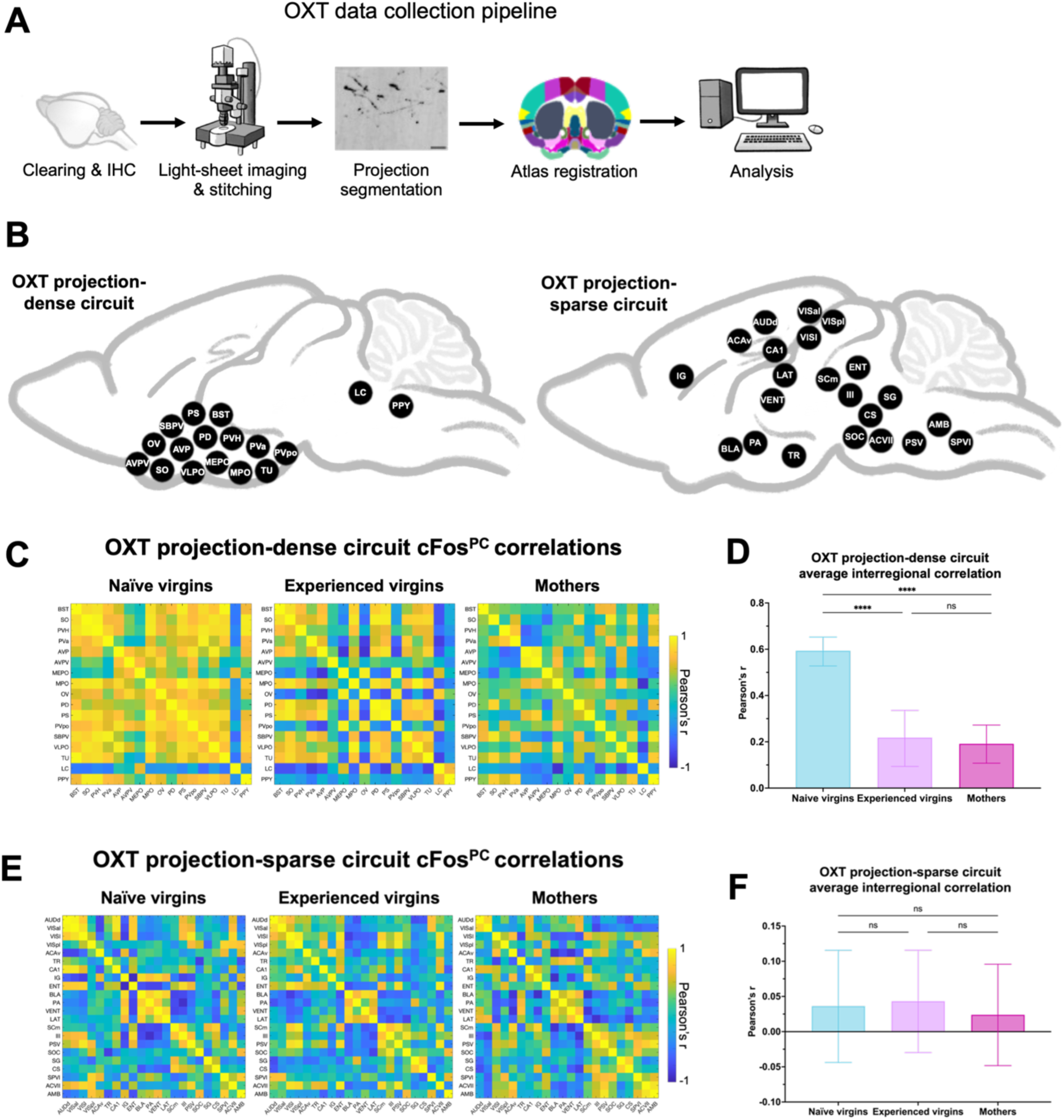
OXT projection-dense circuit reveals experience-dependent synchrony upon pup call exposure between groups. (A) OXT data collection pipeline, which included iDISCO+ clearing, anti-OXT immunohistochemistry (IHC), light-sheet imaging and tile stitching, OXT projection segmentation via our custom segmentation model, Allen Brain Atlas registration, and analysis (see Methods for details). Scale bar in representative image represents 25µm. (B) OXT projection-dense circuit (see Table 1 for abbreviations). (C) Pearson correlation matrices indicating interregional correlations in OXT projection-dense circuit, based on ***cFos***^***PC***^in pup call-exposed samples across groups (n = 8 x 17 regions). Matrix pixel colors represent Pearson correlation coefficients, see scale to the right. (D) Mean interregional correlation among OXT projection-dense circuit regions per group. Kruskal-Wallis test following Fisher r-to-z transformation revealed significant group effect (p < 0.0001). Comparisons on graph represent Tukey’s post-hoc tests (naïve virgins vs. experienced virgins: p < 0.0001; naïve virgins vs. mothers: p < 0.0001; experienced virgins vs. mothers: p > 0.9999; n = 8 x 17 regions). Error bars denote mean ± 95% CI. **** p < 0.0001.

**Table 7:** GLMM results. Results from fitting a full and reduced GLMM (see Methods) per brain region, and comparing the fit of the 2 models. Significant p-value would indicate that full GLMM explains the OXT+ voxel counts in that brain region better than the reduced GLMM. Middle column indicates p-value of model comparison (green shading highlights regions with p < 0.05), rightmost column indicates adjusted p-value of model comparison (red shading highlights regions with FDR < 0.10).

| Region | p-value from model comparison | Adjusted p-value (FDR < 0.10) |
| --- | --- | --- |
| FRP | 0.88463197 | 0.97038711 |
| MOp | 0.53115262 | 0.87010215 |
| MOs | 0.81183505 | 0.96744583 |
| SSp | 0.2650033 | 0.76072212 |
| SSs | 0.04433771 | 0.59578798 |
| ILA | 0.4453256 | 0.85534112 |
| GU | 0.55649087 | 0.87010215 |
| VISC | 0.85969596 | 0.96744583 |
| AUDd | 0.69844534 | 0.92845449 |
| AUDp | 0.12339947 | 0.66128526 |
| AUDpo | 0.03574206 | 0.59578798 |
| AUDv | 0.12616597 | 0.66128526 |
| VISal | 0.00352761 | 0.37921769 |
| VISam | 0.70160869 | 0.92845449 |
| VISl | 0.11146917 | 0.66128526 |
| VISp | 0.16660817 | 0.68886069 |
| VISpl | 0.0533417 | 0.64336192 |
| VISpm | 0.77079325 | 0.95125265 |
| ACAd | 0.45672744 | 0.86444671 |
| ACAv | 0.98997561 | 0.99927115 |
| PL | 0.49329965 | 0.87010215 |
| ORBl | 0.31788528 | 0.82035051 |
| ORBm | 0.0913891 | 0.66128526 |
| ORBvl | 0.15352374 | 0.66255375 |
| AId | 0.62882562 | 0.88945729 |
| AIp | 0.53248062 | 0.87010215 |
| AIv | 0.49464937 | 0.87010215 |
| RSPagl | 0.47604108 | 0.87010215 |
| RSPd | 0.32665885 | 0.82035051 |
| RSPv | 0.21298983 | 0.74607731 |
| TEa | 0.10662704 | 0.66128526 |
| PERI | 0.49496162 | 0.87010215 |
| ECT | 0.22325802 | 0.74607731 |
| MOB | 0.24898863 | 0.74607731 |
| AOB | 0.00340371 | 0.37921769 |
| AON | 0.18805144 | 0.7219832 |
| TT | 0.24984915 | 0.74607731 |
| DP | 0.85519355 | 0.96744583 |
| PIR | 0.12653861 | 0.66128526 |
| NLOT | 0.36944157 | 0.82035051 |
| COA | 0.35984535 | 0.82035051 |
| PAA | 0.09023485 | 0.66128526 |
| TR | 0.00861251 | 0.43152577 |
| CA1 | 0.50104694 | 0.87010215 |
| CA2 | 0.87745087 | 0.96744583 |
| CA3 | 0.72545167 | 0.93475934 |
| DG | 0.73246751 | 0.93475934 |
| FC | 0.0672615 | 0.66128526 |
| IG | 0.09625729 | 0.66128526 |
| ENT | 0.02988062 | 0.59578798 |
| PAR | 0.34888389 | 0.82035051 |
| POST | 0.76776163 | 0.95125265 |
| PRE | 0.03322533 | 0.59578798 |
| SUB | 0.31521928 | 0.82035051 |
| CLA | 0.33971307 | 0.82035051 |
| EP | 0.49026998 | 0.87010215 |
| LA | 0.68194001 | 0.92795635 |
| BLA | 0.60300102 | 0.87010215 |
| BMA | 0.90291996 | 0.97718426 |
| PA | 0.46685123 | 0.87010215 |
| CP | 0.12252117 | 0.66128526 |
| ACB | 0.50943987 | 0.87010215 |
| FS | 0.75435714 | 0.94846073 |
| OT | 0.65055124 | 0.9006051 |
| LS | 0.41367077 | 0.83007681 |
| SF | 0.04185403 | 0.59578798 |
| SH | 0.10662301 | 0.66128526 |
| AAA | 0.69770583 | 0.92845449 |
| BA | 0.30438002 | 0.82035051 |
| CEA | 0.24762345 | 0.74607731 |
| IA | 0.71528073 | 0.93203247 |
| MEA | 0.06861947 | 0.66128526 |
| GPe | 0.40512507 | 0.82415358 |
| GPi | 0.40632688 | 0.82415358 |
| SI | 0.03476809 | 0.59578798 |
| MA | 0.14784291 | 0.66255375 |
| MSC | 0.37011162 | 0.82035051 |
| TRS | 0.01003548 | 0.43152577 |
| BST | 0.06883298 | 0.66128526 |
| BAC | 0.5611744 | 0.87010215 |
| VERM | 0.13278187 | 0.66128526 |
| HEM | 0.55995953 | 0.87010215 |
| FN | 0.53868818 | 0.87010215 |
| IP | 0.79261735 | 0.95971508 |
| DN | 0.74709564 | 0.94485625 |
| VENT | 0.15008326 | 0.66255375 |
| SPF | 0.09388679 | 0.66128526 |
| SPA | 0.90900861 | 0.97718426 |
| PP | 0.54576201 | 0.87010215 |
| GENd | 0.96253946 | 0.99493261 |
| LAT | 0.99974091 | 0.99974091 |
| ATN | 0.98296291 | 0.99927115 |
| MED | 0.66490262 | 0.91053543 |
| MTN | 0.43705281 | 0.85423959 |
| ILM | 0.94916273 | 0.99493261 |
| RT | 0.73216501 | 0.93475934 |
| GENv | 0.99480946 | 0.9994581 |
| EPI | 0.22405525 | 0.74607731 |
| SO | 0.84460907 | 0.96744583 |
| ASO | 0.15939555 | 0.67196165 |
| PVH | 0.12608408 | 0.66128526 |
| PVa | 0.09779936 | 0.66128526 |
| PVi | 0.08060429 | 0.66128526 |
| ARH | 0.61594702 | 0.8828574 |
| ADP | 0.23489165 | 0.74607731 |
| AVP | 0.95610281 | 0.99493261 |
| AVPV | 0.64613085 | 0.9006051 |
| DMH | 0.5043899 | 0.87010215 |
| MEPO | 0.56269805 | 0.87010215 |
| MPO | 0.40397505 | 0.82415358 |
| OV | 0.27961256 | 0.78529472 |
| PD | 0.58146625 | 0.87010215 |
| PS | 0.23557755 | 0.74607731 |
| PVp | 0.77254559 | 0.95125265 |
| PVpo | 0.11979533 | 0.66128526 |
| SBPV | 0.35282531 | 0.82035051 |
| SCH | 0.3598652 | 0.82035051 |
| SFO | 0.31286682 | 0.82035051 |
| VLPO | 0.98787803 | 0.99927115 |
| AHN | 0.39403858 | 0.82415358 |
| MBO | 0.54711792 | 0.87010215 |
| MPN | 0.29779322 | 0.82035051 |
| PMd | 0.24044945 | 0.74607731 |
| PMv | 0.62448619 | 0.88916908 |
| PVHd | 0.52609149 | 0.87010215 |
| VMH | 0.33280264 | 0.82035051 |
| PH | 0.59693365 | 0.87010215 |
| LHA | 0.82934272 | 0.96744583 |
| LPO | 0.09963081 | 0.66128526 |
| PST | 0.58075761 | 0.87010215 |
| PSTN | 0.39745404 | 0.82415358 |
| RCH | 0.90869235 | 0.97718426 |
| STN | 0.96077852 | 0.99493261 |
| TU | 0.9541251 | 0.99493261 |
| ZI | 0.40015679 | 0.82415358 |
| SCs | 0.48460134 | 0.87010215 |
| IC | 0.31542643 | 0.82035051 |
| NB | 0.25535839 | 0.75208292 |
| SAG | 0.00647539 | 0.43152577 |
| PBG | 0.94770284 | 0.99493261 |
| MEV | 0.05572317 | 0.64336192 |
| SNr | 0.45835779 | 0.86444671 |
| VTA | 0.58798278 | 0.87010215 |
| RR | 0.35413132 | 0.82035051 |
| MRN | 0.79455481 | 0.95971508 |
| SCm | 0.41696882 | 0.83007681 |
| PAG | 0.85212583 | 0.96744583 |
| PRT | 0.35594239 | 0.82035051 |
| CUN | 0.97007192 | 0.99792088 |
| RN | 0.24852848 | 0.74607731 |
| III | 0.14544031 | 0.66255375 |
| EW | 0.36791604 | 0.82035051 |
| IV | 0.22736713 | 0.74607731 |
| VTN | 0.82409942 | 0.96744583 |
| AT | 0.65346231 | 0.9006051 |
| LT | 0.87319686 | 0.96744583 |
| SNC | 0.24531827 | 0.74607731 |
| PPN | 0.53083842 | 0.87010215 |
| RAmb | 0.38713104 | 0.82415358 |
| NLL | 0.34019588 | 0.82035051 |
| PSV | 0.93564126 | 0.99493261 |
| PB | 0.22060215 | 0.74607731 |
| SOC | 0.86780225 | 0.96744583 |
| B | 0.01722509 | 0.52905632 |
| DTN | 0.13217395 | 0.66128526 |
| PCG | 0.03930586 | 0.59578798 |
| PG | 0.12758925 | 0.66128526 |
| PRNc | 0.13184129 | 0.66128526 |
| SG | 0.04068224 | 0.59578798 |
| SUT | 0.5950596 | 0.87010215 |
| TRN | 0.1353328 | 0.66128526 |
| V | 0.58817856 | 0.87010215 |
| CS | 0.86286482 | 0.96744583 |
| LC | 0.18213445 | 0.71198013 |
| LDT | 0.03886348 | 0.59578798 |
| NI | 0.42729406 | 0.84282774 |
| PRNr | 0.21500252 | 0.74607731 |
| RPO | 0.60003824 | 0.87010215 |
| SLC | 0.51081719 | 0.87010215 |
| SLD | 0.15408227 | 0.66255375 |
| AP | 0.52741678 | 0.87010215 |
| CN | 0.65278955 | 0.9006051 |
| DCN | 0.3998506 | 0.82415358 |
| ECU | 0.57344843 | 0.87010215 |
| NTB | 0.32171754 | 0.82035051 |
| NTS | 0.77869984 | 0.95125265 |
| SPVC | 0.77833661 | 0.95125265 |
| SPVI | 0.0143999 | 0.51599624 |
| SPVO | 0.05685524 | 0.64336192 |
| VI | 0.28124508 | 0.78529472 |
| VII | 0.17054689 | 0.69184115 |
| ACVII | 0.83889517 | 0.96744583 |
| AMB | 0.19617655 | 0.73996418 |
| DMX | 0.15155161 | 0.66255375 |
| GRN | 0.37873254 | 0.82415358 |
| ICB | 0.86971068 | 0.96744583 |
| IO | 0.89653114 | 0.97718426 |
| IRN | 0.71174169 | 0.93203247 |
| ISN | 0.69543189 | 0.92845449 |
| LIN | 0.26536818 | 0.76072212 |
| LRN | 0.98757939 | 0.99927115 |
| MARN | 0.81311542 | 0.96744583 |
| MDRN | 0.86548333 | 0.96744583 |
| PARN | 0.70389806 | 0.92845449 |
| PAS | 0.09206406 | 0.66128526 |
| PGRN | 0.53391791 | 0.87010215 |
| PHY | 0.86486487 | 0.96744583 |
| PPY | 0.568148 | 0.87010215 |
| VNC | 0.20864546 | 0.74607731 |
| x | 0.91988361 | 0.9839551 |
| XII | 0.44557305 | 0.85534112 |
| y | 0.54755158 | 0.87010215 |
| RM | 0.17989375 | 0.71198013 |
| RPA | 0.82168412 | 0.96744583 |
| RO | 0.73476432 | 0.93475934 |

**Table 8:**
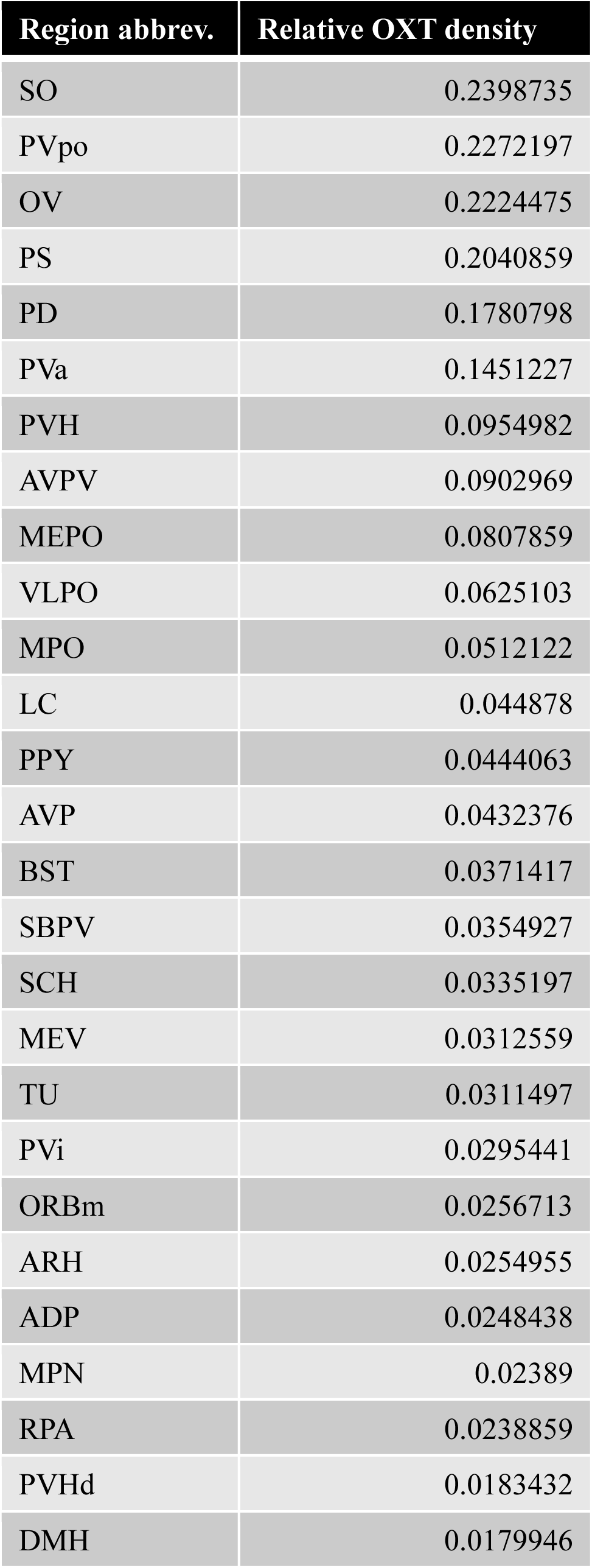
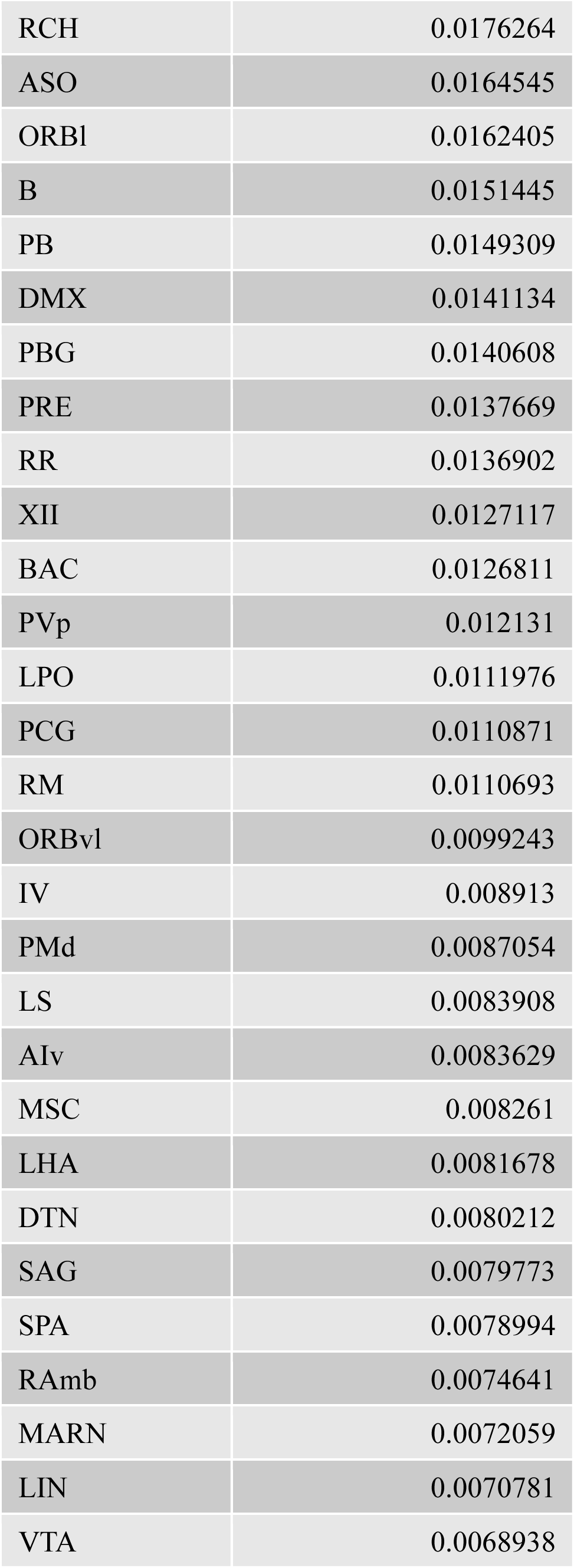
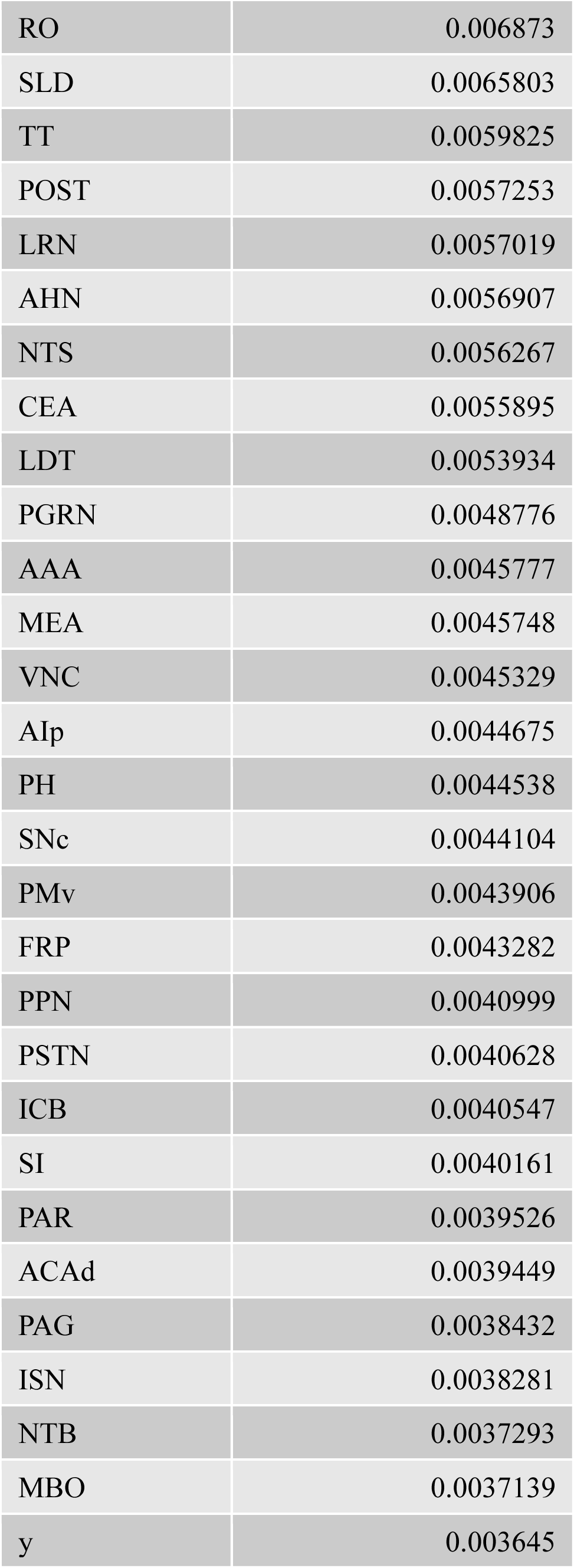
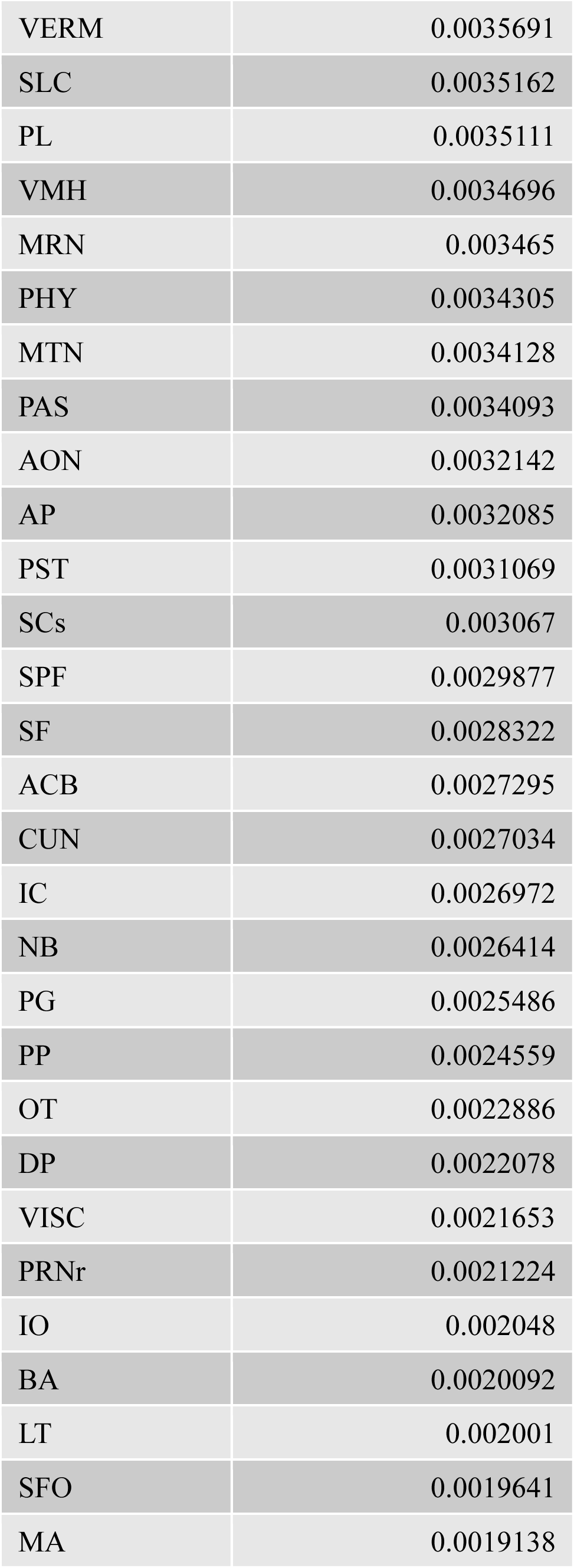
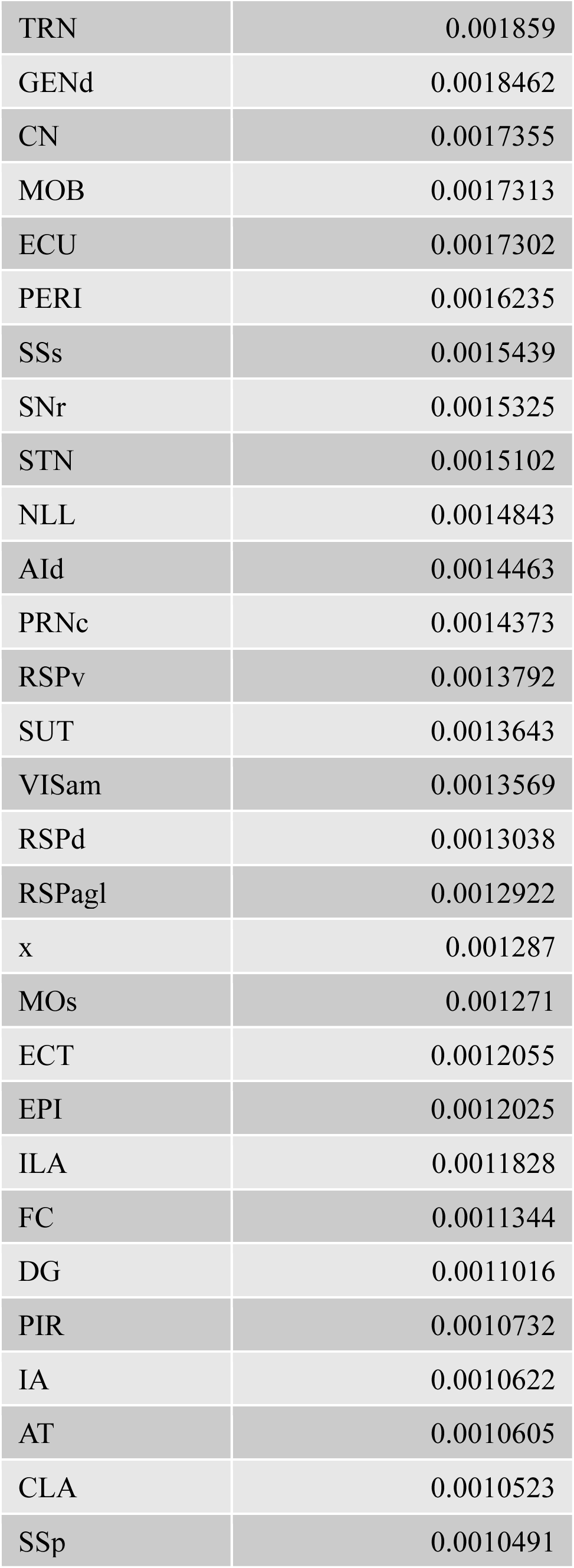
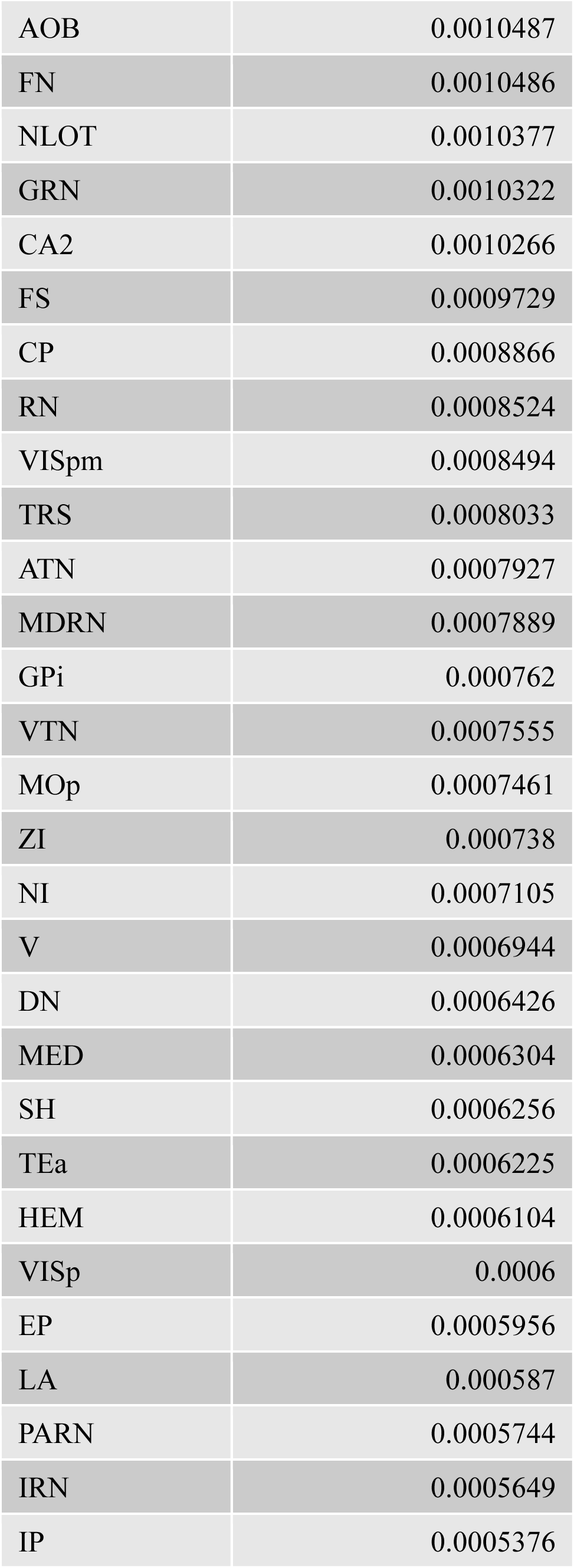
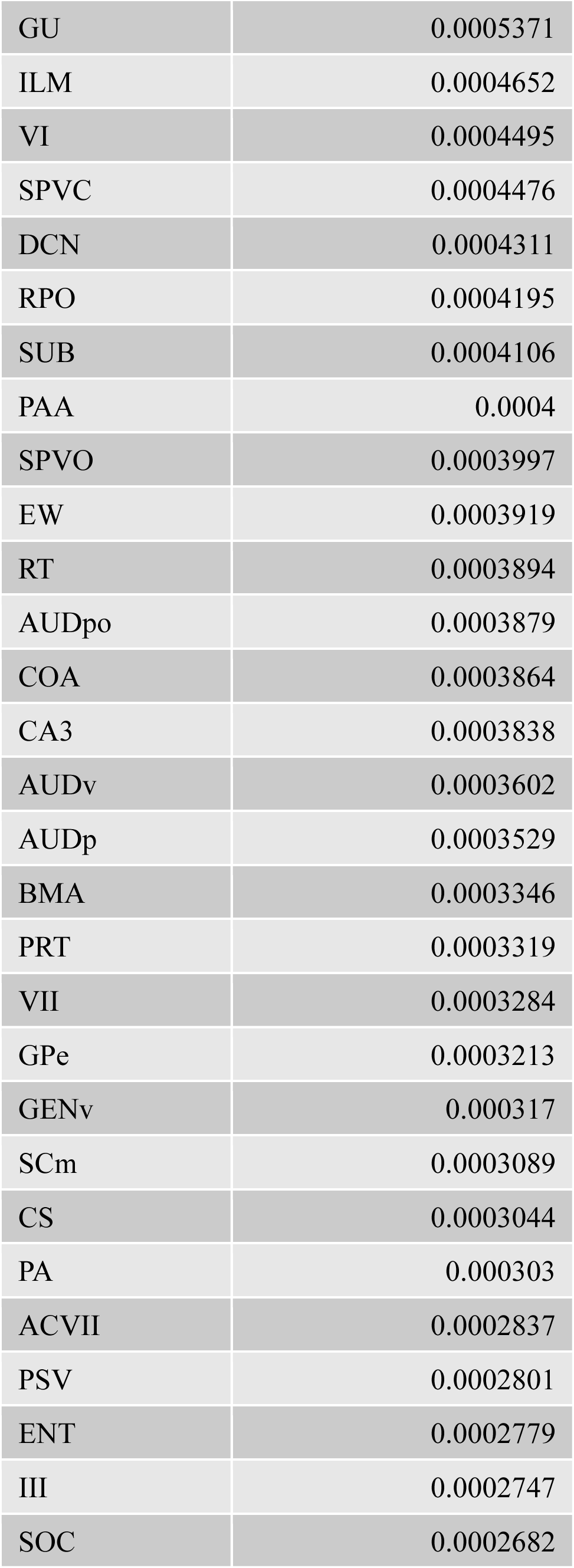
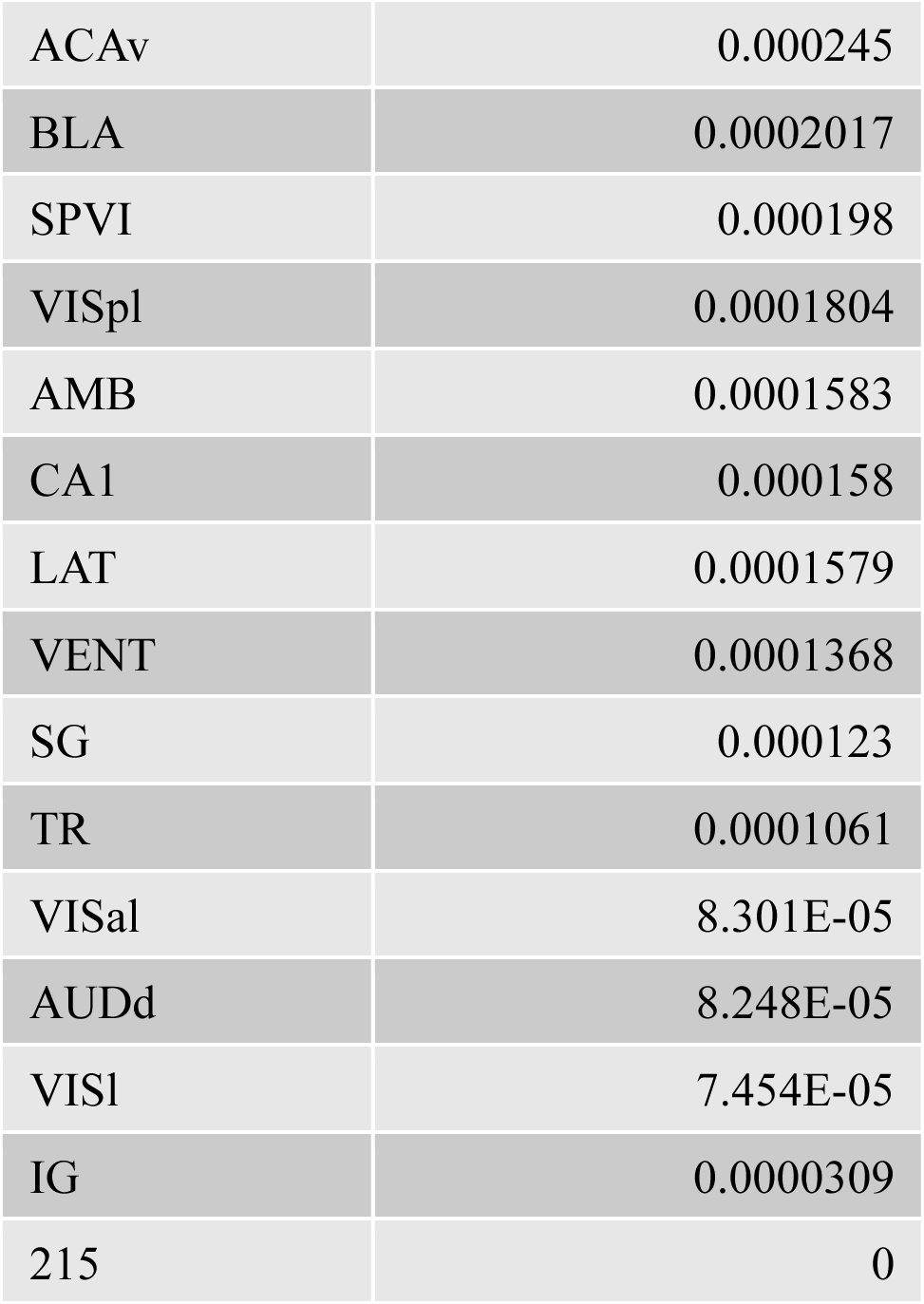
Relative OXT projection density per brain region, per group. For each of the 215 brain regions conserved from Allen Brain Atlas, this table lists their abbreviation (see Table 1 for abbreviations) and relative OXT projection density (see Methods) per group, in descending degree order.

To assess coordinated pup call-evoked responses in regions that receive direct OXT input, we correlated *cFos*^*PC*^ across the OXT projection-dense circuit per group and calculated the average interregional correlation (**Figure 6C-D**). We found a significant group effect that suggested that our two maternally experienced groups recruited this OXT-dense circuit similarly, mirroring our findings in the whole brain. Specifically, experienced virgins and mothers showed lower synchrony in this circuit, compared to naïve virgins, perhaps indicating that inhibitory signaling was at play in these regions during pup call perception in experienced mice. Notably, this group effect was not present when we correlated *cFos*^*PC*^ across the OXT projection-sparse circuit (**Figure 6B, E-F**), suggesting that OXT signaling may drive the experience-dependent group differences we observed at the level of the whole brain.

## Discussion

Prior work has revealed that maternal care can be unlocked in different ways: through innate processes in the case of mothers, or through observational learning in the case of experienced virgins. Both experienced virgins and mothers ultimately prove to be successful pup retrievers and to have robust, reliable neural responses to pup calls in left A1 (Marlin et al., 2015). However, we hypothesized that there may be nuanced differences in their pup call responses, especially if we extend our focus beyond left A1 to the entire brain. Our findings demonstrate that the experience of providing maternal care leads to previously unidentified changes in pup call-evoked behavioral and neural responses.

By designing and conducting a two-choice behavioral assay that solely presents auditory stimuli, in the absence of other multisensory pup cues, we found surprising differences between our two maternally experienced groups. As expected based on prior literature (Ehret, 1987; Ehret et al., 1987; Lin et al., 2013a; Schiavo et al., 2020; Sewell, 1970; Tasaka et al., 2020), mothers exhibited a preference for investigating a speaker playing pup calls, compared to a speaker playing white noise, while naïve virgins did not. However, experienced virgins also did not show a preference toward pup calls. This result conflicted with prior expectation due to the fact that both experienced virgins and mothers retrieve pups and show strong A1 pup call responses.

While both experienced virgins and mothers help pups in need of care, their difference in pup call preference suggests they may perform maternal behavior for different underlying motivations. It is also possible that experienced virgins require multisensory cues to respond appropriately to pup calls, such as the pup odors that were incorporated into prior studies of their pup call preference (Ehret et al., 1987; Lin et al., 2013b; Schiavo et al., 2020). This could be explained by the phenomenon that coincident pup odor presentation amplifies pup call responses in A1 in mice with maternal experience (Cohen et al., 2011; Cohen & Mizrahi, 2015).

To uncover potential explanations regarding this unexpected difference in pup call-evoked behavior, we took advantage of whole-brain activity mapping via c-Fos to explore how regions across the brain are cooperatively recruited during pup call perception. One way we approached analyzing this data was to start with a priori circuits of interest, namely (1) the Pup-Directed Avoidance and Aggression Circuit and (2) the Parental Care Circuit (Dulac et al., 2014). In comparing the coordinated recruitment of the Pup Avoidance Circuit during pup call exposure, we found that naïve virgins, experienced virgins, and mothers were relatively similar. This result was unsurprising, as none of these groups showed an aversion to pup calls in the two-choice assay, and none of the mice in this study were aggressive towards pups during the pup retrieval assay. In comparing the coordinated recruitment of the Parental Care Circuit during pup call exposure, however, we found more striking differences between groups. The pattern of pup call-evoked correlation was most similar between naïve virgins and experienced virgins, and the least similar between naïve virgins and mothers. Unlike the Pup Avoidance Circuit, the Parental Care Circuit includes higher-order processing areas related to emotional valence, reward, arousal, and decision-making. Therefore, this group difference in c-Fos correlation pattern in the higher-order processing of pup calls may facilitate mothers’ unique pup call preference. Being attracted to pup calls would serve to support adaptive maternal care, especially pup retrieval. It may even confer resiliency, allowing mothers to still be motivated to investigate the source of pup calls even in uncertain or risky environments. In fact, recent work has shown that activity in the prelimbic (PL) and infralimbic (ILA) areas, both part of the Parental Care Circuit, is crucial to successful pup retrieval under threatening conditions in both experienced virgins and mothers (Wu et al., 2025).

Expanding our focus to the entire brain, we found that pup call exposure led to experience-dependent changes in brain-wide activity. Specifically, we found that pup calls, compared to baseline, elicited an increase in average interregional correlation in naïve virgins, while in the two maternally experienced groups, pup calls led to a decrease. Average interregional correlation can be interpreted as a measure of global coupling, or brain-wide synchrony. While at first it may be perceived as counterintuitive, the concept that pup calls shift the experienced brain into a state of lower synchrony may suggest that the maternally experienced brain is more prepared to process this stimulus with less energetic demand. This idea would align with prior work in the field of the Neural Efficiency Hypothesis, which suggests that with expertise comes more efficient neural processing during a task (Bassett & Bullmore, 2017; Haier et al., 1988; L. Li & Smith, 2021; Neubauer & Fink, 2009). This notion was reinforced by our network-based analyses. We found that the pup call response network of mothers was more refined – sparser, yet stronger, compared to that of naïve virgins, and experienced virgins were positioned in between the two extremes.

By labeling and quantifying OXT projections across the whole brain, we identified a circuit of OXT projection-dense regions that were shared across naïve virgins, experienced virgins, and mothers. Looking specifically at pup call-evoked coordinated activity within this circuit, we found a maternal experience-dependent shift toward lower synchrony during pup call perception. This shift may be facilitated by OXT-mediated inhibition. Pup calls have been shown to activate OXT-expressing neurons in the paraventricular nucleus of the hypothalamus (PVH) of mothers, leading to a release in OXT (Valtcheva et al., 2023). Prior work, which also employed c-Fos-iDISCO+, showed that endogenous OXT release via PVH stimulation leads to less brain-wide c-Fos expression in wild-type mice (Choe et al., 2022). This brain-wide activity suppression may be achieved by OXT binding to OXT receptor-expressing cells, which have a high likelihood of being inhibitory interneurons (K. Li et al., 2016; Marlin et al., 2015; Mitre et al., 2016; Nakajima et al., 2014; Schimmer et al., 2026). Taken together, pup call-evoked OXT release could lead to brain-wide inhibition in OXT receptor-expressing cell populations, which in turn could lead to lower synchrony, both in our defined OXT projection-dense circuit and in the brain at large. However, it should be noted that pup calls are so far only known to induce OXT release in mothers, thus it is unclear if pup calls could have the same OXT-mediated, brain-wide effect in experienced virgins.

Our pup call response network results parallel nicely with recent human neuroimaging research. Recent studies have shown that the transition to motherhood is marked by a reduction in grey matter volume (GMV) that is most drastic around the time of parturition and has been associated with measures of mother-infant attachment. This GMV reduction only partially recovers, even after six years postpartum, and it is primarily only observed, at least to this degree, in gestational mothers and not their partners who are fathers or non-birthing mothers (Hoekzema et al., 2017; Martínez-García et al., 2021; Pawluski et al., 2022; Pritschet et al., 2024; Servin-Barthet et al., 2025). A current hypothesis in the field is that these GMV reductions indicate a period of maturation that supports the transition into motherhood, enhancing the cognitive abilities of new mothers in areas such as social and emotional processing – abilities that play a significant role in adaptive maternal caregiving (Anderson & Rutherford, 2012).

Taken together, the results presented in this study demonstrate that adaptive responses to pups and pup calls are associated with more refined pup call response networks that span the entire brain. These sparser, but stronger pup call response networks may allow maternally experienced mice to perceive and respond to pup calls more efficiently, supporting caregiving behavior.

## Materials and Methods

### Mice

All procedures were approved by the Columbia University Institutional Animal Care and Use Committee. All mice in this project were from a C57BL/6J background, age-matched, and obtained from the Jackson Laboratory. All mice were housed with a 12 hour light/12 hour dark cycle, with food and water available ad libitum with the same food source supplied to all animals.

### Breeding and cohousing

During breeding, a virgin male and female were paired per cage and monitored regularly while they completed two rounds of breeding, two produce experienced mothers. Just before the birth of the second litter, the father was removed. The mothers studied were one to four weeks past the weaning of their second litter, to avoid capturing maternal separation effects in our experiments, and tested to ensure they retrieved pups reliably (at least 70% success in pup retrieval assay). The experienced virgins studied were first tested for baseline pup retrieval while they were naïve to pups, to ensure they were not spontaneous retrievers (no more than 10% success in pup retrieval assay) nor aggressive towards pups. Then, they were gently tail-marked with a permanent marker and placed in the home cage of a second-time mother with a newborn litter of age postnatal day one. If cohoused virgins were tested daily for pup retrieval, then they were tested every 24 hours until 72 hours after the start of cohousing. Otherwise, they were just tested at 72 hours. If the cohoused virgins met the criteria of successful retrieval (at least 70%) after 72 hours, then they were considered experienced. If they were not successful after 72 hours, they were placed back into cohousing and tested every 24 hours going forward until they met criteria. The naïve virgins studied were also tested for pup retrieval to ensure they were not spontaneous retrievers (no more than 10% success) nor aggressive towards pups.

### Pup retrieval assay

Pup retrieval tests were performed in a home cage with normal nesting material under normal white light, and they were video recorded from above. The test mouse was first allowed to habituate to the room for a minimum of two hours. To start a trial, one pup was placed in a corner of the cage, specifically in one of the two corners furthest from the nest (alternated per trial). The test mouse was then allotted two minutes. If within those two minutes, the test mouse retrieved the pup back to the nest, the trial was noted as a success. If those two minutes passed and the pup was still outside the nest, or if any pup-directed aggression was observed, the trial was deemed a failure. Each pup retrieval test consisted of 10 trials. Pups used for retrieval testing were generally of the age postnatal day four to six, except in the case of daily pup retrieval testing in cohoused virgins, in which case test pups could be as young as postnatal day one for baseline testing before cohousing.

### Pup retrieval assay behavioral analysis

Pup retrieval behavior was scored by blinded experimenters using BORIS (Behavioral Observation Research Interactive Software) (Friard & Gamba, 2016), and these observations were used to determine retrieval success per trial and retrieval latency. Average latency to retrieve per trial, latency to first retrieval, and overall retrieval success out of 10 trials was recorded per test mouse. The criteria used for pup retrieval included that naïve animals must exhibit 10% or lower success, otherwise they were considered spontaneous retrievers, and experienced animals must exhibit 70% or higher success. Experienced virgins and mothers were held to the same criteria.

### Pup call recordings

Pup isolation calls were recorded in a sound-isolated chamber. Male and female pups of age postnatal day six were individually placed on a clean Petri dish and placed in the recording chamber. An ultrasonic microphone (Avisoft Bioacoustics Condenser Ultrasound Microphone CM16/CMPA paired with UltraSoundGate 116H and controlled by Avisoft-RECORDER software) was hung from an opening in the center of the chamber ceiling and lowered to be around one inch above the pup. Following one minute of habituation, after which the pups usually began to vocalize, a recording of three to five minutes was acquired.

### Auditory stimulus files

To create pup call stimulus files without audible background noise, all pup call recordings were 15 kHz high-pass filtered. To maintain the natural bout structure of pup calls, bouts from male and female postnatal day six pups were cropped and interlaced into a stimulus file, with an inter-bout-interval of one second. To ensure that the stimuli represented typical pup isolation calls, the stimulus file was run through the VocalMat pipeline (Fonseca et al., 2021), and the distribution of inter-syllable-intervals was compared to that which is reported in existing literature (Schiavo et al., 2020). To create pup call versus white noise stimulus files for the two-choice assay, a pup call stimulus file and a zero to 100 kHz white noise file were matched in average volume and combined into a two-channel stimulus file.

### Auditory two-choice assay

Two-choice assays were performed in a custom-built, opaque white acrylic Y-shaped arena, with transparent acrylic lids and vertically removable partitions separating each chamber. Tests were video recorded from above. Ultrasonic speakers (Avisoft Bioacoustics Vifa Ultrasonic Dynamic Speaker, paired with UltraSoundGate Player 216H) were located outside of the arena, on the other side of a mesh opening in the arena wall. An ultrasonic microphone (Avisoft Bioacoustics CM16/CMPA Condenser Ultrasound Microphone, paired with UltraSoundGate 116H) was placed near one of the two speakers for confirmation of stimulus delivery. The test was split into two days: one habituation day and one test day. The habituation day began with one hour of habituation to the room in the home cage. Then, the test mouse was placed in the center chamber and allotted five minutes to explore the center chamber, followed by 10 minutes to freely explore the entire three-chamber arena after the doors were removed. The test day also began with one hour of habituation to the room in the home cage. Then, the test mouse was placed in the center chamber and allotted two minutes to explore the center chamber, followed by three minutes to freely explore the entire arena. Then, once the mouse was back in the center chamber, the doors were reinserted. Finally, the 10-minute stimulus file containing contrasting auditory stimuli (pup calls and white noise) was started just as the doors were removed once again, and the test mouse was allotted two to 10 minutes to freely explore the full arena during the playback of auditory stimuli. The speaker that played pup calls versus white noise was randomized per test mouse.

### Auditory two-choice assay behavioral analysis

Two-choice assay videos were flipped before analysis such that pup calls were always on the left. Test animals had to spend at least one second in the center chamber to be included in analysis, to ensure both auditory stimuli were sampled. Animal head position was tracked via the software ANY-maze (Stoelting Co.), and interaction zones were delineated around each speaker face. Time spent in each interaction zone, as well as zone entries, were recorded per test mouse. A pup call interaction preference index was calculated per mouse, following the formula:

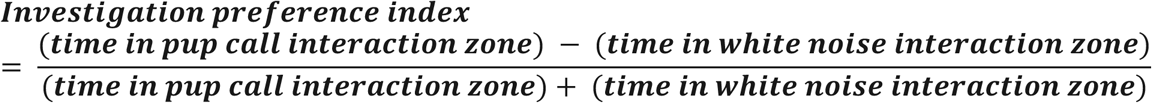

We fit a reduced GLMM for each brain region with this formula:

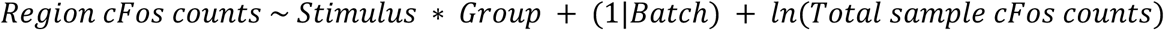

### Sound exposure assay

Sound exposure assays were performed in a clean home cage with a transparent acrylic lid, normal nesting material, and an ultrasonic speaker (Avisoft Bioacoustics Vifa Ultrasonic Dynamic Speaker, paired with UltraSoundGate Player 216H) under red light. Tests were video recorded from above. The test mouse was first allowed to habituate to the room and cage for two hours. To start a trial, the ultrasonic speaker was gently plugged in, and the experimenter began video recording and playing either a 30-minute pup call stimulus file or no stimulus file.

Following 30 minutes of the sound exposure period, the test mouse was left unperturbed until intracardiac perfusion 15 to 30 minutes after the end of sound exposure, or 45 to 60 minutes after the start of sound exposure, following published protocols assessing pup call-evoked c-Fos expression (Fichtel & Ehret, 1999; Geissler et al., 2016).

### Sound exposure assay behavioral analysis

For tracking of the nose, left ear, right ear, midbody, and tailbase, we used DeepLabCut (version 3) (Mathis et al., 2018; Nath et al., 2019). We labeled 20 frames taken from each of the 48 sound exposure videos, with 95% used for training. We used a ResNet-50 based neural network with default parameters for 200 epochs. Following analysis, body part coordinates were then extracted and further processed using custom MATLAB code.

We used MATLAB (version 2022a) to analyze mouse behavior during the sound exposure period. We used tracking data of the nose to represent each mouse’s location in the cage. For each top-view sound exposure assay video, the speaker location was outlined, and the speaker interaction zone was defined as the rectangle with the same width of the speaker and the height of five centimeters. The interaction zone extended five centimeters from the center point of the speaker, to capture timepoints when mice were either on the ground in front of the face of the speaker or climbing the face of the speaker, thus defining a timepoint of speaker interaction.

### Tissue clearing and staining

Mice were deeply anesthetized with isoflurane and fixed with an intracardiac perfusion of 4% paraformaldehyde (PFA) in PBS with heparin. All harvested brains were post-fixed overnight at 4°C in 4% PFA in PBS. Samples were processed with the iDISCO+ tissue clearing and immunolabeling protocol, as described at http://www.idisco.info/ (Renier et al., 2014).

Samples were incubated in primary and secondary antibody solutions for 10 days each. For c-Fos, the antibodies used were rat monoclonal anti-c-Fos (primary; 1:1500; Synaptic Systems #226-017) and goat anti-rat 647 (secondary; 1:1000; ThermoFisher #A-21247). For oxytocin (OXT), the antibodies used were rabbit polyclonal anti-OXT (primary; 1:2000; Phoenix Pharmaceuticals #H-051-01) and goat anti-rabbit plus 555 (secondary; 1:1000; ThermoFisher #A32732).

### Light-sheet microscopy

Imaging was performed with resources available in the Zuckerman Institute’s Cellular Imaging Platform. Cleared and stained brains were imaged on a light-sheet microscope (Ultramicroscope II, Miltenyi Biotech) equipped with a sCMOS camera (Andor Neo) and a 2×/0.5 NA objective lens (MVPLAPO 2×). Brains were kept fully intact and imaged one hemisphere at a time. Samples were oriented sagittally such that the dorsal surface faced the right light-sheet, and only the right light-sheet, at 90% sheet width, was used for imaging. Tiles were set to overlap by 10%. Two acquisitions were completed per session: one for autofluorescence (488 nanometers channel) and one for c-Fos and OXT staining (647 nanometers and 555 nanometers, respectively and simultaneously). For autofluorescence, the following parameters were used: 0.63x zoom (1.26x effective magnification); no dynamic focus; 10 micrometer Z step-size; 50% laser power; 30 millisecond exposure time. For c-Fos and OXT, the following parameters were used: 0.8x zoom (1.6x effective magnification); 20 dynamic focus acquisitions per plane; three micrometer Z step-size; 90% laser power; 50 millisecond exposure time. All images were saved as 3D tiffs, and tiles were stitched together with the software Stitchy (Translucence Biosystems).

### c-Fos+ cell counting

All c-Fos+ cell detection was performed with the open-source software ClearMap 2, available at https://github.com/ClearAnatomics/ClearMap (Renier et al., 2016). The settings used for cell detection were as follows: the background was removed by subtraction of the morphological opened image with a disk shape structure element with main axis of seven pixels in diameter. Cells were detected from peaks and subsequent watershedding, removing background pixels below an intensity cutoff of 700 and selecting cells with sizes between 20 and 500 voxels. These settings corresponded to ClearMap 2 cell detection parameters used in other studies, to facilitate future comparative analyses (Levy et al., 2019; Renier et al., 2016; Topilko et al., 2022). Samples were registered using the average autofluorescence STPR brain registered to the Allen Brain Institute 25 μm map, and its companion annotation map (http://alleninstitute.org/).

### c-Fos expression analysis

We simplified the Allen Brain Atlas down from 1,204 parcellations to 215 regions. This subset of 215 regions still tiled the whole brain, but truncated the granularity to exclude layer-specificity and combine parcellations that were deemed too small for our purposes. Given that the Allen Brain Atlas is symmetrical, c-Fos+ cell counts were summed across bilateral regions for all analyses except those that explicitly tested lateralization.

After using MATLAB (version 2022a) to collate how many c-Fos+ cells were detected per brain region of the 215 region subset, we used R (version 4.4.3) to model the data with generalized linear mixed models (GLMM). We fit a full GLMM for each brain region using the R package glmmTMB (Brooks et al., 2017), with a negative binomial link function and the formula:

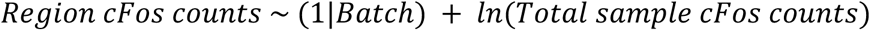

We fit a reduced GLMM for each brain region with this formula:

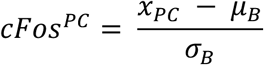

The fixed-effect categorical variables included *Stimulus* (pup calls or baseline) and *Group* (naïve virgin, experienced virgin, or mother), with the asterisk representing both main effects and their interaction. *Batch* (1, 2, 3, or 4) was a random effect for each technical batch (each set of 12 samples that underwent tissue clearing, immunolabelling and light-sheet imaging together). *ln(Total sample cFos counts)* was an offset term for the total number of c-Fos+ cells detected per sample. For each brain region, we fit the full and reduced model, then compared the fit of the 2 models using a likelihood-ratio chi-squared test and adjusted the resulting p-values using the Benjamini-Hochberg method to permit a 10% false discovery rate (FDR) across all brain regions.

For analyses aside from the GLMM, the number of c-Fos+ counts detected per brain region was divided by the total number of c-Fos+ cells detected within the whole sample. This within-sample normalization was done as an attempt to account for potential batch effects in antibody penetration and imaging. To assess which brain regions had a significant batch effect, even after this normalization, we conducted a 1-way ANOVA with Batch being the sole factor.

Following normalization by total sample count, each region’s c-Fos expression was then within-group baseline-standardized. This was achieved with the formula:

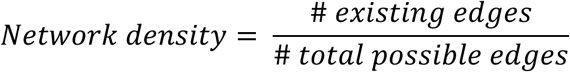

Per brain region in a pup call-exposed sample, *x*_*PC*_ is the observed c-Fos expression; *μ*_*B*_ is the average c-Fos expression in within-group baseline samples; *σ*_*B*_ is the standard deviation of c-Fos expression in within-group baseline samples.

### c-Fos expression correlation matrices

We used MATLAB (version 2022a) to compute, visualize, and compare Pearson correlation coefficients between pairs of brain regions, correlated over samples within the same condition, based on either (1) normalized c-Fos+ cell counts or (2) *cFos*^*PC*^ values. To compare average interregional correlation across conditions, we transformed all interregional Pearson correlation coefficients using a Fisher r-to-z transformation, then conducted either one-way or two-way ANOVAs. To compare similarity between two correlation matrices, we conducted a Mantel test by vectorizing the upper triangle of each correlation matrix, excluding the diagonal, and calculating the Spearman’s correlation coefficient between the vectors.

### c-Fos expression correlation networks

Following the production of brain-wide *cFos*^*PC*^ correlation matrices, we extracted the upper triangle of each matrix, excluding the diagonal, and applied a threshold to all pairwise correlations by their statistical significance, using a FDR of 0.05 and the Benjamini, Krieger, and Yekutieli two-stage step-up method (Benjamini et al., 2006) to correct for multiple comparisons across all pairwise correlations. The pairwise correlations depicted as weighted edges in network graphs all passed this thresholding, and these thresholded networks were then used for network analyses. Treating the resulting networks as unweighted graphs at first, network density was calculated using the formula:

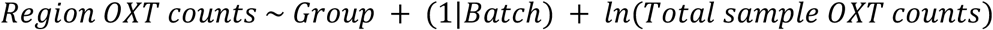

Still treating the networks as unweighted graphs, node degree was calculated per node, defined as the number of edges that node participates in. The average node degree per network was calculated across all 215 nodes. Finally, treating the networks as weighted graphs, we calculated average correlation magnitude by averaging the absolute values of all edges per network. To extract hub regions, each network’s nodes were sorted in descending degree order and the top 10% most-connected nodes out of the 215 total nodes were identified as hubs.

To compare network density, average node degree, and average correlation magnitude between the groups’ pup call response networks, we conducted three permutation tests (n = 1,000 permutations) by shuffling *cFos*^*PC*^ data between (1) naïve virgins and experienced virgins, (2) naïve virgins and mothers, and (3) experienced virgins and mothers. For each permutation, we computed the difference between the two network’s densities, average node degrees, and average correlation magnitudes, and compared the observed differences to their respective null difference distributions to extract one-sided p-values.

### OXT projection mapping

OXT projections were characterized using a custom, consolidated deep learning-based analysis pipeline that automatically preprocesses, segments, and quantifies labeled projections across the whole brain using multiple MATLAB (version R2024b) and Python (version 3.14) scripts. This pipeline was based on TrailMap, developed to segment axonal projections from whole-brain light-sheet images, and DeepTraCE, which aligns TrailMap output to the Allen Brain Atlas and quantifies projection density (Friedmann et al., 2020; Gongwer et al., 2023). We used our pipeline to analyze the left hemisphere of each sample that was imaged left hemisphere first, to avoid any datasets with potential photobleaching. In brief, following brightness normalization and image chunking, we performed OXT projection segmentation with our custom deep learning-based segmentation model. The model was trained with a 3D U-Net architecture (Çiçek et al., 2016) in the open-source PyTorch deep learning framework with brightness-normalized image chunks. The loss function was an equally weighted combination of binary cross-entropy and squared Dice losses (Milletari et al., 2016). To overcome the need for a large amount of initial training data, we used the output of the TrailMap model on their publicly available serotoninergic data to train the foundation of our model. We then refined the model using our OXT training data, acquired by manually tracing entire chunks using the SNT neuroanatomy plugin on ImageJ (version 1.54p) (Arshadi et al., 2021). Following segmentation, image chunks were re-adjoined, and segmented images were binarized using probability threshold of 11 (out of 255), which we found maintained axonal integrity. The binary mask was then skeletonized, connected components were identified and thresholded by a voxel length of 10, and voxels that survived thresholding were deemed to be OXT-labeled. To identify where in the brain these OXT-labeled voxels were located, alignment to the Allen Brain Atlas was achieved via Elastix (Klein et al., 2010), using the same transformation parameters derived from running samples through the ClearMap 2 pipeline (Renier et al., 2016). After acquiring the spatial coordinates and granular region membership of OXT-labeled voxels, we assigned each OXT-labeled voxel to one region in our subset of 215 brain regions.

### OXT expression analysis

After using MATLAB (version 2022a) to collate how many OXT+ voxels were detected per brain region of the 215 region subset, we used R (version 4.4.3) to model the data with generalized linear mixed models (GLMM). We fit a full GLMM for each brain region using the R package glmmTMB (Brooks et al., 2017), with a negative binomial link function and the formula:

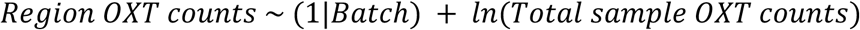

The fixed-effect categorical variable included was *Group* (naïve virgin, experienced virgin, or mother). *Batch* (1, 2, 3, or 4) was a random effect for each technical batch (each set of 12 samples that underwent tissue clearing, immunolabelling and light-sheet imaging together). *ln(Total sample OXT counts)* was an offset term for the total number of OXT+ voxels detected per sample. For each brain region, we fit the full and reduced model, then compared the fit of the 2 models using a likelihood-ratio chi-squared test and adjusted the resulting p-values using the Benjamini-Hochberg method to permit a 10% false discovery rate (FDR) across all brain regions.

For analyses aside from the GLMM, for each brain region, the number of OXT-labeled voxels was divided by the total number of OXT-labeled voxels in the full hemisphere, allowing us to capture the relative proportion of OXT projections present in this region compared to the whole hemisphere. We then calculated relative projection density per region by dividing that number by region volume in cubic millimeters. To define the OXT projection-dense (or sparse) circuit, the relative OXT projection density per region was averaged across all animals, and top (or bottom) 10% most (or least) dense regions were identified.

### Statistical analysis

Results were plotted and tested for statistical significance using MATLAB (versions 2022a and 2024b), R (version 4.4.3), and GraphPad Prism (version 10). Statistical tests are described in earlier Methods sections and figure legends.

## Supporting information

Behavior Supplemental Videos

c-Fos Supplemental Videos

Oxytocin Supplemental Videos

## Acknowledgements

We would like to thank the Marlin lab for their insightful feedback and discussions. We are especially grateful to Drs. V. Andreu and N. Mimouni, as well as J. Osty for their support. We also thank L. Hammond and H. Ibarra Avila of the Zuckerman Institute Cellular Imaging Platform, T. Tabachnik and M. Sullivan of the Zuckerman Institute Advanced Instrumentation Platform, as well as Dr. M. Farinella. We also thank Drs. A. Bendesky, I. Abdus-Saboor, R. Froemke, and C. Zuker for their feedback on this manuscript. Research reported in this publication was supported by the Eunice Kennedy Shriver National Institute of Child Health & Human Development of the National Institutes of Health under award number F31HD114466 (BRM), the Howard Hughes Medical Institute (BJM), the UNCF E.E. Just Fellowship CU20-1071 (BJM), the BBRF NARSAD Young Investigator Grant 30380 (BJM), The Whitehall Foundation (BJM), and the National Institutes of Health under award number 1S10OD023587-01 (Zuckerman Institute Cellular Imaging Platform). The content is solely the responsibility of the authors and does not necessarily represent the official views of the National Institutes of Health.

## Author Contributions

Conceptualization: BRM, DKDF, KS, GA, BJM

Methodology: BRM, DKDF, KS, GA, BJM

Investigation: BRM, DKDF, KS, BJM

Visualization: BRM, DKDF, KS, BJM

Funding Acquisition: BRM, BJM

Project Administration: BRM, BJM

Supervision: BJM

Writing – original draft: BRM

Writing – review & editing: BRM, DKDF, BJM

## Extended Figures

**Extended Figure 1:**
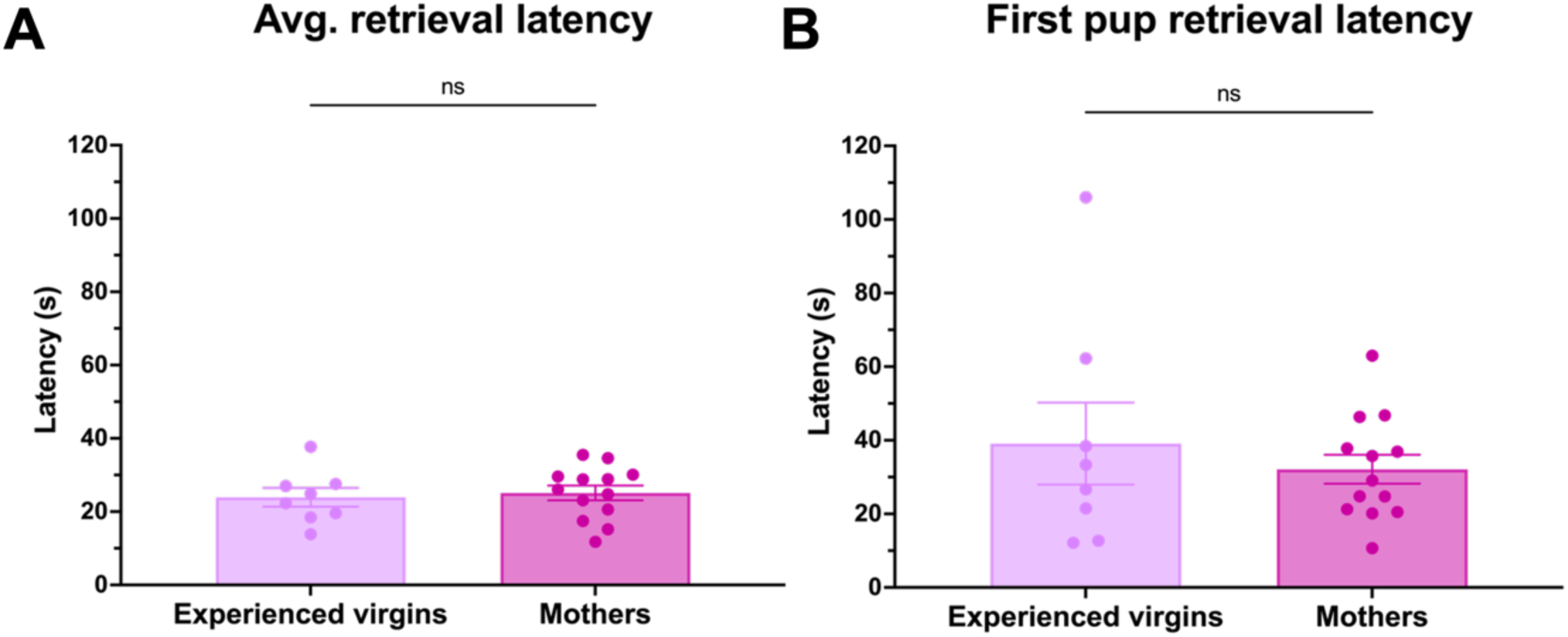
Mothers and experienced virgins exhibit indistinguishable pup retrieval latency. (A) Average pup retrieval latency per mouse. Unpaired t-test revealed no group effect (p = 0.7181; n = 8 experienced virgins, 13 mothers). Error bars denote mean ± SEM. (B) First trial pup retrieval latency per mouse. Unpaired t-test revealed no group effect (p = 0.4914; n = 8 experienced virgins, 13 mothers). Error bars denote mean ± SEM.

**Extended Figure 2:**
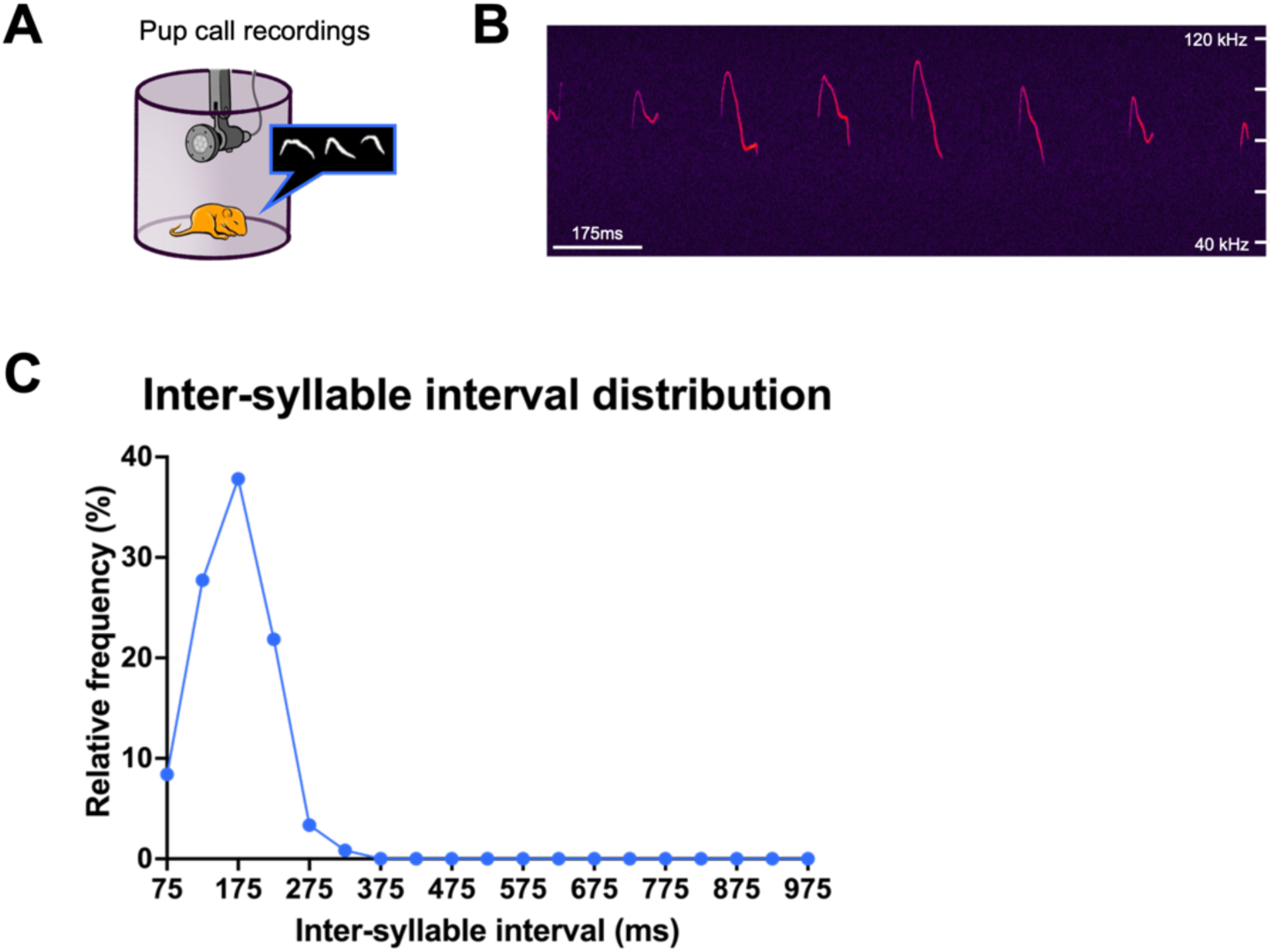
Pup call isolation and analysis. (A) Pup call recording protocol, see Methods for details. (B) Representative excerpt from pup call stimulus file, see Methods for details. (C) Frequency distribution of inter-syllable intervals in pup call stimulus file (n = 188). Dots indicate bin center (± 25 ms).

**Extended Figure 3:**
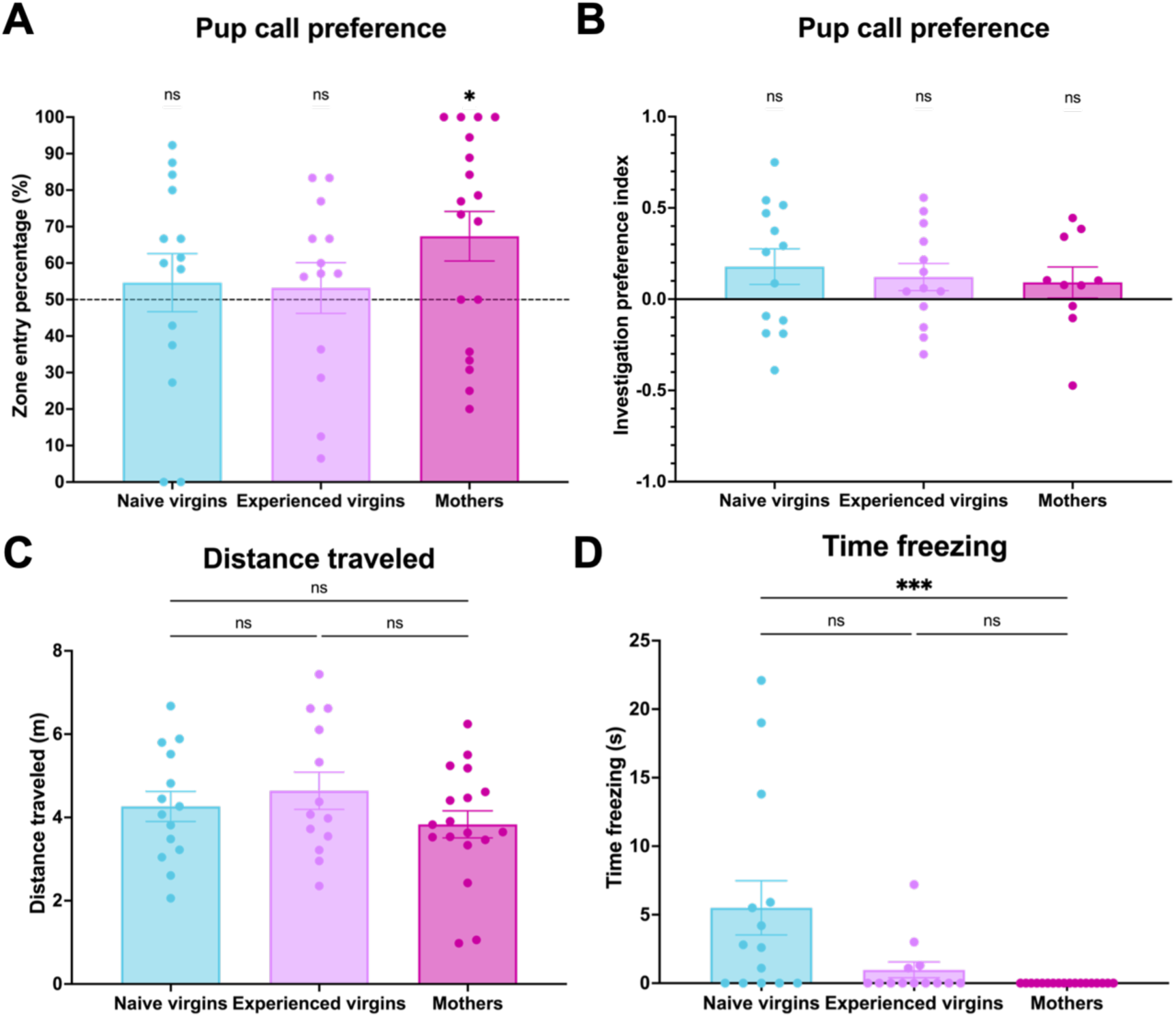
Additional auditory 2-choice assay behavioral measures. (A) Pup call zone entry percentage per mouse during first 2 minutes of trial. 1-sample Wilcoxon tests (versus 50%) revealed only mothers prefer pup calls (naïve virgins: p = 0.4915; experienced virgins: p = 0.6257; mothers: p = 0.0127; n = 14 naïve virgins, 13 experienced virgins, 18 mothers). Kruskal-Wallis revealed no significant group effect (p = 0.2674; n = 14 naïve virgins, 13 experienced virgins, 18 mothers). Dashed line represents no preference (50%). Error bars denote mean ± SEM. Comparisons on graph represent 1-sample Wilcoxon tests. * p < 0.05. (B) Investigation preference index per mouse during full 10 minutes of trial. 1-sample t-tests revealed no preferences (naïve virgins: p = 0.0919; experienced virgins: p = 0.1287; mothers: p = 0.3083; n = 13 naïve virgins, 13 experienced virgins, 10 mothers). 1-way ANOVA revealed no significant group effect (p = 0.7794; n = 13 naïve virgins, 13 experienced virgins, 10 mothers). Error bars denote mean ± SEM. Comparisons on graph represent 1-sample t-tests. (C) Distance traveled during first 2 minutes of trial. 1-way ANOVA revealed no significant group effect (p = 0.3101; n = 14 naïve virgins, 13 experienced virgins, 18 mothers). Error bars denote mean ± SEM. (D) Time spent freezing (threshold: at least 1 second) during first 2 minutes of trial. Kruskal-Wallis test revealed significant group effect (overall: p = 0.0003; naïve virgins vs. experienced virgins: p = 0.0977; naïve virgins vs. mothers: p = 0.0002; experienced virgins vs. mothers: p = 0.2915; n = 14 naïve virgins, 13 experienced virgins, 18 mothers). Error bars denote mean ± SEM. *** p < 0.001.

**Extended Figure 4:**
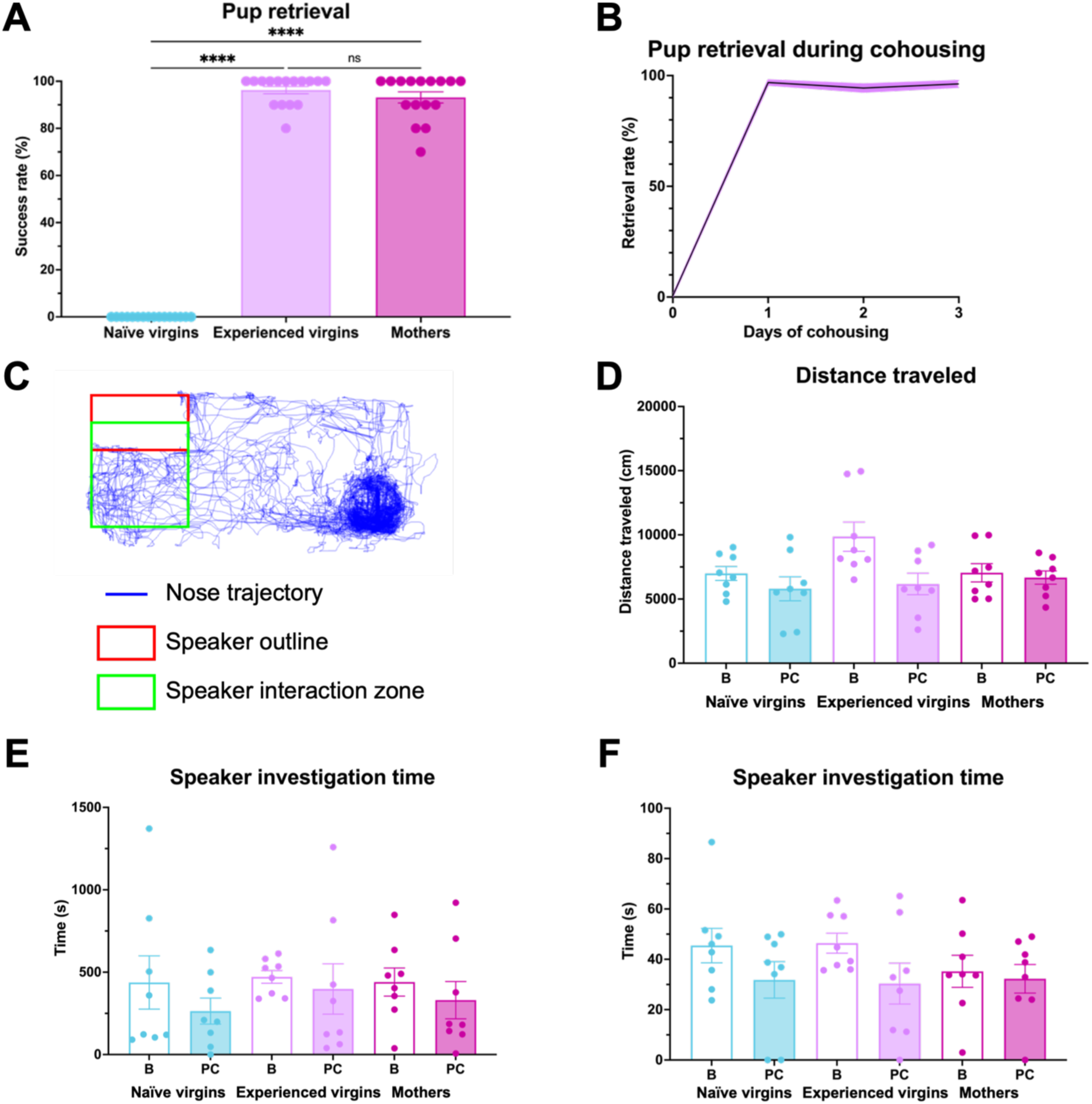
Behavior of mice included in whole-brain imaging data. (A) Pup retrieval success per mouse. Kruskal-Wallis test revealed significant group effect (p < 0.0001). Comparisons on graph represent Tukey’s post-hoc tests (naïve virgins vs. experienced virgins: p < 0.0001; naïve virgins vs. mothers: p < 0.0001; experienced virgins vs. mothers: p > 0.9999; n = 16 per group). Error bars denote mean ± SEM. **** p < 0.0001, ** p < 0.01. (B) Pup retrieval success of virgin females during cohousing with mother and litter (n = 16 experienced virgins). Purple shading denotes mean ± SEM. (C) Representative nose trajectory of a mouse during 30-minute sound exposure period, with speaker location (red) and interaction zone (green) outlined, see Methods for details. (D) Distance traveled per mouse during 30-minute sound exposure period (B = baseline, PC = pup calls). 2-way ANOVA revealed significant stimulus effect (group: p = 0.1330; stimulus: p = 0.0115; interaction: p = 0.1160; n = 8 per condition). Error bars denote mean ± SEM. (E) Speaker investigation time per mouse during 30-minute sound exposure period. 2-way ANOVA revealed no significant effects (group: p = 0.7594; stimulus: p = 0.2065; interaction: p = 0.9052; n = 8 per condition). Error bars denote mean ± SEM. (F) Speaker investigation time per mouse during first 2 minutes of sound exposure period. 2-way ANOVA revealed significant stimulus effect (group: p = 0.7000; stimulus: p = 0.0464; interaction: p = 0.5663; n = 8 per condition). Error bars denote mean ± SEM.

**Extended Figure 5:**
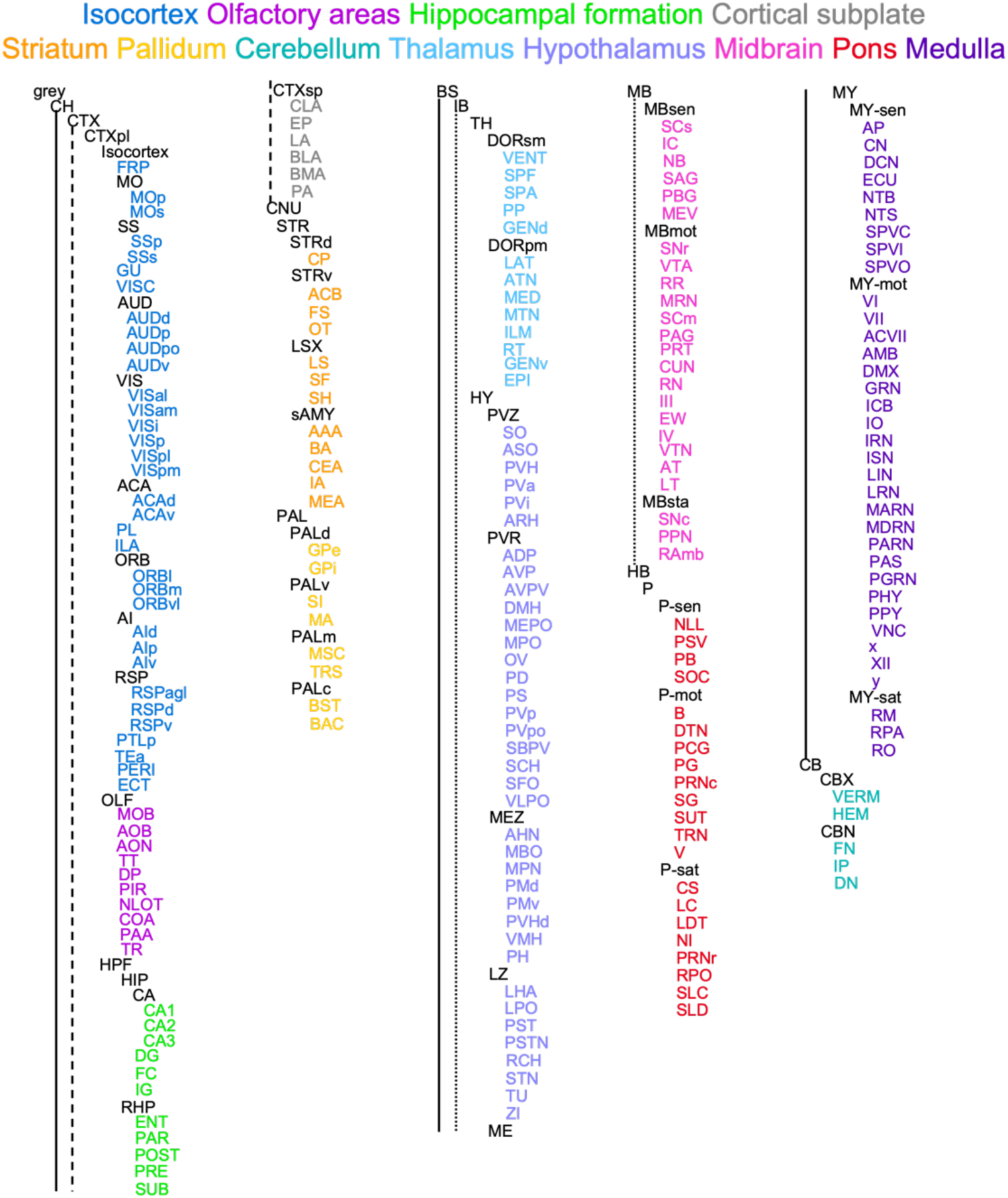
Partially collapsed Allen Brain Atlas. Rather than keeping the full 1,204 parcellations of the Allen Brain Atlas, the atlas was simplified down to 215 parcellations (see Table 1 for abbreviations). This was achieved by eliminating nonsense regions (i.e., ventricles, fibers, unlabeled) and limiting the level of specificity (i.e., layers). The 215 retained parcellations are represented as leaves in this tree, color-coded by broader brain area membership as indicated by the key at the top. Regions written in black text indicate that they were not included in the final 215.

**Extended Figure 6:**
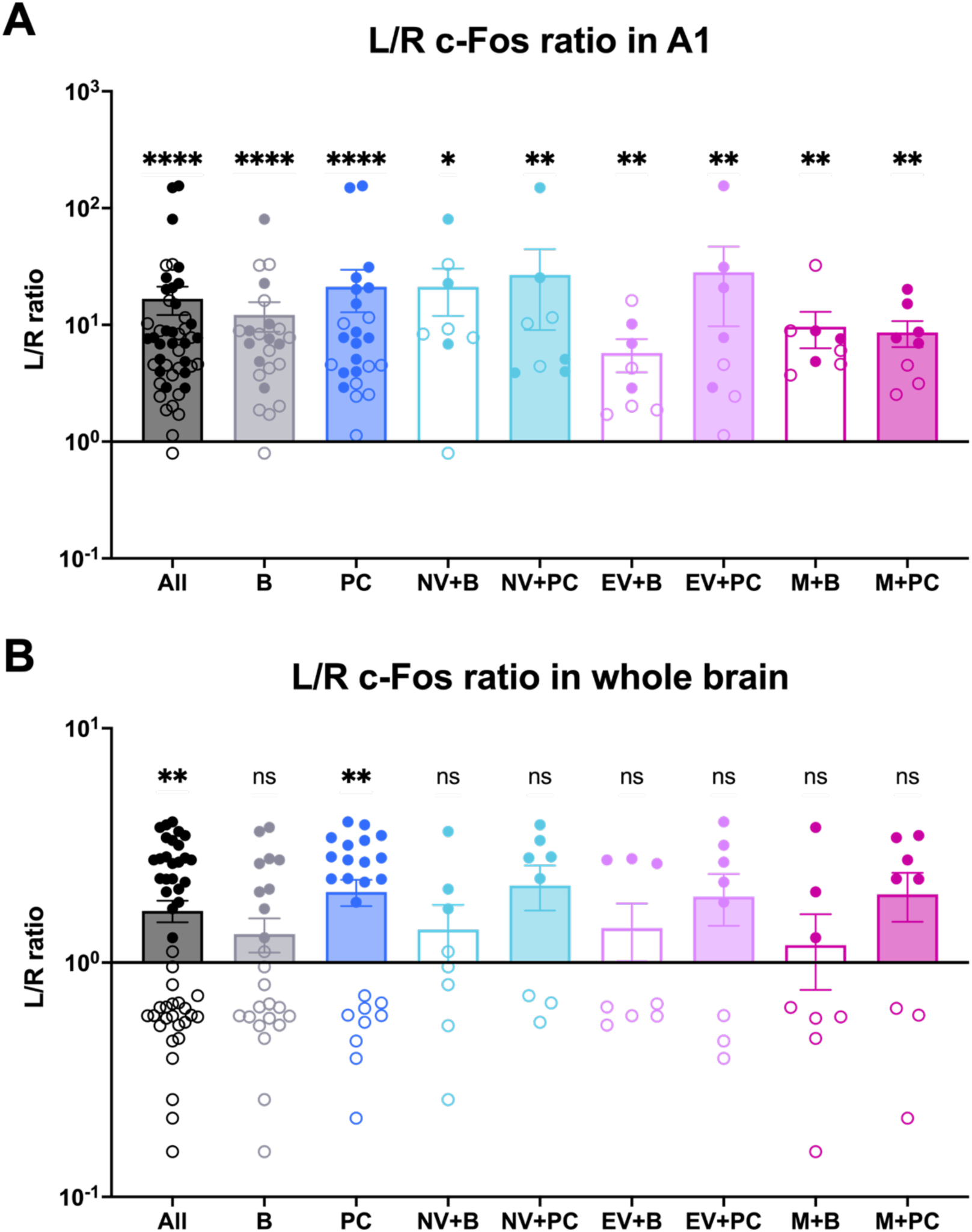
Lateralization in c-Fos dataset. Ratio (left/right) of number of c-Fos+ cells detected in A1 (A) or whole brain (B) of each sample, split by condition (All = all groups, all stimuli; B = all groups, baseline only; PC = all groups, pup calls only; NV+B = naïve virgins only, baseline only; NV+PC = naïve virgins only, pup calls only; EV+B = experienced virgins only, baseline only; EV+PC = experienced virgins only, pup calls only; M+B = mothers only, baseline only; M+PC = mothers only, pup calls only). Unfilled dots represent samples for which right hemisphere was imaged first. Error bars denote mean ± SEM. (A) Wilcoxon tests (versus 1) revealed significant lateralization in A1, regardless of condition (All: p < 0.0001, n = 48; B: p < 0.0001, n = 24; PC: p < 0.0001, n = 24; NV+B: p = 0.0156, n = 8; NV+PC: p = 0.0078, n = 8; EV+B: p = 0.0078, n = 8; EV+PC: p = 0.0078, n = 8; M+B: p = 0.0078, n = 8; M+PC: p = 0.0078, n = 8). **** p < 0.0001, ** p < 0.01, * p < 0.05. (B) Wilcoxon tests (versus 1) revealed minimal lateralization in whole brain, regardless of condition (All: p = 0.0059, n = 48; B: p = 0.6231, n = 24; PC: p = 0.0018, n = 24; NV+B: p = 0.6406, n = 8; NV+PC: p = 0.1094, n = 8; EV+B: p = 0.7422, n = 8; EV+PC: p = 0.1094, n = 8; M+B: p = 0.8438, n = 8; M+PC: p = 0.1094, n = 8). ** p < 0.01.

**Extended Figure 7:**
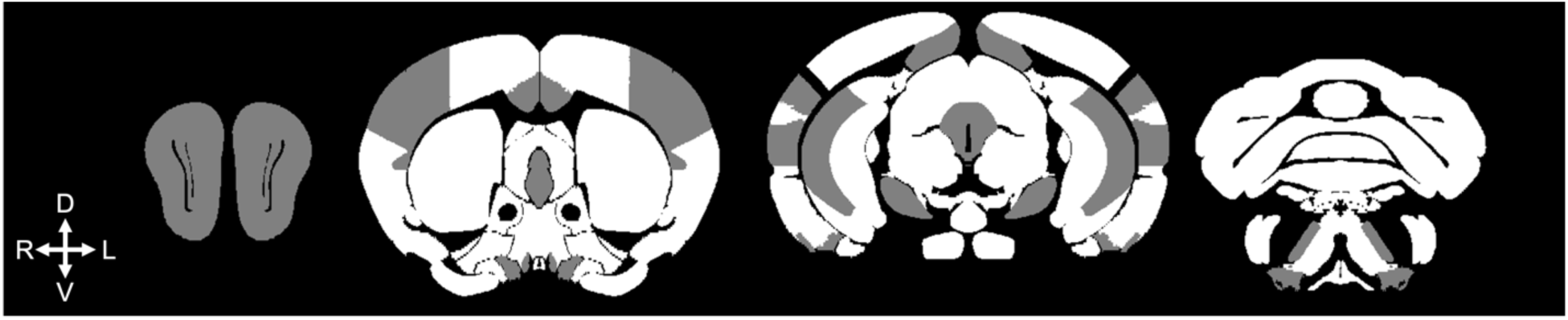
Randomly distributed batch effect. 4 representative coronal slices representing brain regions with (white) and without (gray) batch effect, as determined by 1-way ANOVA that used batch to explain normalized c-Fos per brain region (see Methods). Note the seemingly random spatial distribution that did not appear to have any directionality or clustering (D = dorsal, V = ventral, R = right, L = left).

**Extended Figure 8:**
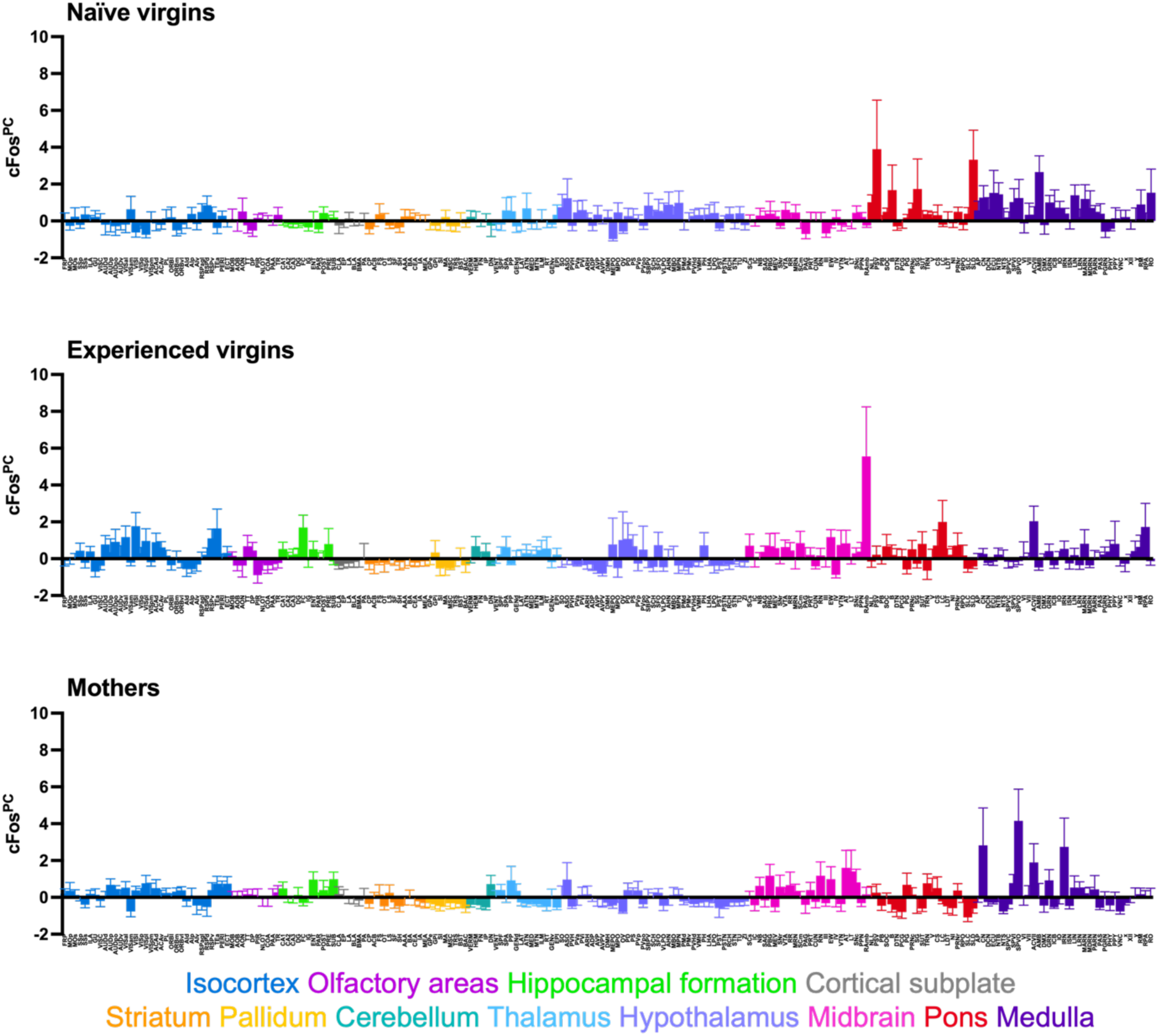
Baseline-standardized c-Fos expression. ***cFos***^***PC***^per brain region (215 regions; see Table 1 for abbreviations), averaged per group (n = 8). See Methods for details. Error bars denote mean ± SEM.

**Extended Figure 9:**
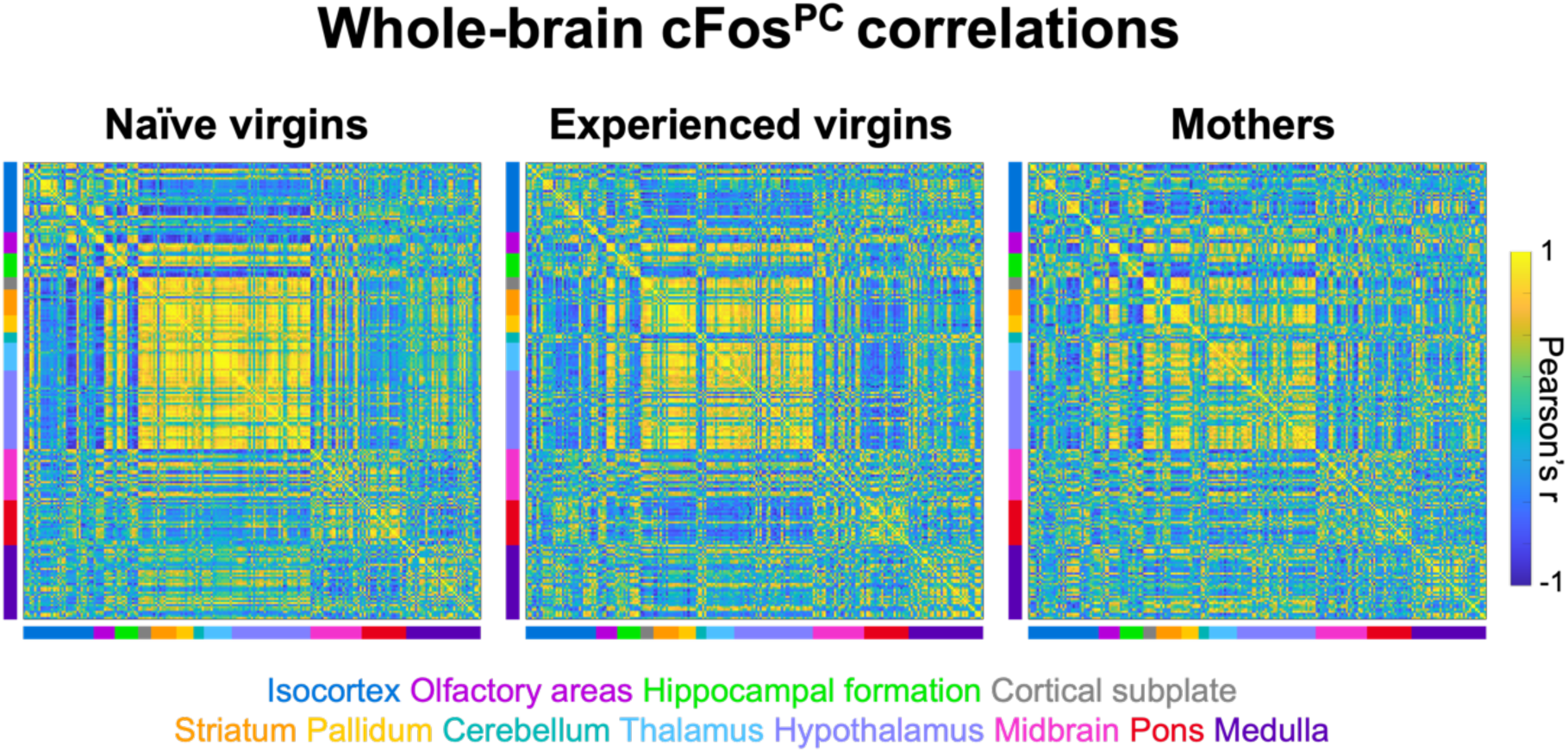
Brain-wide coordinated activation upon pup call exposure differs between groups. Pearson correlation matrices indicating interregional correlations, based on ***cFos***^***PC***^in pup call-exposed samples across groups (n = 8 x 215 regions). Color-blocked bars on axes correspond to broader brain areas according to the Allen Brain Atlas, see text below matrices for color coding. Matrix pixel colors represent Pearson correlation coefficients, see scale to the right.

**Extended Figure 10:**
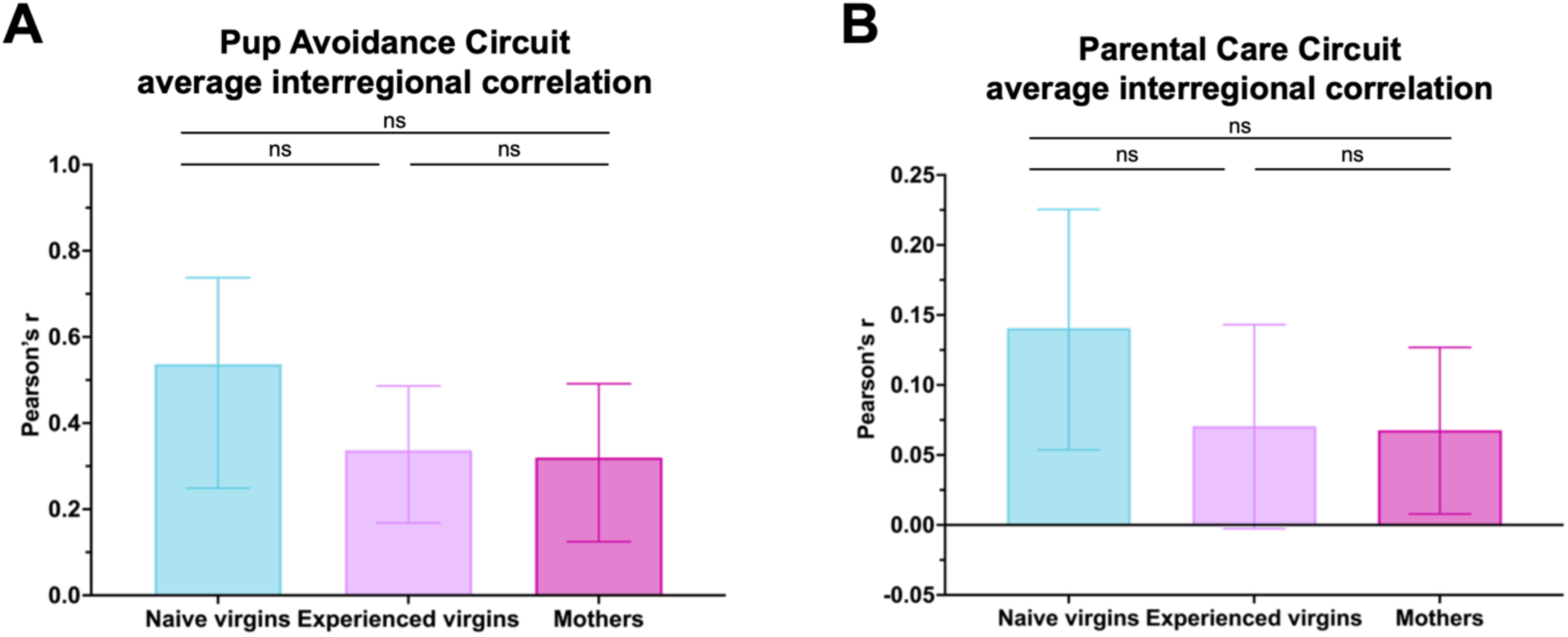
Pup Avoidance and Parental Care Circuits reveal similar synchrony between groups. (A) Mean interregional correlation in Pup Avoidance Circuit per group. Kruskal-Wallis test following Fisher r-to-z transformation revealed no significant group effect (p = 0.1752; n = 8 x 9 regions). Error bars denote mean ± 95% CI. (B) Mean interregional correlation in Parental Care Circuit per group. Kruskal-Wallis test following Fisher r-to-z transformation revealed no significant group effect (p = 0.6972; n = 8 x 27 regions). Error bars denote mean ± 95% CI.

**Extended Figure 11:**
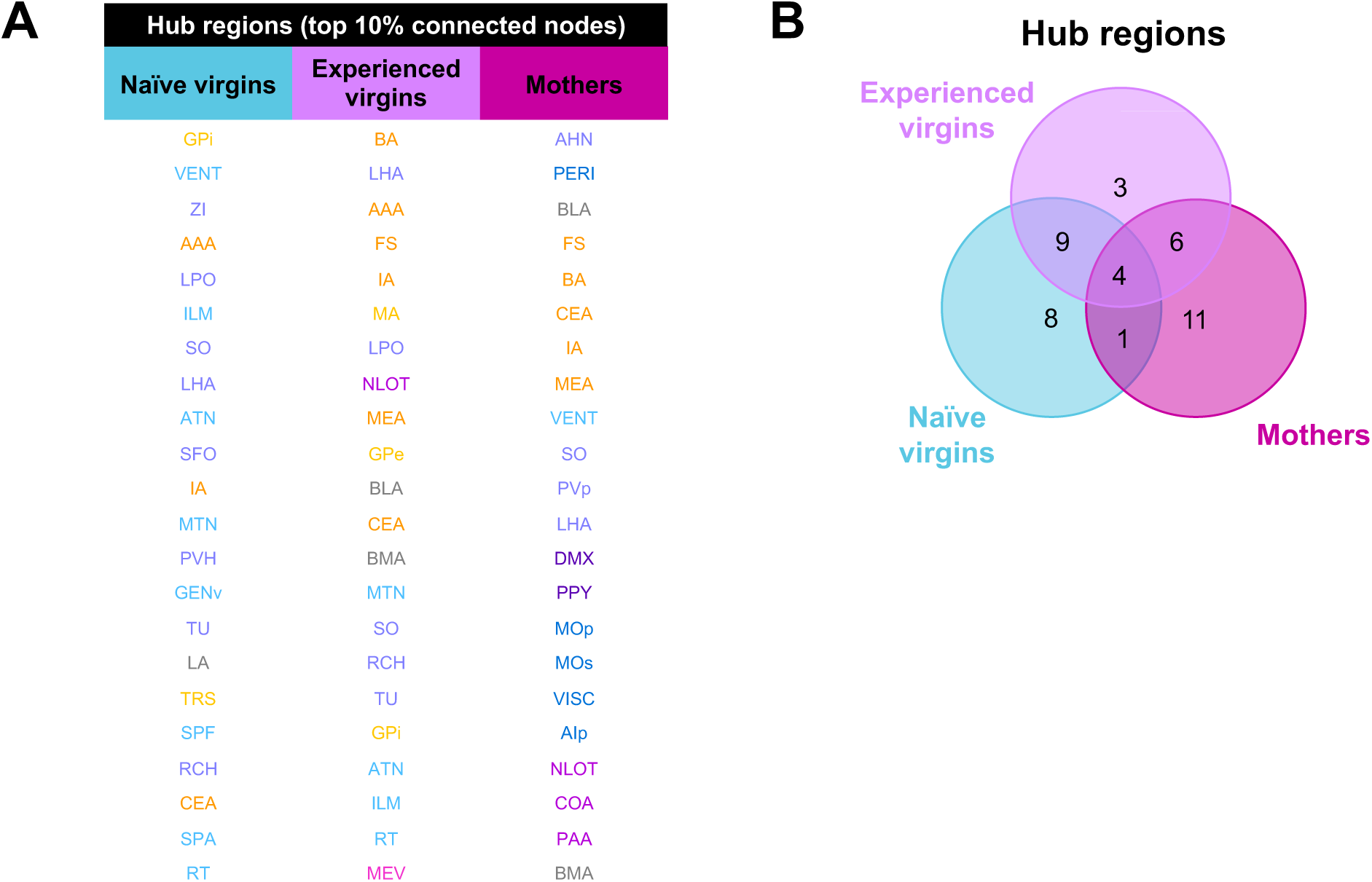
Pup call response hub regions across groups. (A) Pup call response hub regions (top 10% most connected nodes, as defined by node degree; see Table 4) per group. (B) Venn diagram depicting number of unique and overlapping hub regions.

## Notes

https://github.com/BJMarlinLab

### Competing Interest Statement

The authors have declared no competing interest.

